# Development of ultrafast camera-based imaging of single fluorescent molecules and live-cell PALM

**DOI:** 10.1101/2021.10.26.465864

**Authors:** Takahiro K. Fujiwara, Shinji Takeuchi, Ziya Kalay, Yosuke Nagai, Taka A. Tsunoyama, Thomas Kalkbrenner, Kokoro Iwasawa, Ken P. Ritchie, Kenichi G.N. Suzuki, Akihiro Kusumi

**Author notes:** To whom correspondence should be addressed: Akihiro Kusumi, Fx: 011-81-98-966-1062.

## Abstract

The spatial resolution of fluorescence microscopy has recently been greatly improved. However, its temporal resolution has not been improved much, despite its importance for examining living cells. Here, by developing an ultrafast camera system, we achieved the world’s highest time resolutions for single fluorescent-molecule imaging of 33 (100) µs (multiple single molecules simultaneously) with a single-molecule localization precision of 34 (20) nm for Cy3 (best dye found), and for PALM data acquisition of a view-field of 640×640 pixels at 1 kHz with a single-molecule localization precision of 29 nm for mEos3.2. Both are considered the ultimate rates with available probes. This camera system (1) successfully detected fast hop diffusion of membrane molecules in the plasma membrane, detectable previously only by using less preferable 40-nm gold probes and bright-field microscopy, and (2) enabled PALM imaging of the entire live cell, while revealing meso-scale dynamics and structures, caveolae and paxillin islands in the focal adhesion, proving its usefulness for cell biology research.

**Summary:** An ultrafast camera developed by Fujiwara et al. allows single fluorescent-molecule imaging every 33 μs with a localization precision of 34 nm (every 100 μs; 20 nm), and enables ultrafast PALM imaging of whole live cells.

## Introduction

Extensive attention has recently focused on improving the spatial resolution of fluorescence microscopy, leading to the development of various types of super-resolution methods (Shcherbakova et al., 2014; Liu et al., 2015; Nicovich et al., 2017; Sahl et al., 2017; von Diezmann et al., 2017; Baddeley and Bewersdorf, 2018; Sigal et al., 2018; Sezgin et al., 2019; Lelek et al., 2021). Meanwhile, efforts toward enhancing the temporal resolution of fluorescence microscopy, particularly in single fluorescent-molecule imaging and single-molecule localization microscopy (SMLM), have been quite limited (Wieser et al., 2007; Jones et al., 2011; Huang et al., 2013; Hiramoto-Yamaki et al., 2014; Kinoshita et al., 2017). However, for observing living cells, and particularly for investigating the dynamic movements and interactions of molecules at the single molecule level (Baboolal et al., 2016), improving the temporal resolution is critically important (Koyama-Honda et al., 2020). This was exemplified by our detection of hop diffusion of membrane molecules in the plasma membrane (PM) by improving the time resolution of single-*particle* tracking down to 25 µs (Fujiwara et al., 2002, 2016; Murase et al., 2004). However, this method required the use of large (40-nm diameter) gold particles as probes. Since fluorescent probes are much smaller (<1 nm) and used far more broadly than gold probes, in the present study, we tried to improve the temporal resolution of imaging single fluorescent molecules (many molecules at the same time) to levels similar to those of gold-particle tracking.

For tracking just one molecule at a time, a method called MINFLUX has recently been developed. For molecules adsorbed on glass, it provides 2.4-nm and <20 nm localization precisions with 400- and 117-µs temporal resolutions, respectively; in live *E. coli*, it provides 48-nm precision with 125-µs time resolution (Balzarotti et al., 2017; Eilers et al., 2018; Schmidt et al., 2021). However, MINFLUX does not allow observations of more than one molecule at a time, and thus it cannot be used to study molecular interactions and simultaneous events occurring in various parts of the cell. Consequently, its applications would be quite limited.

Here, we developed a camera-based method that allows the observations of several tens to hundreds of molecules simultaneously in a field of view (like in a cell). This method would be useful for investigating the interactions and assemblies of molecules in live cells and simultaneously studying the different behaviors of molecules in various parts of the cell. This ultrafast camera system enables the simultaneous tracking of many single fluorescent molecules in the view-field at 10 kHz or every 100 µs, with a localization precision down to 20 nm with a frame size of 14 x 14 µm^2^ (256 x 256 pixels), and in the case of 30 kHz or every 33 µs, to a localization precision of 34 nm with a frame size of 7.1 x 6.2 µm^2^ (128 x 112 pixels) (for previous results using other cameras, see the summary in **Table 1**).

**Table 1.**
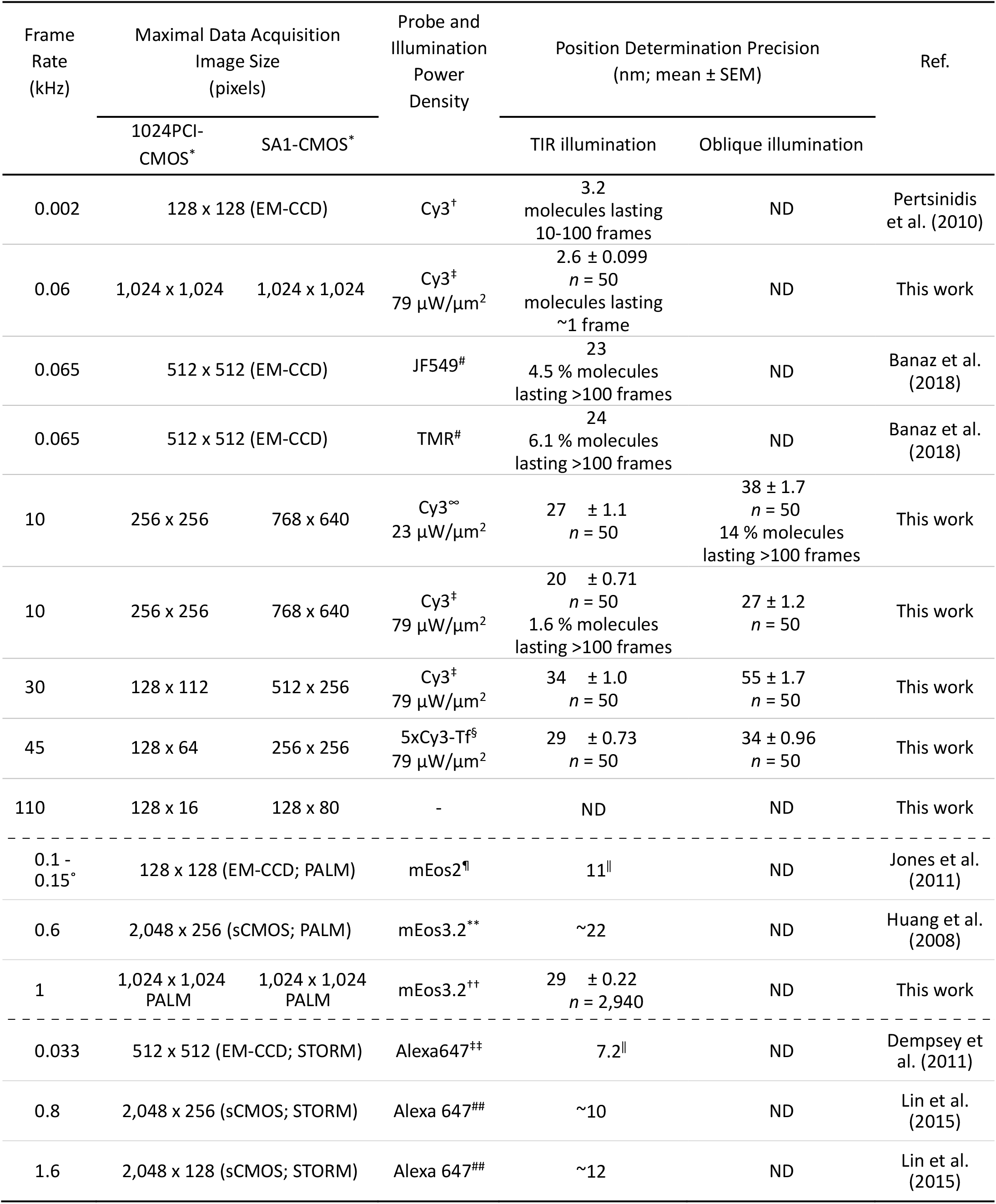
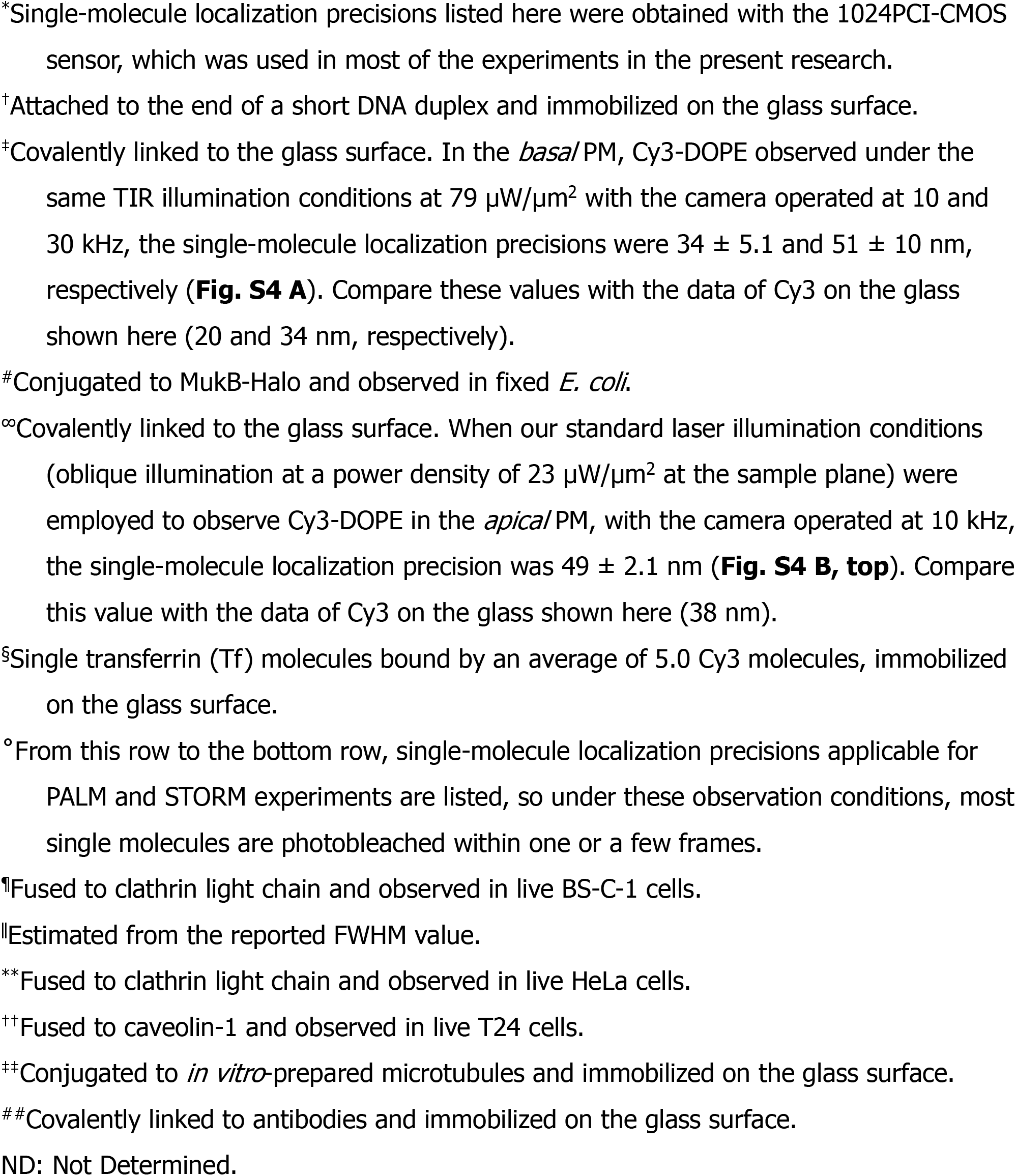
Summary of the camera frame rate, maximal data acquisition image size (pixels), position determination precision, and number of observable frames for various fluorescent probes for the developed camera system and other cameras.

| Frame Rate (kHz) | Maximal Data Acquisition Image Size (pixels) | | Probe and Illumination Power Density | Position Determination Precision (nm; mean ± SEM) | | Ref. |
| --- | --- | --- | --- | --- | --- | --- |
|  | 1024PCI-CMOS* | SA1-CMOS* |  | TIR illumination | Oblique illumination |  |
| 0.002 | 128 x 128 (EM-CCD) |  | Cy3<sup>†</sup> | 3.2 molecules lasting 10-100 frames | ND | Pertsinidis et al. (2010) |
| 0.06 | 1,024 x 1,024 | 1,024 x 1,024 | Cy3<sup>‡</sup><br>79 $\mu\text{W}/\mu\text{m}^2$ | 2.6 ± 0.099<br>$n = 50$<br>molecules lasting ~1 frame | ND | This work |
| 0.065 | 512 x 512 (EM-CCD) |  | JF549<sup>#</sup> | 23<br>4.5 % molecules lasting >100 frames | ND | Banaz et al. (2018) |
| 0.065 | 512 x 512 (EM-CCD) |  | TMR<sup>#</sup> | 24<br>6.1 % molecules lasting >100 frames | ND | Banaz et al. (2018) |
| 10 | 256 x 256 | 768 x 640 | Cy3<sup>∞</sup><br>23 $\mu\text{W}/\mu\text{m}^2$ | 27 ± 1.1<br>$n = 50$ | 38 ± 1.7<br>$n = 50$<br>14 % molecules lasting >100 frames | This work |
| 10 | 256 x 256 | 768 x 640 | Cy3<sup>‡</sup><br>79 $\mu\text{W}/\mu\text{m}^2$ | 20 ± 0.71<br>$n = 50$<br>1.6 % molecules lasting >100 frames | 27 ± 1.2<br>$n = 50$ | This work |
| 30 | 128 x 112 | 512 x 256 | Cy3<sup>‡</sup><br>79 $\mu\text{W}/\mu\text{m}^2$ | 34 ± 1.0<br>$n = 50$ | 55 ± 1.7<br>$n = 50$ | This work |
| 45 | 128 x 64 | 256 x 256 | 5xCy3-Tf<sup>§</sup><br>79 $\mu\text{W}/\mu\text{m}^2$ | 29 ± 0.73<br>$n = 50$ | 34 ± 0.96<br>$n = 50$ | This work |
| 110 | 128 x 16 | 128 x 80 | - | ND | ND | This work |
| 0.1 - 0.15° | 128 x 128 (EM-CCD; PALM) |  | mEos2<sup>¶</sup> | 11<sup>¶</sup> | ND | Jones et al. (2011) |
| 0.6 | 2,048 x 256 (sCMOS; PALM) |  | mEos3.2<sup>**</sup> | ~22 | ND | Huang et al. (2008) |
| 1 | 1,024 x 1,024 PALM | 1,024 x 1,024 PALM | mEos3.2<sup>++</sup> | 29 ± 0.22<br>$n = 2,940$ | ND | This work |
| 0.033 | 512 x 512 (EM-CCD; STORM) |  | Alexa647<sup>††</sup> | 7.2<sup>¶</sup> | ND | Dempsey et al. (2011) |
| 0.8 | 2,048 x 256 (sCMOS; STORM) |  | Alexa 647<sup>###</sup> | ~10 | ND | Lin et al. (2015) |
| 1.6 | 2,048 x 128 (sCMOS; STORM) |  | Alexa 647<sup>###</sup> | ~12 | ND | Lin et al. (2015) |

These frame rates are 333x and 1,000x faster than video rate, respectively, and comparable to the rate at which MINFLUX tracks just one molecule. For camera-based technology, this rate is likely to represent the ultimate attainable with the currently available fluorescent dyes: the camera system itself could be operated faster, and thus the instrument is no longer the limitation for faster observations. With the future development of fluorescent dyes with faster fluorescent photon emission, we should be able to observe single molecules at 45 kHz (every 22 µs; with a frame size of 7 x 3.5 µm^2^) or even faster rates.

We examined the applicability of this camera system to live cells by testing whether it can detect the hop diffusions of (Cy3-labeled) single molecules, a phospholipid and a transmembrane protein, transferrin receptor (TfR), which were previously detectable only by tracking 40-nm gold particles bound to these molecules in the *apical* (dorsal) plasma membrane (PM; since 40-nm-diameter gold particles cannot enter the space between the basal PM and the coverslip, the hop diffusion in the basal PM could not be investigated previously; this will be described in the companion paper [Fujiwara et al., 2021]). The single fluorescent-molecule imaging system based on the ultrafast camera developed here successfully reproduced the gold particle data, without affecting cell viability. Furthermore, coupled with improved analysis methods, it allowed the determination of the residency time of each single molecule in each membrane compartment.

At the same time, the developed camera system has greatly improved the time resolution and the obtainable image size of SMLM (in the present report, we focus on PALM, but it would also improve the time resolutions of fPALM, iPALM, STORM, and dSTORM). With the developed ultrafast camera system, the PALM instrument should be able to operate at 10 kHz or faster, but just like the case of ultrafast single-molecule imaging, we found that one of the best photoconvertible fluorescent proteins available for PALM today, mEos3.2 (Zhang et al., 2012), only allowed us to operate the instrument at 1 kHz. However, even under these conditions, the PALM data acquisition of a view-field of 640 x 640-pixels (35.3 x 35.3 µm^2^) for 10,000 (or 300) frames was shortened to 10 (0.33) s, with a single-molecule localization precision for mEos3.2 of 29 ± 0.22 (SEM) nm. The spatial resolution of the PALM image was ∼75 nm, as determined by a parameter-free decorrelation analysis (Descloux et al., 2019), which is somewhat worse than the conventional live-cell PALM images (generally ∼60 nm; Shroff et al., 2008; Huang et al., 2013) due to the ∼2x lower quantum efficiency of the photosensor of this camera system, as compared with those of generally employed cameras (for comparisons with previously used cameras, see **Table 1**). Namely, this camera system now allows observations of almost the entire live cell at ∼75-nm resolutions every 0.33 – 10 s. With the future development of photoconvertible molecules with faster photon emission and photobleaching, we would be able to obtain PALM images >10 times faster; i.e., at video rate or faster.

In the companion paper, we describe further applications of this ultrafast camera system (Fujiwara et al., 2021). We report the compartmentalization of the basal PM as well as the apical PM, the hop diffusion of phospholipids and transmembrane proteins, TfR and EGF receptor (EGFR), in the basal PM, and the compartmentalization of the focal adhesion region for the diffusion of membrane molecules.

## Results

### Basic design of the ultrafast camera system

Our camera system consists of a microchannel-plate image intensifier (Fig. 1 A-a), a high-speed complementary metal-oxide semiconductor (CMOS) sensor (Fig. 1 A-b), and an optical-fiber bundle, which couples the phosphor screen of the image intensifier to the CMOS sensor chip (Fig. 1 A-c; **Materials and methods**). The image intensifier (Hamamatsu, V8070U-74) comprises a third-generation GaAsP photocathode with a quantum efficiency of 40% at 570 nm (Fig. 1 A-a-α), a three-stage microchannel plate allowing maximal gain over 10^6^, in which a non-linear noise increase was virtually undetectable (Fig. 1 A-a-β), and a P46 phosphor screen with a decay time of approximately 0.3 µs from 90% to 10% (Fig. 1 A-a-γ). A CMOS sensor (the sensor developed for a Photron 1024PCI camera), with a global shutter exposure was used (**Table 1**).

**Figure 1.**
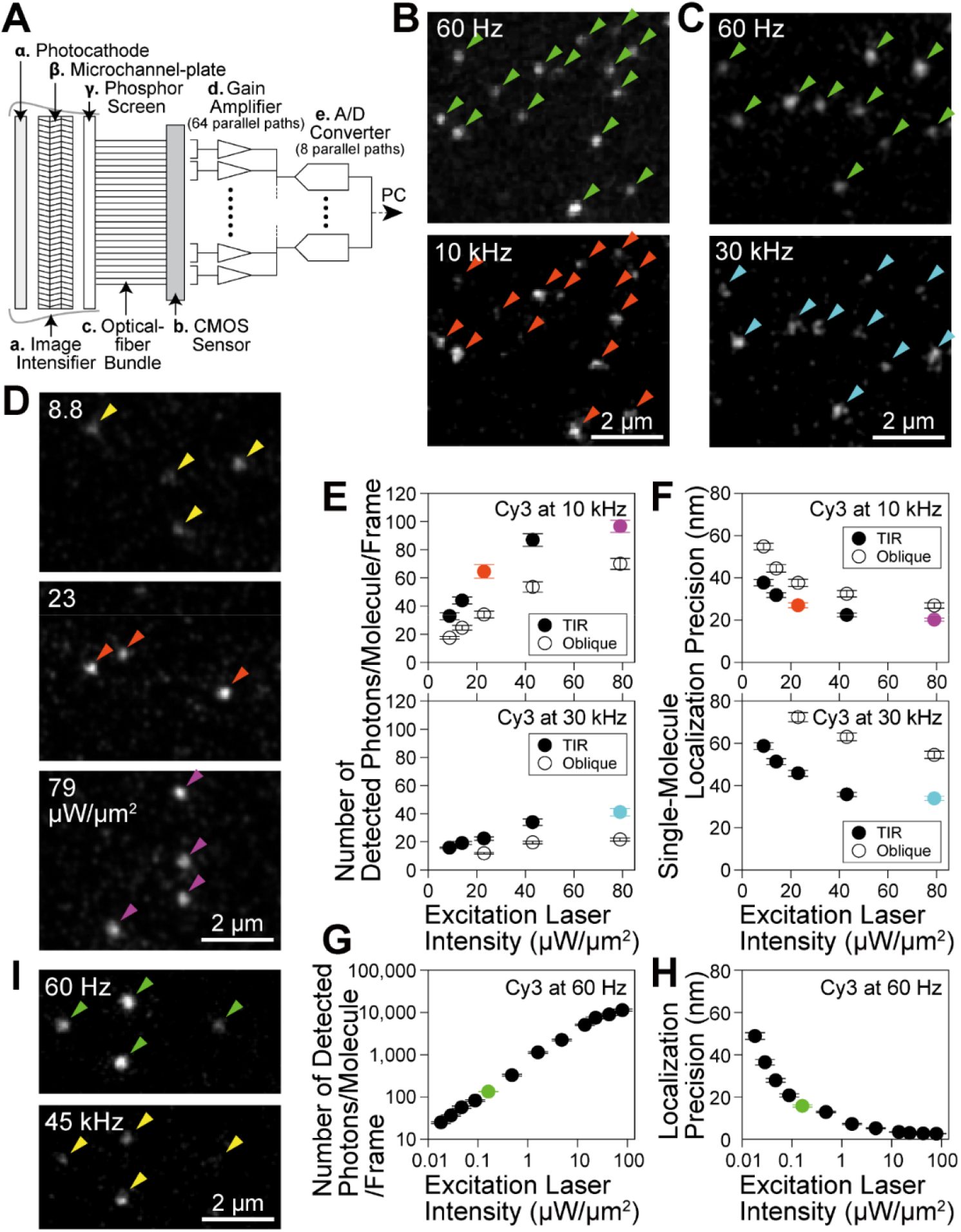
Basic design of the ultrahigh-speed intensified CMOS camera system and ultrafast single fluorescent-molecule imaging (TIR illumination) of Cy3 molecules immobilized on coverslips. The mean ± SEM and/or the median values are provided in the figures throughout this report. (**A**) Schematic diagram of the camera system. See **Materials and methods**. (**B, C**) All of the single Cy3 molecules observed at 60 Hz were detectable at 10 (**B**) and 30 (**C**) kHz (the same field of view); typical snapshots. (**D**) Typical snapshots of single Cy3 molecules excited at various laser powers, observed at 10 kHz. (**E**) The number of detected photons/molecule/frame emitted from single Cy3 molecules during frame times of 0.1 ms (10 kHz) and 0.033 ms (30 kHz) (*n* = 50 Cy3 molecules for each measured point; the same for **F**, **G**, and **H**), plotted against the excitation laser power density. See the caption to Fig. S3 A. (**F**) The localization errors for single Cy3 molecules imaged at 10 and 30 kHz, plotted against the excitation laser intensity. Orange, cyan, and purple data points are color-matched with the arrowheads in (**B**), (**C**), and (**D**). (**G, H**) The number of detected photons/molecule/frame (**G**) and the localization error (**H**) for single Cy3 molecules imaged at 60 Hz, plotted as a function of the excitation laser power density. See the caption to Fig. S2 D. The green data points are color-matched with the arrowheads in (**B**), (**C**), and (**I**). (**I**) Virtually all of the single 5xCy3-Tf molecules (3-8 molecules of Cy3 bound to Tf) were detectable at 45 kHz (compare with the 60-Hz image); typical snapshots.

A straight (1:1) optical-fiber bundle (Fig. 1 A-c) was directly adhered to the phosphor screen of the image intensifier on one side (Fig. 1 A-a-γ, input side) and to the CMOS sensor on the other side (Fig. 1 A-b, output side). This enhanced the signal reaching the CMOS sensor by a factor of 5-10, as compared with the lens coupling used for our standard camera system employed for single fluorescent-molecule imaging (Suzuki et al., 2012; Kinoshita et al., 2017). The electrons generated at the CMOS sensor were transferred, amplified (Fig. 1 A-d), and digitized (Fig. 1 A-e) in 64 and 8 parallel paths, respectively, and then transferred to the host PC.

### Basic conceptual strategy to address the large readout noise of the CMOS sensor

We used the CMOS sensor instead of the sCMOS sensor, which is prevalently used in biomedical research. This is because the frame rates of the sCMOS sensor cannot reach 10∼100 kHz (10,000∼100,000 frames per second), the speed that we hoped to achieve, in contrast to the CMOS sensor. However, the readout noise of the CMOS sensor is generally greater than that of the sCMOS sensor by factors of several 10s (this is why CMOS sensors are rarely used in fluorescence microscopy). Therefore, to employ the CMOS sensor for single-molecule imaging, the problem of high levels of readout noise had to be solved.

Our basic conceptual strategy to address this problem was the following: we should place an amplifier before the noisy detector, so that both the input signal and noise can be amplified (they will be amplified in the same way by the amplifier) at least to the level that the amplified *noise* (be mindful that this must be the noise and not the signal) becomes comparable to the *noise* that the detector generates. In the case of the CMOS camera (noisy detector), we placed an image intensifier (more specifically the microchannel plate; **a-β** in Fig. 1 A), before the CMOS sensor (**b** in Fig. 1 A), so that the *noise* (including the background) generated by the photocathode of the image intensifier (the first step of light detection; **a-α** in Fig. 1 A) is amplified at least to the level comparable to the readout noise of the CMOS sensor (signal at the photocathode is amplified by the same factor as the noise, but in the present argument, the amplification of the *noise* is the critical issue). Here, we explain why this basic strategy would work.

We refer to the average intensities of the signal and noise + background at the photocathode of the image intensifier as *S*_p_ and *N*_p_, respectively, that of the readout noise as *N*r, and the image intensifier amplification gain as *G*. Note that when using the CMOS sensor for single-molecule observations, generally *N*_p_ << *N*r.

The signal-to-noise (including the background intensity) ratio (*S* /*N*) of the output from the CMOS sensor can be written as a function of *G*,

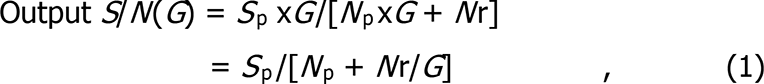

because both the signal and noise at the photocathode stage are amplified by a factor of *G* (neglecting the noise generated by the amplifier). Note that when there is no gain; i.e., *G* =1,

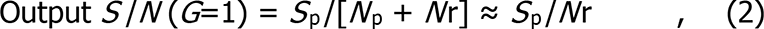

because *N*_p_ << *N*r. Namely, the *S* /*N*ratio of the CMOS sensor output is dominated by the large readout noise of the CMOS sensor *N*r, and we will not be able to detect the single-molecule signal, *S*_p_, due to the large *N*r (i.e., *S*_p_ /*N* (*G*=1) < 1).

With an increase of the gain *G*, *N*r/*G* becomes much smaller than *N*_p_ (or *N*r << *N*_p_ x*G*; namely, when the *noise* at the photocathode is amplified so that the amplified *noise* becomes greater than the readout noise of the CMOS sensor *N*r) and Eq. 1 can be written as

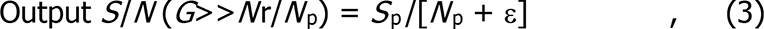

where ε represents the very small value of *N*r/G. Eq. 3 shows that with an increase of the amplifier gain G, the *S*/*N* of the output signal from the CMOS sensor is no longer dominated by *N*r, and thus increased close to that at the photocathode (*S*_p_/*N*_p_). Eq. 3 is consistent with the general truth that amplifier placement cannot increase the output *S*/*N* to more than the input *S*/*N*. Note that, in this argument, the key is not to compare *S*_p_ with *N*r, but to compare *N*_p_ (or *N*_p_ x*G*) with *N*r.

Namely, when the detector itself is a large noise generator and the limiting factor for the *S*/*N* of the entire system, this problem can be minimized by placing an amplifier before the noisy detector. Eqs. 1 and 3 indicate that larger gains are preferrable. However, since the amplifier itself can also generate noise and since the full-well capacity of each pixel of the CMOS sensor is limited, there is an upper limit for the amplifier gain.

When we set the gain *G* so that *N*_p_ x*G* = *N*r (the noise at the photocathode *N*_p_ is amplified to become comparable to the readout noise *N*r); i.e., if we set *G* = *N*r /*N*_p_,

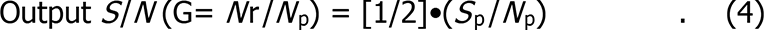

Namely, under these conditions, the output S/N of the CMOS sensor is half the S/N at the photocathode output. This provides a rule of thumb that the noise at the photocathode (*N*_p_) should be amplified at least to a level comparable to the readout noise of the CMOS sensor (*N*r). For the actual values of *N*r and *N*_p_, see **Materials and methods**, “Ultrahigh-speed intensified CMOS camera system: design and operation-*Basic design concept of the camera system* (B) The use of an image intensifier”.

Apart from the basic concept, for determining the useful amplifier gain for experiments, we obtained actual image readout noise, images of single photons obtained as a function of the intensifier gain, stochastic gain variations (fluctuations) of the image intensifier, and the probability of detecting single photons (Fig. S1; **Materials and methods**, “Ultrahigh-speed intensified CMOS camera system: design and operation-*Basic design concept of the camera system* (B) The use of an image intensifier”). We decided to generally employ the overall electron amplification gain of 8,100x, because it enables the detection of a photon-converted electron emitted at the photocathode of the image intensifier with a probability as high as 90.0% (Fig. S1; **Materials and methods**).

The developed camera system is typically operated at 10 and 30 kHz (100- and 33-µs resolutions) with frame sizes of 256 x 256 and 128 x 112 pixels, respectively, but can be operated faster with reduced image sizes (for example at 45 kHz or every 22 µs, with a frame size of 7 x 3.5 µm^2^). Virtually all of the single Cy3 molecules immobilized on coverslips that were observed at 60 Hz were also detectable at frame rates of 10 and 30 kHz (Fig. 1, B-D).

### Ultrafast imaging of many single molecules at 10 and 30 kHz

The single-molecule localization precision is fundamentally determined by the number of detected photons/molecule during the single frame time of the camera, as described by the Mortensen equation (Mortensen et al., 2010; see the caption to Fig. S2). We thus evaluated whether a sufficient number of photons can be emitted from single fluorescent molecules of various fluorophores during the 33 and 100 µs frame times of the developed camera system (30 and 10 kHz, respectively; neglecting a readout time of 812 ns). The plots of single-molecule localization precision vs. the number of detected photons/molecule during a single-frame time (using fluorescent molecules bound to the coverslip) were consistent with the Mortensen equation for all dye molecules investigated here (Fig. S2). These plots demonstrated that the number of detected photons/molecule/frame and the single-molecule localization precision were enhanced with an increase of the excitation laser intensity, but to a limited degree (Fig. 1, E **and** F; Fig. S3). This occurs because the number of photons emitted from a single fluorescent molecule during a frame time under higher excitation light intensities is limited by the triplet bottleneck saturation (**Materials and methods**, “Estimation of the number of photons that can be emitted by a single Cy3 molecule during 0.1 ms: triplet bottleneck saturation”). The saturation became apparent from around 23 µW/µm^2^ for Cy3 (Fig. 1, E and F; Fig. S3). Among the eight dyes we tested, Cy3 exhibited the lowest tendency to saturate (Figs. S2 B and S3 A). Therefore, we employed Cy3 most often, but also used tetramethylrhodamine (TMR) as a membrane-permeable probe.

The best single-molecule localization precision for Cy3 at a frame integration time of 0.1 ms (10-kHz frame rate) was 20 ± 0.71 nm, using total internal reflection (TIR) laser illumination at a density of 79 µW/µm^2^ at the sample plane (*n* = 50 molecules; mean ± standard error of means [SEM] are given throughout this report; Fig. 1 F; Figs. S2, A and B and S3, B and C). At 30 kHz, since fewer photons were emitted from a Cy3 molecule during a frame time of 33 µs, the localization precision was worse (34 ± 1.0 nm; *n* = 50 molecules; Fig. 1 F; Figs. S2 A and S3, B **and** C; **Table 1**). The best single-molecule localization precision achieved with this camera system was 2.6 ± 0.099 nm, with a maximum number of detected photons/molecule/frame of 11,400 ± 700, at a frame integration time of 16.7 ms (60-Hz frame rate) in which most of the Cy3 molecules were photobleached (within a single frame; *n* = 50 molecules; Fig. 1, G **and** H; Fig. S2, C **and** D) (summarized in **Table 1**).

Taken together, we conclude that the observation frame rates of 10 – 30 kHz (with localization precisions of 20 – 34 nm, using the view-fields of 256 x 256 – 128 x 112 pixels [14.1 x 14.1 µm^2^ – 7.1 x 6.2 µm^2^]), represent the ultimate fastest frame rates employable for single-molecule tracking in living cells, using the currently available fluorescent molecules. Meanwhile, the camera itself could be operated much faster. The Photron 1024PCI CMOS sensor, which was used most extensively here, can be operated at 10 kHz (256 x 256 pixels), 45 kHz (128 x 64 pixels; or 256 x 256 pixels using the newer SA1 sensor), and 110 kHz (128 x 16 pixels) (summarized and compared with the results with other cameras in **Table 1**; **Materials and methods**, “Ultrahigh-speed intensified CMOS camera system: design and operation”). Therefore, the camera is no longer the limitation for the faster frame rates, and with the future development of fluorescent dyes with higher throughputs, we should be able to perform single-molecule tracking at even better time resolutions. In fact, single transferrin (Tf) molecules bound by an average of 5.0 Cy3 molecules (5xCy3-Tf) adsorbed on the coverslip could be imaged and tracked at 45 kHz, with localization precisions of 30 (38) and 29 (34) nm, using TIR (oblique-angle) illuminations of 43 and 79 µW/µm^2^, respectively (Figs. S2 A and S3, B **and** C; **Table 1**).

### TIR and oblique illuminations for ultrafast single-molecule imaging

In this study, we employed both TIR and oblique-angle laser illuminations. TIR illumination is useful for observing single molecules in the basal (ventral, bottom) PM by suppressing the signals from the cytoplasm and the enhanced evanescent electric field for the same laser intensity. The TIR illumination provides better localization precisions than the oblique-angle illumination at the same laser intensities (27 vs. 38 nm; Fig. 1 F).

On the other hand, the oblique-angle illumination is more versatile: it can be used to illuminate the apical PM, endomembranes, and cytoplasm, which cannot be accomplished using the TIR illumination. Particularly, in this research, since we hoped to compare the present single fluorescent-molecule tracking results of the phospholipid diffusion with our previous 40-kHz single gold-probe tracking data obtained in the *apical* PM (Fujiwara et al., 2002, 2016; Murase et al., 2004), we had to use the oblique-angle illumination. Therefore, as our “standard test conditions” in the present research (the end of the caption to Fig. S3), we employed the oblique-angle illumination with a laser intensity of 23 µW/µm^2^ at the sample plane (the dye saturation starts around this laser intensity; Fig. 1, E **and** F; Fig. S3), despite its worse localization precisions as compared with the TIR illumination.

### Trajectory length and cell viability under the standard conditions

Under the standard test conditions (oblique-angle illumination at 23 µW/µm^2^), single Cy3 molecules immobilized on the coverslip could be observed at 10 kHz with an average signal-to-noise ratio (SNR) of 2.5 ± 0.11 (Fig. 2, A **and** B; **Video 1**), and the fractions of the trajectories with durations (uninterrupted length of the trajectories) longer than 100, 300, and 1,000 frames (after the 3-frame gap-closing, which neglects the non-detectable periods lasting for 3 frames or less) were 14, 2.9, and 0.31% among all obtained trajectories (Fig. 2 C). Therefore, performing single-molecule tracking for 100 – 300 frames (10 – 30 ms at 10 kHz) is quite practical. Meanwhile, under the conditions for the best localization precision with TIR illumination of 79 µW/µm^2^ (20 ± 0.71 nm), the fractions were 1.6, 0.13, and 0.0%, respectively (Fig. 2 C), making it difficult to obtain trajectories longer than 100 frames (for the summary and comparison with results using other cameras, see **Table 1**).

**Figure 2.**
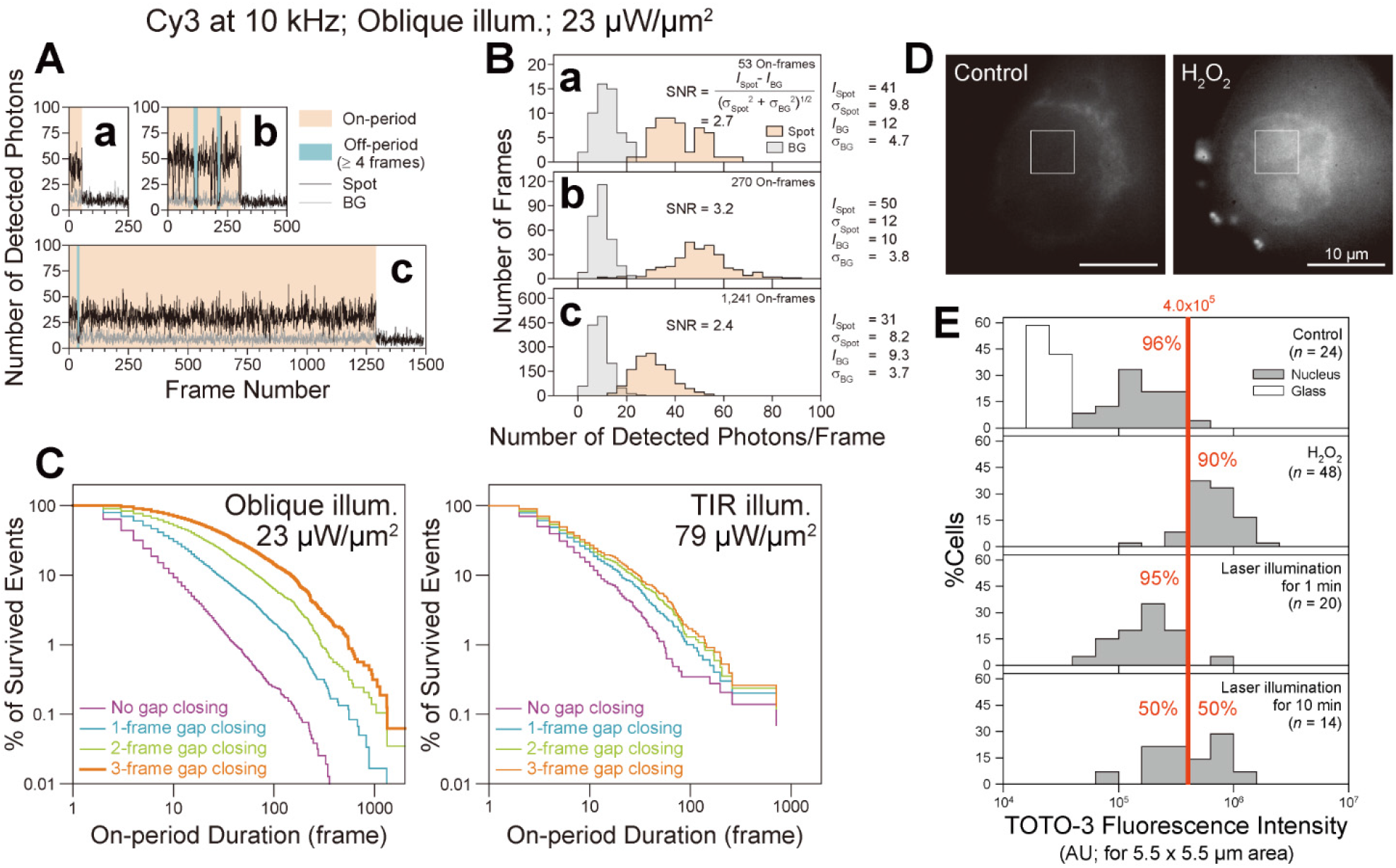
The observable signal durations of single Cy3 molecules immobilized on a coverslip, which are limited by photoblinking/photobleaching, stochastic fluctuations of the signal, and photon shot noise (A - C), and minimal photo-induced damage to the cells during ultrahigh-speed single fluorescent-molecule imaging (D, E). (**A**) Typical time-dependent fluorescence signal intensities (number of detected photons/0.1-ms-frame) of three single Cy3 molecules (**a**, **b**, and **c**) excited by oblique-angle illumination at 23 µW/µm^2^, and observed at 10 kHz (background signal in grey). See **Materials and methods** for details. (**B**) The signal-to-noise ratios (SNRs) for the images of the three molecules shown in (**A**) are in the range between 2.4 and 3.2. See **Materials and methods**. (**C**) The distributions of the durations of the on-periods and those after neglecting the off-periods (non-emission periods) lasting for 1, 2, or 3 frames (gap closing; see **Materials and methods**) of single Cy3 molecules immobilized on a coverslip and excited by oblique-angle illumination at 23 µW/µm^2^ (standard conditions; **left**) or by TIR illumination at 79 µW/µm^2^ (**right**), observed at 10 kHz (totals of 264 and 593 molecules, respectively). The 3-frame gap closing was employed in single-molecule tracking under the standard test conditions (thick orange curve; **left**). (**D, E**) Photo-induced damage to the cells during ultrahigh-speed single fluorescent-molecule imaging. Cell viability was examined by staining with 1 µM TOTO-3 iodide, which only stains dead cells, at 37°C for 5 min, and then observing the stained cells using epi-illumination with a 594-nm laser. (**D**) Representative images of the nuclei stained with TOTO-3 iodide. Control, a reference image of a living cell (*n* = 24 images). H_2_O_2_, a reference image of a dead cell after the treatment with 100 µM H_2_O_2_ at 37°C for 1 h (*n* = 48 images). (**E**) Histograms showing the fluorescence intensity of TOTO-3 in the 5.5 x 5.5-µm area inside the nucleus (see the square box in A) (*n* = the number of examined cells). (**Top**) Live cells without laser illumination (negative control). (**Second**) Dead cells after the H_2_O_2_ treatment (positive control). Based on the results of the negative and positive controls (top and second boxes, respectively), a threshold fluorescence intensity of 4.0 x 10^5^ (arbitrary unit = AU) was selected to categorize the live and dead cells (96% [90%] of the negative [positive] control cells were categorized as alive [dead]). (**Third and Fourth**) Cells were illuminated under our typical 10-kHz single Cy3 molecule imaging conditions (oblique-angle 532-nm laser illumination at 23 µW/µm^2^) for 1 and 10 min, respectively, which are longer by factors of 12 and 120 than our longest illumination duration of 5 s/cell (ten 500-ms image sequences = 5 x 10^4^ frames). We concluded that the light-induced damage to the cells is very limited under our standard experimental conditions.

These results indicate that, for successful single-molecule tracking, a proper compromise between the localization precision and the trajectory length is necessary. This is one of the major differences between single-molecule tracking and SMLM. SMLM basically requires only one localization for a single molecule (observations lasting only for a single frame) before photobleaching, while single-molecule tracking needs many localizations to obtain long trajectories (essentially, longer is better) before photobleaching/photoblinking.

Under the standard test conditions, the laser illumination (oblique-angle at 23 µW/µm^2^ of the 532-nm line) for 1 min hardly affected the cell viability, although half of the T24 cells did not survive after 10-min irradiation (Fig. 2, D **and** E). Because all our ultrafast measurements were finished within 5 s, we conclude that the toxic effect of the illumination laser is minimal for our observations. In the previous report (Wäldchen et al., 2014), the light toxicity was found strongly dependent on the cell type, and our result using T24 cells is close to their result using HeLa cells. However, the direct comparison with the present result with the previous result is difficult due to the differences in the viability test method, laser wavelength (532 vs. 514 nm), and illumination durations (1 and 10 min vs. 4 min; our result suggests non-linear dependence of the viability on the illumination duration).

### Testing the ultrafast camera system using PM molecules undergoing hop diffusion 1: qualitative observations

As stated in the **Introduction**, we previously detected non-Brownian diffusion (hop diffusion) of phospholipid and transmembrane protein molecules in the PM, using large (40-nm diameter) gold particles as a probe (Fujiwara et al., 2002, 2016; Murase et al., 2004; Kusumi et al., 2005, 2012a, b; also see Sheetz, 1983). This was made possible by improving the time resolution of single-particle imaging using bright-field microscopy down to 25 µs (Fujiwara et al., 2002, 2016; Murase et al., 2004). The hop diffusion is influenced by modulating the actin-based membrane-skeleton meshes (Fig. 3 A). Therefore, we concluded that the entire apical PM is compartmentalized, by the steric hindrance of actin-based membrane-skeleton meshes (fences) and the steric hindrance + hydrodynamic friction-like effects from rows of transmembrane-protein pickets anchored to and aligned along the actin fence (Fig. 3 A). In the compartmentalized PM, both transmembrane proteins and lipids undergo short-term confined diffusion within a compartment plus occasional hop movements to an adjacent compartment, which was termed hop diffusion (Kusumi et al., 2005; Kusumi et al., 2012a; Jacobson et al., 2019).

**Figure 3.**
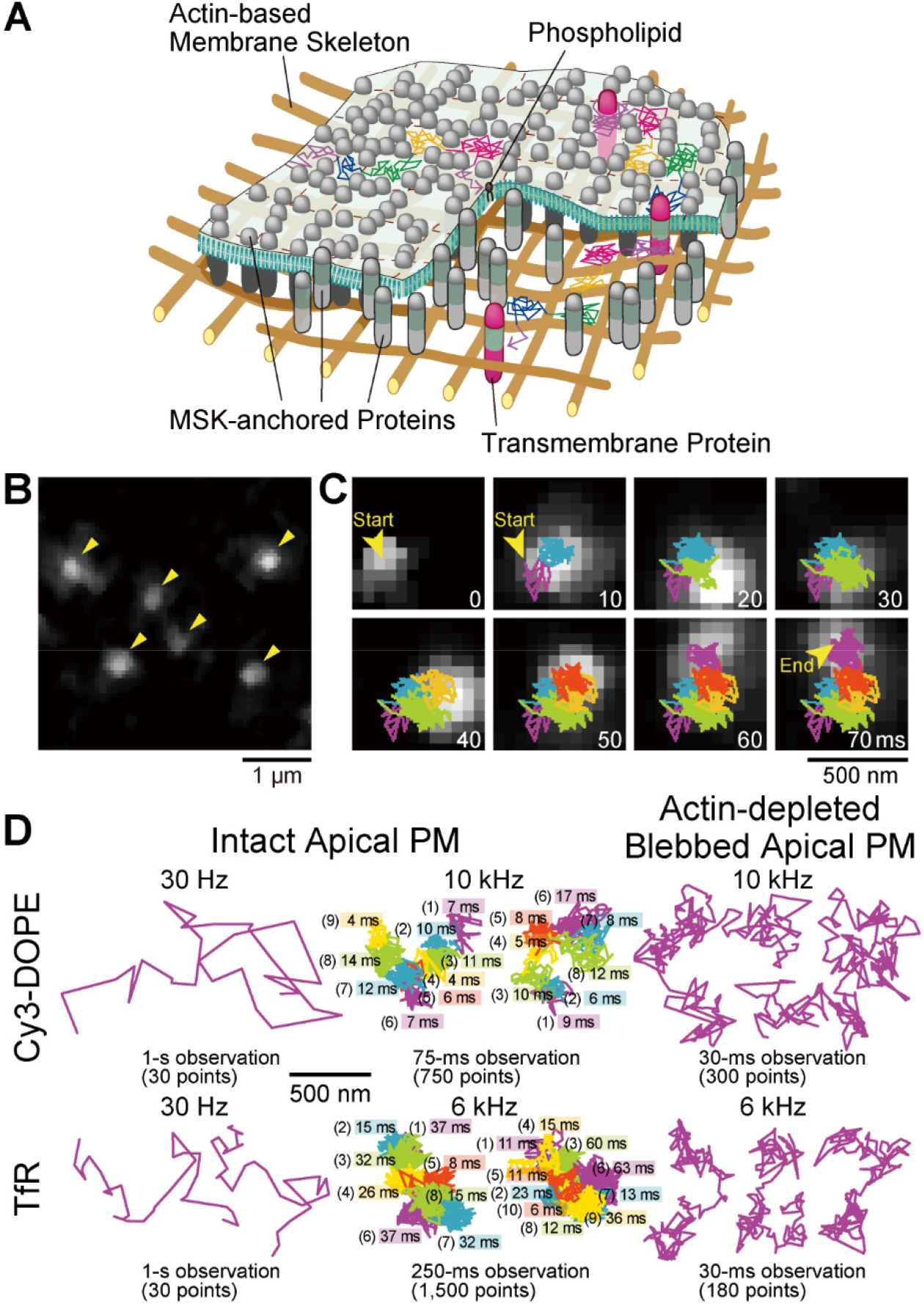
Ultrafast single fluorescent-molecule imaging and tracking of Cy3-labeled DOPE and TfR in the apical PM. (**A**) Schematic drawing showing that membrane molecules undergo hop diffusion in the PM, which is compartmentalized by the actin-based membrane-skeleton meshes (fences; brown mesh) and rows of transmembrane-protein pickets anchored to and aligned along the actin fence (grey molecules). (**B**) A typical snapshot of single Cy3-DOPE molecules in the apical PM observed under our standard conditions, with an integration time of 0.1 ms. (**C**) A representative image sequence of a single Cy3-DOPE molecule diffusing in the apical PM, recorded every 0.1 ms (shown every 100 image frames = every 10 ms). The colors in the trajectory indicate the diffusion in different plausible compartments (order of appearance: purple, blue, green, orange, red, and then back to purple; the same color order is used throughout this report), detected by the TILD analysis software (Fig. S5). (**D**) Typical single-molecule trajectories of Cy3-DOPE and Cy3-Tf-TfR in intact and actin-depleted blebbed apical PMs, recorded at normal video rate and enhanced rates. The order of the compartments that the molecule visited (parenthesized integers) and the residency times there, as determined by the TILD analysis, are indicated (intact PMs).

Each gold-tagged molecule in the PM exhibited two diffusion coefficients: the microscopic diffusion coefficient (*D*_micro_), which describes the unhindered diffusion within a compartment, and the macroscopic diffusion coefficient (*D*_MACRO_), which is largely determined by the compartment size and the hop frequency across intercompartmental boundaries composed of the picket-fence. *D*_MACRO_ is significantly smaller than *D*_micro_.

The compartment sizes were in the range of 40 - 230 nm depending on the cell type (∼110 nm in human epithelial T24 cell line used here), and the dwell lifetimes of membrane molecules within a compartment are in the range of several to a few 10s of milliseconds (Murase et al., 2004; Fujiwara et al., 2016). Therefore, the hop diffusion of membrane molecules in the apical PM appeared to be quite suitable for testing single fluorescent-molecule imaging using the developed ultrafast camera system. Indeed, this project was originally undertaken to develop a fast single *fluorescent*-molecule imaging method to observe the hop diffusion of membrane molecules in the PM. We examined whether we could detect the hop diffusion of L-α-dioleoylphosphatidylethanolamine (DOPE) conjugated with Cy3, Cy3-DOPE, at a time resolution of 0.1 ms (10 kHz, 333 times faster than normal video rate) and transferrin receptor (TfR) tagged with Cy3-conjugated transferrin (average dye to protein molar ratio of 0.2) at a frame rate of 6 kHz (0.167-ms resolution; 200 times faster than normal video rate; slowed from 10 kHz because the hop rate of TfR was expected to be slower than that of Cy3-DOPE). The observations were made in the apical PM using the oblique-angle illumination (Fig. 3, B-D; **Videos 2-4**), for making the direct comparisons with the previous single gold-particle tracking data. All of the cell experiments reported here were performed at 37°C, using human epithelial T24 cells (previously called ECV304 cells).

Virtually all of the single-molecule images and trajectories (Fig. 3, B-D; **Videos 2-4**) gave the impression that the Cy3-DOPE and TfR molecules underwent rapid diffusion within a confined domain of ∼100 nm and occasionally moved out of this domain, became confined again at the place where it moved to, and repeated such behaviors. These typical behaviors appear to reproduce the movement of gold-tagged DOPE and TfR quite well.

### Testing the ultrafast camera system using PM molecules undergoing hop diffusion 2: quantitative and statistical analyses

Apart from the subjective impression, the trajectories of single fluorescent molecules were examined quantitatively and statistically, using the plot of mean-square displacement (MSD) vs. time interval (Δ*t*) (Qian et al., 1991) (called the MSD-Δ*t* plot, which also provides single-molecule localization precisions; Fig. 4 A; Fig. S4). The results were compared with those obtained by single gold-particle imaging.

**Figure 4.**
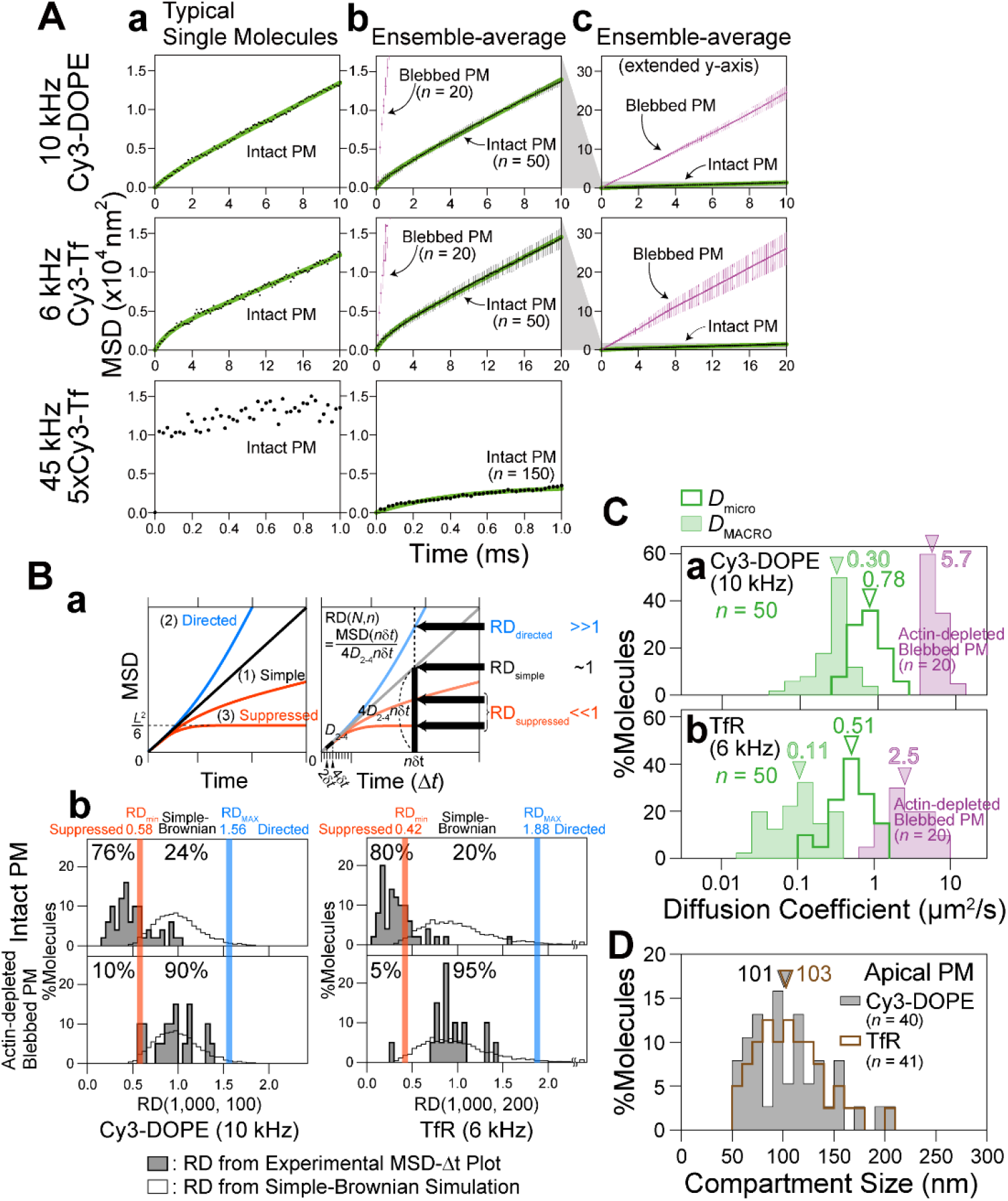
Examination of whether our ultrafast single fluorescent-molecule imaging can detect hop diffusion of Cy3-DOPE and TfR in the *apical* PM, previously found by ultrafast single gold-particle tracking. (**A**) Typical MSD-Δ*t* plot for a single molecule (**a**), and the ensemble-averaged MSD-Δ*t* plots with two different y-axis scales (**b, c**: 20x of **b**) in the intact *apical* PM (black) and the *blebbed apical* PM (purple). The MSD-Δ*t* plots of the top and middle boxes in (**b**) are those after subtracting 4σ_xy_^2^, shown in Fig. S4 (σ_xy_ = single-molecule localization precision). Only in the typical MSD-Δ*t* plot for a single 5xCy3-Tf molecule bound to TfR obtained at 45 kHz (**a, bottom**), 4σ_xy_^2^ was not subtracted due to large errors in its estimation. Green curves are the best-fit functions for the hop-diffusion fitting (**top and middle rows**) and confined-diffusion fitting (**bottom row** for 45 kHz observations of 5xCy3-Tf). (**B**) The method for the statistical classification of each single-molecule trajectory into a suppressed-, simple-Brownian-, or directed-diffusion mode. (**a**) The basic idea for the classification: the parameter RD (relative deviation) describes the extent to which the observed diffusion deviates from simple-Brownian diffusion at a time sufficiently later from time 0; i.e., the actual MSD divided by the calculated MSD from the short-term diffusion coefficient (*D*2-4) assuming simple-Brownian diffusion. See **Materials and methods**. The RD value is ≪, ≈, or ≫ 1, when the molecules are undergoing suppressed, simple-Brownian, or directed diffusion, respectively (Fujiwara et al., 2002; Kusumi et al., 1993; Murase et al., 2004; Suzuki et al., 2005). The suppressed-diffusion mode includes both the confined-diffusion and hop-diffusion modes. (**b**) Classification of individual trajectories based on the RD histograms for simulated simple-Brownian particles (open bars; n = 5,000). The RD values giving the 2.5 percentiles of the particles from both ends of the distribution, referred to as *RD*_min_ and *RD*_MAX_, were obtained (red and blue vertical lines, respectively). Each experimental single-molecule trajectory was classified into suppressed (confined and hop), simple-Brownian, or directed diffusion when its RD value was smaller than *RD*_min_, between *RD*_min_ and *RD*_MAX_, and greater than *RD*_MAX_ (no trajectory fell in this category), respectively. The distributions of *RD*s for the Cy3-DOPE and TfR trajectories are shown by shaded histograms (n = 50 and 20 for the intact and actin-depleted blebbed PM). (**C**) The distributions of *D*_micro_ (=*D*_2-4_; underestimated due to the insufficient time resolution even at a 0.1-ms resolution), *D*_MACRO_ in the apical PM, and simple-Brownian diffusion coefficients in the blebbed apical PM for Cy3-DOPE (**a**) and Cy3-Tf bound to TfR (**b**). Arrowheads indicate the median values (summarized in **Table 1**). These diffusion coefficients in the blebbed PM are slightly smaller than those obtained with gold probes (Fujiwara et al., 2002, 2016). This is probably due to the residual actin filaments in the T24 cells employed here, since the membrane-bound actin filament meshes are much denser in T24 cells than the NRK cells used previously. (**D**) The distributions of the compartment sizes, determined by the hop-diffusion fitting of the MSD-Δ*t* plot for each trajectory of Cy3-DOPE and TfR. Arrowheads indicate the median values. All of the statistical parameters and the statistical test results throughout this report are summarized in **Table 1**.

Based on the MSD-Δ*t* plot (Fig. 4 A-a), each trajectory could be classified into a suppressed-, simple-Brownian-, or directed-diffusion mode (Kusumi et al., 1993; see the caption to Fig. 4 B-a). More than three-quarters of the Cy3-DOPE and TfR trajectories obtained in the intact apical PM at 10 and 6 kHz (0.1-ms and 0.167-ms resolutions, respectively) were categorized into the suppressed-diffusion mode (top boxes in Fig. 4 B-b; also see **Table 2**), reproducing the results obtained by single gold-particle tracking at 40 kHz (**Table 2**, in which all of the statistical parameters and the statistical test results for the diffusion parameters are also summarized).

**Table 2.**
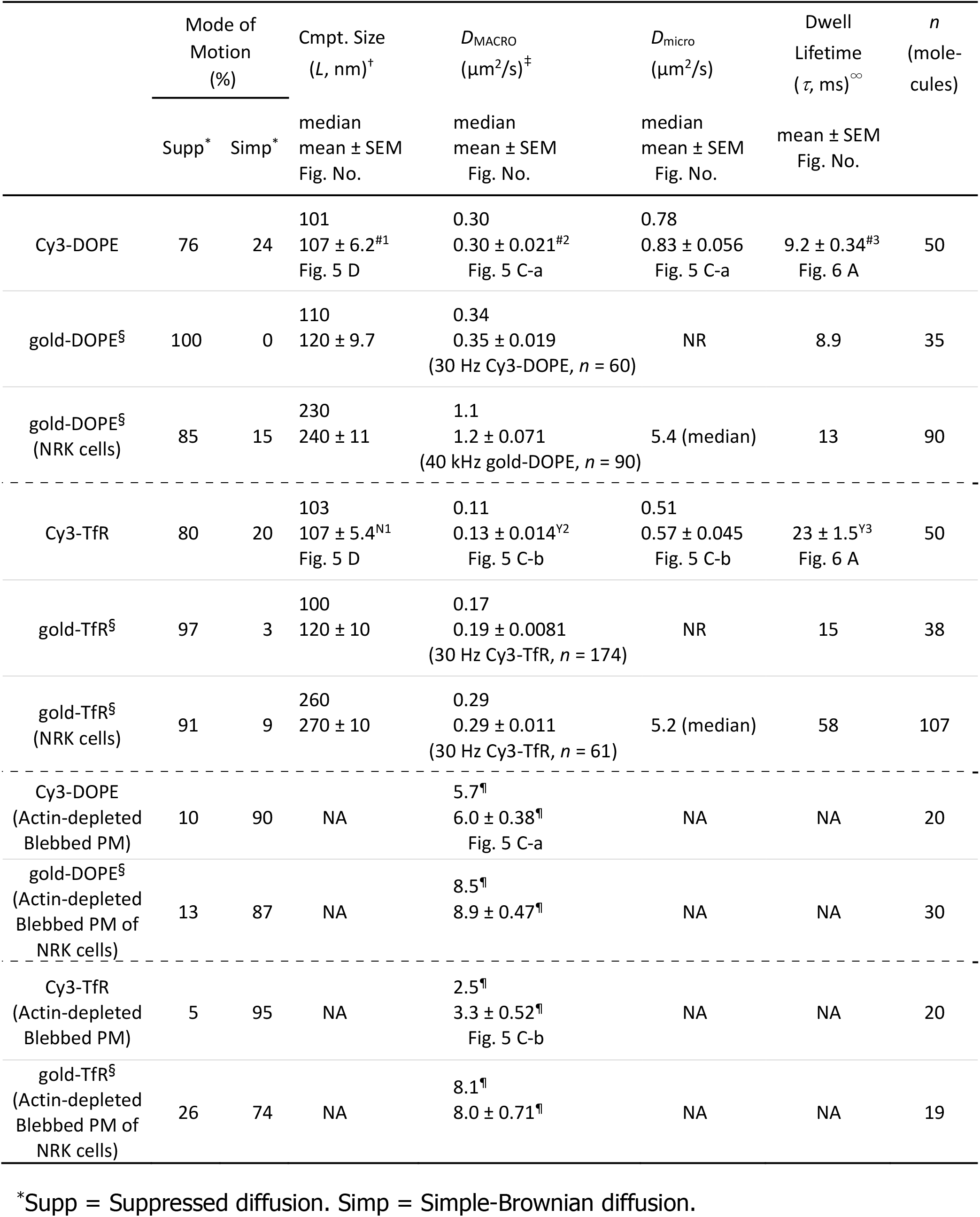
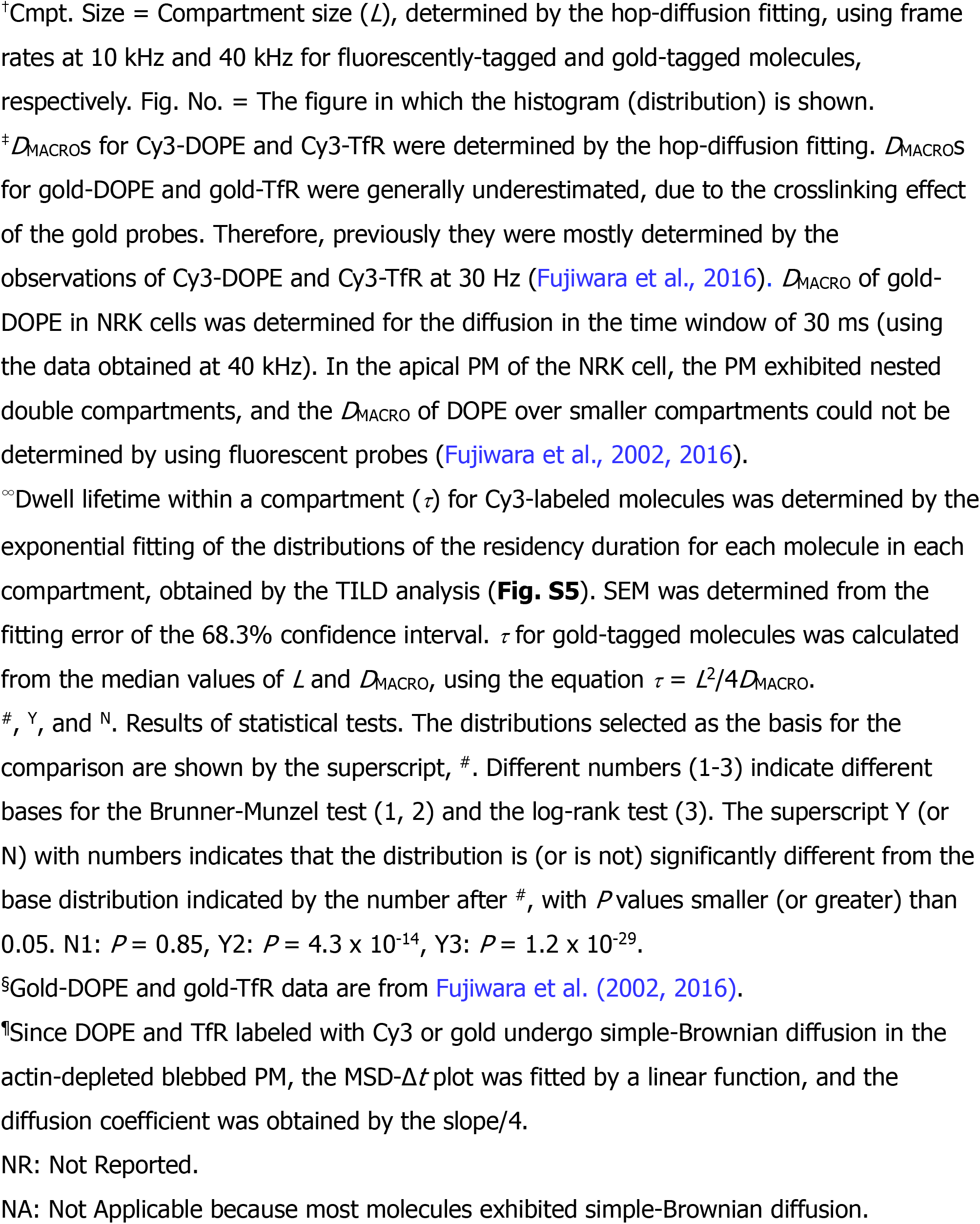
Examination of the ultrafast single fluorescent-molecule imaging developed here to determine whether it can correctly evaluate hop diffusion parameters for Cy3-DOPE and TfR in the apical PM, previously found by ultrafast gold-particle imaging (dwell lifetimes were previously determined by using additional data from video-rate observations of fluorescently-labeled molecules).

| | Mode of Motion (%) | | Cmpt. Size ($L$, nm)<sup>†</sup> | $D_{\text{MACRO}}$ ($\mu\text{m}^2/\text{s}$)<sup>‡</sup> | $D_{\text{micro}}$ ($\mu\text{m}^2/\text{s}$) | Dwell Lifetime ($\tau$, ms)<sup>∞</sup> | $n$ (molecules) |
| --- | --- | --- | --- | --- | --- | --- | --- |
| | Supp<sup>*</sup> | Simp<sup>*</sup> | median mean ± SEM<br>Fig. No. | median mean ± SEM<br>Fig. No. | median mean ± SEM<br>Fig. No. | mean ± SEM<br>Fig. No. | |
| Cy3-DOPE | 76 | 24 | 101<br>107 ± 6.2<sup>#1</sup><br>Fig. 5 D | 0.30<br>0.30 ± 0.021<sup>#2</sup><br>Fig. 5 C-a | 0.78<br>0.83 ± 0.056<br>Fig. 5 C-a | 9.2 ± 0.34<sup>#3</sup><br>Fig. 6 A | 50 |
| gold-DOPE<sup>§</sup> | 100 | 0 | 110<br>120 ± 9.7 | 0.34<br>0.35 ± 0.019<br>(30 Hz Cy3-DOPE, $n$ = 60) | NR | 8.9 | 35 |
| gold-DOPE<sup>§</sup><br>(NRK cells) | 85 | 15 | 230<br>240 ± 11 | 1.1<br>1.2 ± 0.071<br>(40 kHz gold-DOPE, $n$ = 90) | 5.4 (median) | 13 | 90 |
| Cy3-TfR | 80 | 20 | 103<br>107 ± 5.4<sup>N1</sup><br>Fig. 5 D | 0.11<br>0.13 ± 0.014<sup>Y2</sup><br>Fig. 5 C-b | 0.51<br>0.57 ± 0.045<br>Fig. 5 C-b | 23 ± 1.5<sup>Y3</sup><br>Fig. 6 A | 50 |
| gold-TfR<sup>§</sup> | 97 | 3 | 100<br>120 ± 10 | 0.17<br>0.19 ± 0.0081<br>(30 Hz Cy3-TfR, $n$ = 174) | NR | 15 | 38 |
| gold-TfR<sup>§</sup><br>(NRK cells) | 91 | 9 | 260<br>270 ± 10 | 0.29<br>0.29 ± 0.011<br>(30 Hz Cy3-TfR, $n$ = 61) | 5.2 (median) | 58 | 107 |
| Cy3-DOPE<br>(Actin-depleted<br>Blebbled PM) | 10 | 90 | NA | 5.7<sup>¶</sup><br>6.0 ± 0.38<sup>¶</sup><br>Fig. 5 C-a | NA | NA | 20 |
| gold-DOPE<sup>§</sup><br>(Actin-depleted<br>Blebbled PM of<br>NRK cells) | 13 | 87 | NA | 8.5<sup>¶</sup><br>8.9 ± 0.47<sup>¶</sup> | NA | NA | 30 |
| Cy3-TfR<br>(Actin-depleted<br>Blebbled PM) | 5 | 95 | NA | 2.5<sup>¶</sup><br>3.3 ± 0.52<sup>¶</sup><br>Fig. 5 C-b | NA | NA | 20 |
| gold-TfR<sup>§</sup><br>(Actin-depleted<br>Blebbled PM of<br>NRK cells) | 26 | 74 | NA | 8.1<sup>¶</sup><br>8.0 ± 0.71<sup>¶</sup> | NA | NA | 19 |

<sup>\*</sup>Supp = Suppressed diffusion. Simp = Simple-Brownian diffusion.
<sup>†</sup>Cmpt. Size = Compartment size ( $L$ ), determined by the hop-diffusion fitting, using frame rates at 10 kHz and 40 kHz for fluorescently-tagged and gold-tagged molecules, respectively. Fig. No. = The figure in which the histogram (distribution) is shown.
<sup>‡</sup> $D_{\text{MACROS}}$ for Cy3-DOPE and Cy3-TfR were determined by the hop-diffusion fitting. $D_{\text{MACROS}}$ for gold-DOPE and gold-TfR were generally underestimated, due to the crosslinking effect of the gold probes. Therefore, previously they were mostly determined by the observations of Cy3-DOPE and Cy3-TfR at 30 Hz (Fujiwara et al., 2016). $D_{\text{MACRO}}$ of gold-DOPE in NRK cells was determined for the diffusion in the time window of 30 ms (using the data obtained at 40 kHz). In the apical PM of the NRK cell, the PM exhibited nested double compartments, and the $D_{\text{MACRO}}$ of DOPE over smaller compartments could not be determined by using fluorescent probes (Fujiwara et al., 2002, 2016).
<sup>∞</sup>Dwell lifetime within a compartment ( $\tau$ ) for Cy3-labeled molecules was determined by the exponential fitting of the distributions of the residency duration for each molecule in each compartment, obtained by the TILD analysis (**Fig. S5**). SEM was determined from the fitting error of the 68.3% confidence interval. $\tau$ for gold-tagged molecules was calculated from the median values of $L$ and $D_{\text{MACRO}}$ , using the equation $\tau = L^2/4D_{\text{MACRO}}$ .
<sup>#</sup>, <sup>Y</sup>, and <sup>N</sup>. Results of statistical tests. The distributions selected as the basis for the comparison are shown by the superscript, <sup>#</sup>. Different numbers (1-3) indicate different bases for the Brunner-Munzel test (1, 2) and the log-rank test (3). The superscript Y (or N) with numbers indicates that the distribution is (or is not) significantly different from the base distribution indicated by the number after <sup>#</sup>, with $P$ values smaller (or greater) than 0.05. N1: $P = 0.85$ , Y2: $P = 4.3 \times 10^{-14}$ , Y3: $P = 1.2 \times 10^{-29}$ .
<sup>§</sup>Gold-DOPE and gold-TfR data are from Fujiwara et al. (2002, 2016).
<sup>¶</sup>Since DOPE and TfR labeled with Cy3 or gold undergo simple-Brownian diffusion in the actin-depleted blebbed PM, the MSD- $\Delta t$ plot was fitted by a linear function, and the diffusion coefficient was obtained by the slope/4.
NR: Not Reported.
NA: Not Applicable because most molecules exhibited simple-Brownian diffusion.

Meanwhile, in the f-actin-depleted blebbed PM, almost all of the Cy3-DOPE and TfR trajectories, obtained at 10 and 6 kHz, respectively, were classified into the simple-Brownian-diffusion mode (Fig. 4 A-c; Fig. 4 B-b; **Table 2**), again consistent with the data obtained by single gold-particle tracking in the blebbed PMs of NRK cells (Fujiwara et al., 2002, 2016). As concluded previously, these results indicate that the actin-based membrane skeleton is involved in inducing the suppressed diffusion of both phospholipids and transmembrane proteins (Fujiwara et al., 2002, 2016; Murase et al., 2004; Hiramoto-Yamaki et al., 2014).

The MSD-Δ*t* plot obtained for each trajectory was then fitted, using an equation based on the hop-diffusion theory (Kenkre et al., 2008) (called “hop-diffusion fitting”; Fig. 4 A; see the subsection “Hop-diffusion fitting: the function describing the MSD-Δt plot for particles undergoing hop diffusion” in the caption to Fig. S5). The hop-diffusion fitting provided two diffusion coefficients for each trajectory observed in the PM: *D*_micro_ and *D*_MACRO_. The presence of two diffusion coefficients for a single trajectory observed in the PM and only one diffusion coefficient in the actin-depleted blebbed PM (Fig. 4 C) again reproduced the observations obtained by using single gold-particle imaging (Fujiwara et al., 2002, 2016).

The hop-diffusion fitting of each trajectory obtained in the intact PM also provided the average compartment size *L* for each trajectory (Fig. 4 D; **Table 2**). Importantly, the *L*’s for Cy3-DOPE and TfR were quite similar, with median values of 101 and 103 nm, respectively (non-significant difference; in the following discussions and calculations, we will use a compartment size of 100 nm). This result suggests that the underlying mechanisms for confining phospholipids and transmembrane proteins are based on the same cellular structures, perhaps by the actin-based membrane-skeleton meshes (fences) and the transmembrane picket proteins bound to and aligned along the actin-fence (Fig. 3 A). The compartment sizes *L* found here (median sizes of 101 and 103 nm for Cy3-DOPE and TfR, respectively) are quite comparable to those previously detected by using gold-conjugated molecules in the same T24 cells (110 and 100 nm for gold-tagged DOPE and TfR, respectively; **Table 2**).

Taken together, our ultrafast single fluorescent-molecule imaging based on the ultrafast camera system developed here could successfully detect the hop diffusion of membrane molecules, which was previously detectable only by using gold probes and high-speed bright-field microscopy. Therefore, the ultrafast single fluorescent-molecule imaging developed here is likely to be suitable to observe the fast molecular dynamics occurring in living cells.

Here, we add a remark on the *D*_micro_s for Cy3- and gold-labeled molecules in the PM. The model of hop diffusion in the compartmentalized PM suggests that *D*_micro_ is about the same as the diffusion coefficient of unhindered diffusion in the actin-depleted PM, but in all the molecules and cells examined thus far, the *D*_micro_s in the intact PM were smaller than those for the unhindered diffusion coefficients in the actin-depleted PM (**Table 2**; Fujiwara et al., 2002, 2016; Murase et al., 2004; Suzuki et al., 2005). This is probably because the time resolutions of 25 - 100 µs achieved thus far (10 - 40 kHz) are not sufficient to observe trajectories near the compartment boundaries, thus effectively reducing the step sizes in the trajectory, which will reduce the diffusion coefficient within the compartment, *D*_micro_ (Ritchie et al., 2005).

We examined the possibility that the hop diffusion might simply be an apparent phenomenon due to the photo response non-uniformity (PRNU) of the developed camera system. The PRNU induced by the effects of pixel-to-pixel variations on the sensitivity of the developed camera system might have made the simple-Brownian diffusion of membrane molecules appear like hop diffusion. The result is shown in Fig. S1 of the companion paper, and indicates that the PRNU of the developed camera system is not more than that of the conventional EM-CCD camera, and thus is unlikely to induce hop diffusion-like movies.

### Direct determination of the dwell lifetime of each membrane molecule within each compartment using ultrafast single fluorescent-molecule imaging

We directly evaluated the dwell lifetime of *each molecule* in *each compartment* by using ultrafast single fluorescent-molecule imaging and developing an improved method for detecting the moment (instance) at which the observed molecule undergoes the hop movement across the compartment boundary. The moments of hops of the observed molecule can be found in its single-molecule trajectory by the method of detecting a <u>T</u>ransient <u>I</u>ncrease of the effective <u>L</u>ocal <u>D</u>iffusion (TILD) in the trajectory (Fig. S5). TILDs are likely to occur when a molecule hops between two membrane compartments, but the analysis itself remains model-independent. TILDs were detected in all of the experimental trajectories obtained at 0.1- and 0.167-ms resolutions (for Cy3-DOPE and TfR, respectively) in the intact PM (see Fig. 3, C **and** D); however, in the blebbed PM, where all of the trajectories were statistically classified into the simple-Brownian diffusion mode, TILDs were detected in only 4% [9%] of the trajectories (one TILD per 750-step trajectory) for Cy3-DOPE [TfR]. These results support the occurrences of hop diffusion in the intact PM and simple-Brownian diffusion in the blebbed PM.

The duration between two consecutive TILD events represents the dwell lifetime within a compartment. The distributions of the dwell lifetimes are shown in Fig. 5 A. They could be fitted with single exponential decay functions, with lifetimes of 9.2 ± 0.34 ms for Cy3-DOPE and 23 ± 1.5 ms for TfR. The exponential shape of this distribution is consistent with the prediction by hop diffusion theory developed here (in the subsection “Expected distribution of the dwell lifetimes: development of the hop diffusion theory” in the caption to Fig. S5). The dwell lifetime for TfR is longer than that for Cy3-DOPE by a factor of 2.5, probably because the fence and picket effects both act on TfR, whereas only the picket effect works on Cy3-DOPE.

**Figure 5.**
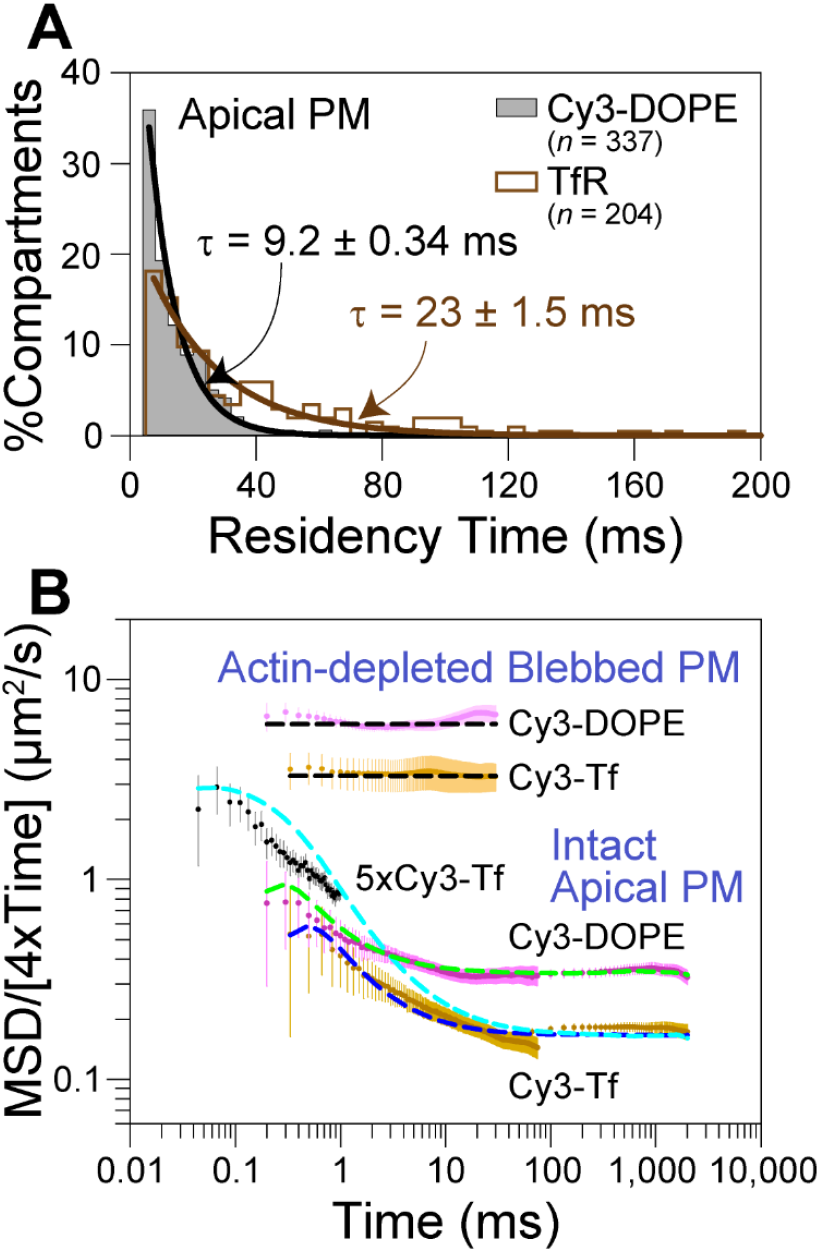
Analyses of the residency time of each molecule within each compartment (A) and the diffusion anomalies based on log(MSD/time) vs. log(time) in the time range over five orders of magnitude (B) were made possible by the development of ultrafast single fluorescent-molecule imaging, providing new insights into the hop diffusion of membrane molecules in the PM. (**A**) The distributions of the residency times within a compartment for Cy3-DOPE and TfR, determined by the TILD analysis, with the best-fit exponential curves. (**B**) An anomaly analysis of single-molecule trajectories based on the plot of log(MSD/time) vs. log(time) (see the main text). Using the data obtained at time resolutions of 0.022 ms (45 kHz; only for TfR labeled with 5xCy3-Tf), 0.1 ms (10 kHz; for Cy3-DOPE), 0.167 ms (6 kHz; for TfR labeled with Cy3-Tf), and 33 ms (30 Hz; for Cy3-DOPE and TfR labeled with Cy3-Tf), the mean values of log(MSD/time) averaged over all trajectories were obtained and plotted as a function of log(time). The results of TfR using Cy3-Tf and 5xCy3-Tf were different in the time ranges shorter than 1 ms, due to the differences in the observation frame rates (6 and 45 kHz, respectively). During 0.167 ms, which is the frame time in 6-kHz observations, TfR still collides with the compartment boundaries, but this occurs much less often when the frame time is 0.022 ms (frame time in 45-kHz observations). Therefore, in shorter time ranges, the results obtained at 45 kHz (using 5xCy3-Tf) are better (see the simulation results shown by the dashed cyan curve). For the same reason, the 10-kHz data using Cy3-DOPE show that due to the insufficient time resolution, the pure simple-Brownian diffusion within a compartment could not be measured even at this frame rate. Dashed curves represent the results of the Monte Carlo simulations, resembling the experimental data (see **Materials and methods** for the simulation parameters). Note that the phospholipid probes are located in the PM outer leaflet, and yet they undergo hop diffusion. This is probably because, as proposed previously (Fujiwara et al., 2002), the transmembrane proteins anchored to and aligned along the actin mesh (pickets; see Fig. 3 A) form the diffusion barrier in both the outer and inner leaflets of the PM. The picket effect is due not only to the steric hindrance of the picket proteins, but also to the hydrodynamic-friction-like effect from the surface of the immobilized picket proteins on the surrounding medium (Fujiwara et al., 2002). Monte Carlo simulations showed that 20 – 30% occupancy of the compartment boundary by the immobile picket proteins (bound to the actin fence) is sufficient to cause confined + hop diffusion of the phospholipids in the PM outer leaflet (Fujiwara et al., 2002).

The *average* dwell lifetime within a compartment could be obtained from the median values of *L* and *D*_MACRO_ determined by the hop fitting of the MSD-*Δt* plot, using the equation *τ* = *L*^2^/4*D*_MACRO_ (Fig. 4, C **and** D; **Table 2**; theory in Fig. S5). This provided dwell lifetimes of 8.5 ms for Cy3-DOPE and 24 ms for TfR, which are in good agreement with the values obtained based on the dwell lifetime distributions determined by the TILD method (**Table 2**).

Previously, with gold probes, we obtained *L* from ultrafast single-gold-particle tracking, but needed to separately obtain *D*_MACRO_ using fluorescently-tagged molecules (by observing them at video rate) to obtain the average dwell lifetime within a compartment (Fujiwara et al., 2002, 2016; Murase et al., 2004). This is because gold probes induced crosslinking of the target molecules, elongating the dwell lifetime, and thus provided reduced *D*_MACRO_ values. Using the ultrafast single fluorescent-molecule imaging, we can directly determine the dwell lifetime of *each molecule* in *each compartment* (not just the averaged lifetime).

### An anomaly analysis provided further proof of the hop diffusion of Cy3-DOPE and TfR in the compartmentalized PM

Generally, the MSD can be expressed as a function of the time interval Δ*t*,

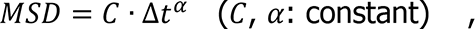

where 0 ≤ *α* ≤ 2 (*α* = 1 for simple-Brownian diffusion, 0 ≤ *α* < 1 for suppressed anomalous diffusion, 1 < *α* < 2 for super diffusion, and *α* = 2 for ballistic motion) (Saxton, 1994, 1996; Feder et al., 1996; Simson et al., 1998; Fujiwara et al., 2002). This equation can be rewritten as,

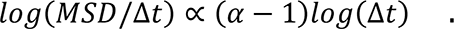

This relationship is plotted in Fig. 5 B. A very broad time range of ∼5 orders of magnitude (from 0.044 ms to 2 s) was covered, which was made possible by the development of the ultrafast single fluorescent-molecule imaging.

In this display, simple-Brownian type diffusion is represented by a flat line (since *α* = 1 in simple-Brownian cases, the slope *α* - 1 ∼ 0; i.e., no time dependence) and suppressed diffusion is characterized by negative slopes ([*α* - 1] < 0; i.e., *α* < 1). Namely, the level of the diffusion anomaly can be parameterized by the parameter *α* (Feder et al., 1996; Simson et al., 1998). In the actin-depleted blebbed PM, both Cy3-DOPE and TfR exhibited flat lines in the entire time range of 0.3 – 30 ms, unequivocally showing that they both undergo simple-Brownian diffusion in the blebbed PM.

On the contrary, in the intact apical PM, the plots for Cy3-DOPE and TfR (Cy3-Tf and 5xCy3-Tf) exhibited suppressed diffusion (negative slopes) in the time ranges of 0.5 – 30 and 0.13 – 70 ms, respectively, indicating that the suppressed diffusion of Cy3-DOPE and TfR was detectable in these time ranges. Meanwhile, in longer and shorter time ranges, the plots asymptotically approached flattened plots, consistent with simple-Brownian diffusion in these time regimes.

These results can readily be explained by the hop diffusion model in the compartmentalized PM for both Cy3-DOPE and TfR. In the shorter time regime (<0.5 and 0.13 ms, respectively), Cy3-DOPE and TfR collide with the compartment boundaries less frequently, approaching simple-Brownian diffusion and thus providing the diffusion coefficient of *D*_micro_ within a compartment (although this is still underestimated due to the collisions with the compartment boundaries. However, in this time regime, translocation across the compartment boundaries hardly occurs). In the longer time regime (>30 and 70 ms, respectively), the diffusion of Cy3-DOPE and TfR represents that among compartments, because the detailed shorter-term behaviors are averaged out, and their movements thus resemble simple-Brownian diffusion again, with a diffusion coefficient of *D*_MACRO_. In the time ranges of 0.5 – 30 and 0.13 – 70 ms for Cy3-DOPE and TfR (Cy3-Tf and 5xCy3-Tf), respectively, since the effect of confinement within a compartment becomes increasingly evident with a lengthening of the observed time window until occasional hops across the compartment boundaries start alleviating this confinement, the MSD/(4*t*), which is the y-axis in this plot, is expected to decrease from *D*_micro_ until it reaches *D*_MACRO_.

The dashed curves in Fig. 5 B represent the results of the Monte Carlo simulations that resemble the experimental data. The cyan curve would be the best for representing the diffusion of TfR in the apical PM. In the longer time regime (>70 ms), it approached the experimental result for Cy3-Tf at 30 Hz, while in the shorter time regime (<0.8 ms), it approached the experimental data for 5xCy3-Tf at 45 kHz, and at its shorter limit, it approached the flat line of the Cy3-Tf observed in the blebbed PM.

### Ultrafast PALM, making PALM practical for live-cell observations

PALM imaging (and all other single-molecule localization microscopy methods) of live cells requires a good balance between the reasonable view-field size and the time resolution (Jones et al., 2011; Huang et al., 2013; see the bottom six rows in **Table 1**). The newly developed camera system greatly improved these parameters. Using one of the best photoconvertible fluorescent proteins available for PALM, mEos3.2 (Zhang et al., 2012), the developed camera system allowed us to employ a PALM data acquisition view-field of 640 x 640 pixels (35.3 x 35.3 µm^2^), with PALM time resolutions quite useful for observing morphological changes of subcellular structures of 0.33 - 10 s (1-kHz data acquisition for 330 - 10,000 frames), with a single-molecule localization precision of 29 nm. Therefore, the time-dependent changes of meso-scale (on the order of several tens to several hundreds nm) structures could be observed in the view-field of a whole live cell in the time scale of 0.33 - 10 s, as explained in the following.

The data acquisition frame rate, and thus the PALM time resolution, could not be increased to 10 kHz, as done for single Cy3 molecule tracking. This is due to the low throughput of the available fluorescent probes. Even mEos3.2 (fused to the C-terminus of mouse caveolin-1 and expressed in T24 cells) cannot be photobleached faster than 1.2 ms (cannot be excited at higher rates) at 561-nm excitation laser intensities up to 100 µW/µm^2^ (Fig. 6 A-a). Therefore, a frame rate of 1 kHz is about the fastest useful frame rate, as long as mEos3.2 is used as the PALM probe. However, even this is 33 times faster than video rate, and thus shortened the PALM data acquisition time by a factor of ∼33 (as compared with the video-rate acquisition method) to 0.33 – 10 s (1 ms/frame x 333 – 10,000 frames).

**Figure 6.**
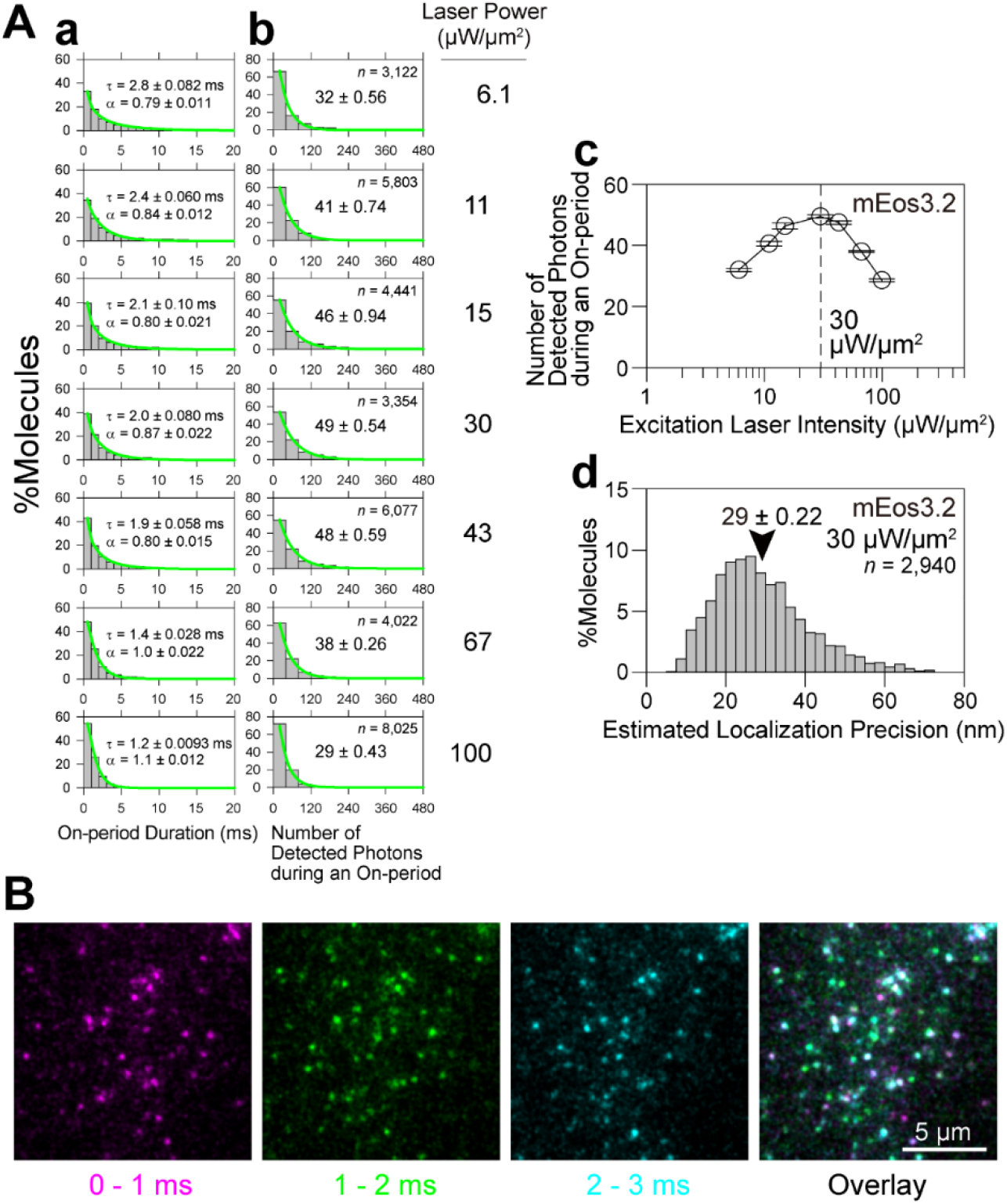
Optimizing the excitation laser power density and the data acquisition frame rate for PALM using mEos3.2 in living cells. The mEos3.2 fused to mouse caveolin-1, expressed in the basal PM of live T24 cells, was imaged. The photoconverted mEos3.2 molecules were observed with an integration time of 1 ms (1-kHz frame rate), using a 561-nm excitation laser with various power densities on the sample plane with TIR illumination. For details, see **Materials and methods** (Ultrafast PALM and its application to live-cell observations). (**A**) **The mean number of photons emitted from an mEos3.2 molecule during an on-period is maximized at a laser power density of 30 µW/µm^2^ at the sample plane (49 ± 0.54 photons), providing a single-molecule localization precision of 29 ± 0.22 nm, with an on-period of ∼2 ms.** (**a**) Histograms of consecutive fluorescent on-periods (with a gap closing of 1 frame) obtained at various laser power densities (indicated on the right of the boxes in **b**). They could be fitted by stretched exponential functions

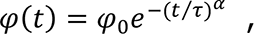

where *φ*_o_ is the prefactor, *α* is the stretching exponent, and *τ* is the time constant (Morimatsu et al., 2007) (mean ± SEM; SEM was determined as a 68.3% confidence limit for the fitting; the numbers of spots observed are the same as those indicated in the boxes in **b**). With an increase of the excitation laser power density, the on-period (*τ*) was reduced while the function became less stretched (the exponent *α* of ∼1 was recovered). (**b**) Distributions of the numbers of detected photons during an on-period. The histograms could be fitted with single exponential decay functions, with decay constants providing the mean numbers of detected photons during an on-period (the SEM was given as a 68.3% confidence limit for the fitting), which became maximal at a laser power density of 30 µW/µm^2^ at the sample plane (49 ± 0.54 photons). *n* = number of observed spots. (**c**) Summary plot for the results in (**b**). A laser power density of 30 µW/µm^2^ at the sample plane maximizes the number of photons emitted from mEos3.2 during an on-period (the measured parameter was the number of “detected” photons). (**d**) The mean localization precision for mEos3.2 in the basal PM was 29 ± 0.22 (SEM) nm, under the optimized laser excitation conditions of 30 µW/µm^2^. The localization precision was estimated from the theoretical equation derived by Mortensen *et al.* (Mortensen et al., 2010) with an “excess noise” factor (*F*) of 1.2 determined for the developed camera system (Fig. S2). (**B**) Single-frame images of mEos3.2-caveolin molecules in the basal PM acquired every 1 ms (1 kHz), using an observation laser power density of 30 µW/µm^2^. While many spots exist in a single image, some spots appear in two or three images (spot colors are changed every 1 ms). This result is consistent with the finding that data acquisition at 1 kHz is nearly optimal for PALM imaging of mEos3.2 molecules.

A single mEos3.2 molecule emits the largest number of photons during its on-time at a laser intensity of 30 µW/µm^2^ at the sample plane (Fig. 6 A-b **and** c; Alexa647 exhibits similar power dependence as reported by Lin et al. [2015]), providing the best single-molecule localization precision of 29 nm (Fig. 6 A-d). Therefore, we employed the laser intensity of 30 µW/µm^2^ and a camera frame rate of 1 kHz throughout this study (Fig. 6 B). With the future advent of photoconvertible molecules that can emit photons faster and photobleach faster, we could use 10 (or 30) kHz, rather than 1 kHz, for the data acquisition, which will allow us to obtain PALM images at least 10 times faster than the rate achieved here using the developed camera system.

The single-molecule localization precision using the more prevalent sCMOS was 22 nm (256 x 256 pixels with a data acquisition time of 0.5 s; 0.6 kHz for 300 frames) (Huang et al., 2013), which is better by a factor of 1.3 compared with the developed camera system (29 nm; **Table 1**). This is due to the lower quantum efficiency of the photosensor of the image intensifier used here (∼40%), as compared with that of sCMOS sensors (70–80%) ([70/40]^1/2^ ≈1.3).

Typical time-series of expanded PALM images of caveolin-1-mEos3.2 in a live T24 cell, generated in a reconstruction time window of 1 s (data acquisition at 1 kHz x 1,000 frames; 10 x 10 µm^2^ in 181 x 181 pixels), are shown in Fig. 7 A. Considering the localization precision of 29 nm, the obtained caveolin PALM images were consistent with the size of a caveola (60 ∼ 80 nm in diameter; Parton and del Pozo, 2013). The whole caveola sometimes moves together on the PM (compare Fig. 7 A with Fig. 7 B), and when the caveola migration occurs, it can suddenly move for ∼100 nm or so in <0.33 s, suggesting the movement from one actin-induced compartment to an adjacent one.

**Figure 7.**
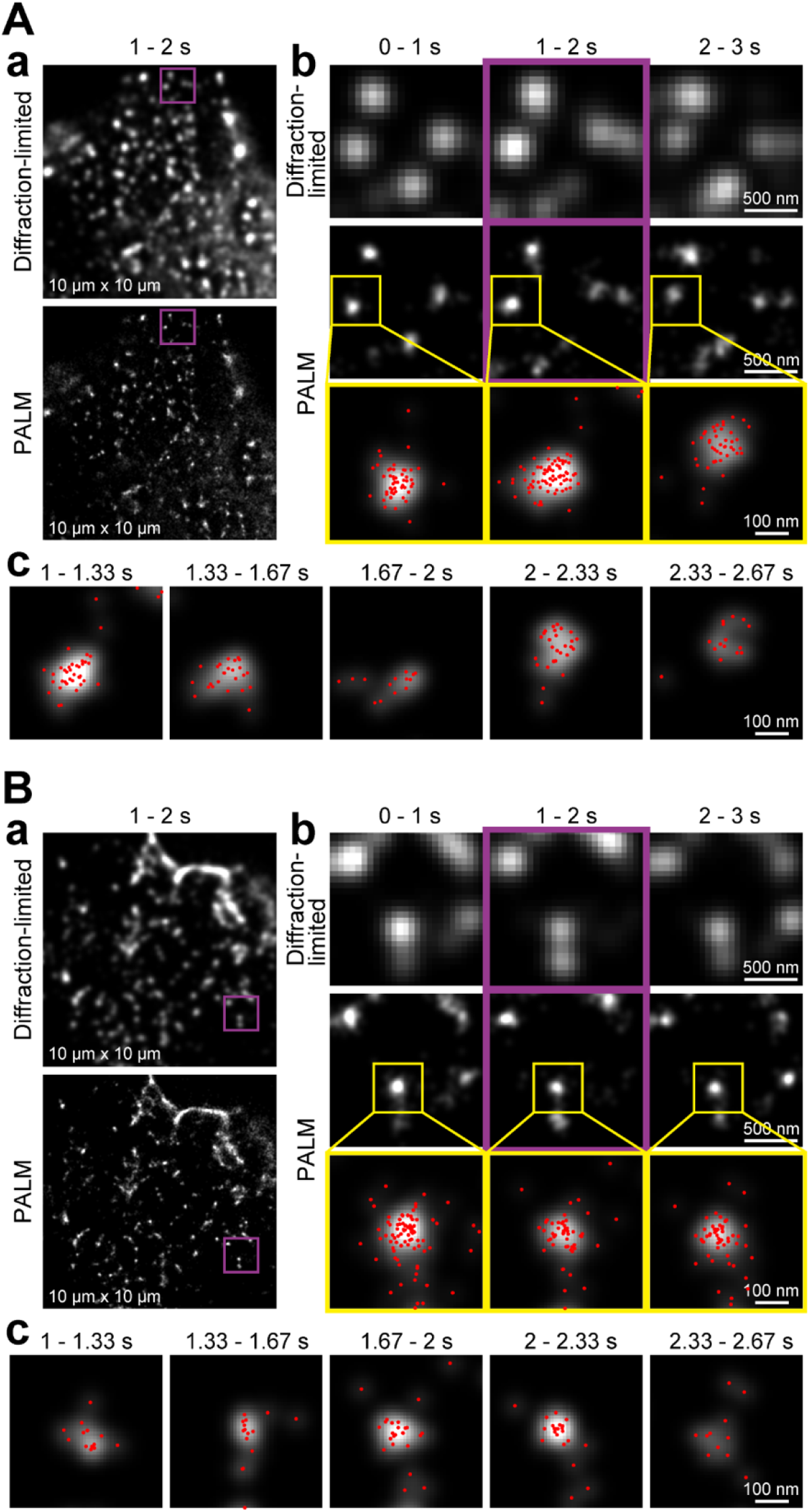
Ultrafast PALM reveals the shape changes, migrations, and formation/disappearance of caveolae in 3 s in live cells. Caveolin-1-mEos3.2 expressed in the basal PM of T24 cells was imaged in live cells (Panel **A**) and fixed cells (Panel **B**), using identical recording conditions. Data acquisitions were performed at a rate of 1 kHz for 3 s (3,000 frames), and PALM images were reconstructed using the data acquired for every 0.33 s (= 333 frames; nine reconstructed images) and every 1 s (= 1,000 frames; three reconstructed images). (**a**) Diffraction-limited (**top**) and PALM (**bottom**) images of caveolae in 10 x 10-µm^2^ observation areas (the same field of view) with a data acquisition period of 1 s. (**b**) Enlarged images of the purple-square regions in (**a**), showing time-dependent changes (every 1 s). The images in the **middle column** (top and middle rows) are the same images in the purple-square regions in (**a**), for the data acquisition between 1 and 2 s. The regions surrounded by yellow squares in the **middle row** are magnified in the **bottom row**, and single mEos3.2 molecule localizations determined within each 1-s period are indicated by red dots. (**c**) The PALM image reconstruction performed every 0.33 s for the same caveola shown in the **bottom row** in (**b**).

The cost of the increased image acquisition rates is the spatial resolution of the PALM images. The spatial resolutions of the PALM images shown in Fig. 7 A-a and B-a were 75 and 77 nm, as evaluated by a parameter-free decorrelation analysis (Descloux et al., 2019), whereas the conventional sCMOS would provide a spatial resolution of >58 [75/1.3] nm (from quantum efficiencies) (Lelek et al., 2021).

### Ultrafast PALM for the entire live cell at a 75-nm spatial resolution

Although the PALM data acquisition rate was limited to 1 kHz using mEos3.2, this camera system allows us to use the data acquisition frame size of 640 x 640 pixels (35.3 x 35.3 µm^2^ with the pixel size of 55.1 nm) at 1 kHz. The PALM image of mEos3.2 fused to human paxillin, a focal adhesion structural protein, expressed on the basal PM, with the acquisition of the images of 640 x 640 pixels at 1 kHz for 10,000 frames (10 s) is shown in Fig. 8. Almost the entire live cell is seen in the view-field with a spatial resolution of 75 nm (Descloux et al., 2019), exhibiting the distributions of focal adhesions in the cell, and at the same time, the nano-scale fine structures including paxillin within each focal adhesion could be visualized. This image size was the largest ever for any PALM data obtained in the time scale of 0.33 – 10 s (1 ms/frame x 333 – 10,000 frames), the time range useful for observing live cells (for the comparison with previous results, see **Table 1**). The presence of paxillin islands is consistent with previous reports (Shroff et al., 2008; Patla et al., 2010; Rossier et al., 2012; Levet et al., 2015; Changede and Sheetz, 2017; Spiess et al., 2018; Orré et al., 2021). For our further analysis of the focal adhesion simultaneously using single-molecule imaging and PALM, please see the companion paper (Fujiwara et al., 2021).

**Figure 8.**
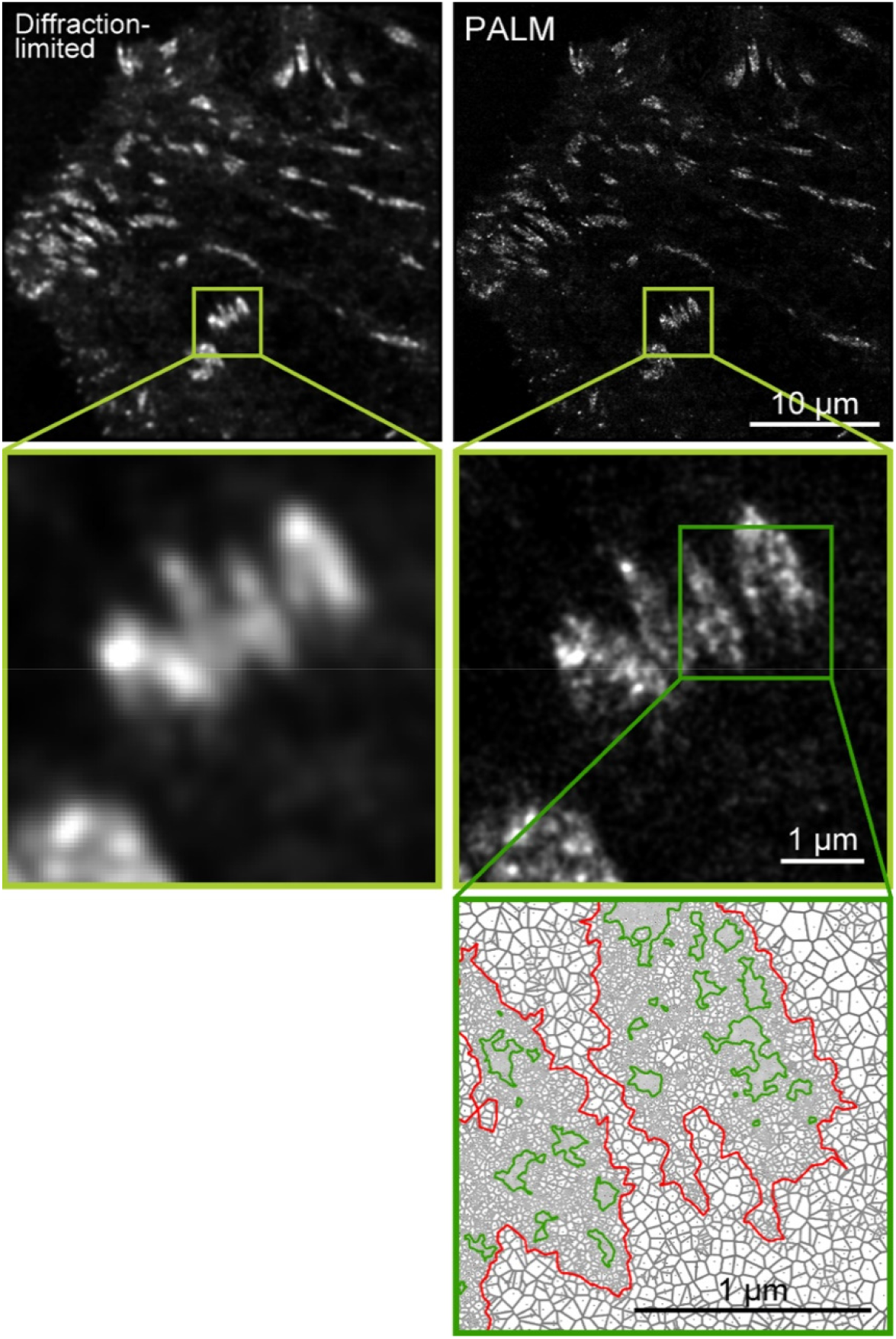
Representative ultrafast PALM superresolution image of focal adhesions in a field of view encompassing almost the entire live cell. The paxillin-mEos3.2 expressed on the basal PM of a live T24 cell, visualized by ultrafast PALM with the data acquisition of a very large 640 x 640 pixel field of view (35.3 x 35.3 µm^2^) at 1 kHz for 10 s (10,000 frames). The diffraction-limited (**left-top**) and PALM (**right-top**; 3,526 x 3,526 pixels with the pixel size of 10 nm) images were reconstructed. The areas indicated by the light green squares in the **top** images are expanded in the **middle** images. The SR-Tesseler software based on Voronoï polygons (Levet et al., 2015) was applied to the locations of mEos3.2-paxillin in the **right-middle** image, and the detected contours of FAs (red) and paxillin-protein islands (dark green) in the dark-green square are expanded in the **bottom** image.

We emphasize here that, with the present camera development, the instrument is no longer the limitation for the speed of PALM imaging. Instead, the probe is now the limitation. With the future development of photoconvertible molecules that could emit photons faster and photobleach *faster*, we would be able to obtain PALM images 10 times more quickly than we can now using the developed instrument (by employing the data acquisition rate of 10 kHz rather than 1 kHz for 640 x 640 pixels), enabling PALM imaging at video rate (for the acquisition of 300 frames).

## Discussion

The ultrahigh-speed camera system developed here has enabled the fastest single fluorescent-molecule imaging ever performed. This would also be the ultimate rate possible with the currently available fluorescent molecules; e.g., 0.1-ms resolution with a 20-nm localization precision for single Cy3 molecules, using a saturating TIR (standard oblique-angle) illumination laser intensity of 79 µW/µm^2^, for a frame size of 256 x 256 pixels (14 x 14 µm^2^). Under these conditions, approximately 1.6% of the single Cy3 molecules could be tracked for periods longer than 100 frames (10 ms; corresponding to 3.3 s at video rate) (for 39-nm precision, the laser intensity is 23 µW/µm^2^, and 14% of single Cy3 molecules could be tracked for the same period). When better fluorophores (faster emission and slower photobleaching) are developed, the time resolution for single-molecule imaging could be enhanced to 22 µs or more (for a frame size of 7.5 x 3 µm^2^ in 128 x 64 pixels), without affecting other parameters (Fig. 1, **Table 1**).

For tracking just one molecule at a time, MINFLUX has recently been developed, and its performance on the glass is superb (Balzarotti et al., 2017; Eilers et al., 2018; Schmidt et al., 2021). It has achieved single-molecule localization precisions of 2.4-nm and <20 nm with 400- and 117-µs temporal resolutions, respectively. However, it can track only one molecule at a time (but could be expanded to track a few isolated molecules at a time). In contrast, the camera-based single molecule imaging allows the observations of several tens to hundreds of molecules simultaneously in a field of view (like an entire cell). The camera-based method can thus be used to investigate the interactions and assemblies of molecules in live cells and study the different behaviors of molecules simultaneously in various parts of the cell, which is impossible with one molecule/cell technologies.

In live *E. coli*, MINFLUX provided a single-molecule localization precision of 48 nm with a 125-µs time resolution, under the conditions of observing the same molecule 742 times (Balzarotti et al., 2017). This is quite comparable to the results we obtained using the camera-based method, which allows us to simultaneously observe many single molecules at single-molecule localization precisions of 20 and 34 nm (27 and 55 nm) at time resolutions of 100 and 33 µs using TIR (oblique-angle) illumination (**Table 1**). However, practically speaking, the same single molecules could be observed for 100 – 300 frames (10 – 30 ms at 10 kHz; localizing them 100 − 300 times), as compared with localizing one molecule 742 times with MINFLUX. Therefore, the ultrafast camera-based single-molecule imaging and MINFLUX provide complementary methods for examining the very fast dynamics of membrane molecules in living cells: MINFLUX is useful for tracking a single molecule for longer periods, whereas the ultrafast camera-based single-molecule imaging provides a method for observing many molecules at once to investigate their interactions and the location-dependent behaviors of single molecules in a cell. Single molecules labeled with two or three different colors could also be investigated readily using the two or three cameras, respectively.

The ultrafast camera system developed here was tested by examining whether it can detect the hop diffusion of Cy3-tagged DOPE and TfR in the PM of live cells, as expected from the previous observations using gold-tagged molecules (Fujiwara et al., 2002, 2016; Murase et al., 2004). We found that it can indeed detect the hop diffusion of Cy3-tagged molecules (Figs. 3 - 5; **Table 2**) and the same compartment size as found by ultrafast gold-tracking (∼100 nm in the PM of T24 cells; Fig. 4 D; **Table 2**).

Ultrafast single fluorescent-molecule tracking has brought forth new knowledge about the hop diffusion of membrane molecules. First, the distribution of the dwell lifetimes in each compartment (not the overall average lifetime) was obtained. By the theory developed here, we found that the distribution of the dwell lifetimes indeed follows single exponential functions (Fig. 5 A). Second, ultrafast single fluorescent-molecule tracking allowed an anomaly analysis of a very broad time range of ∼5 orders of magnitude (from 0.044 ms to 2 s) for determining the diffusion properties of Cy3-tagged molecules. The anomaly analysis confirmed the hop diffusion of Cy3-DOPE and TfR in the PM and their simple-Brownian diffusion in the actin-depleted blebbed PM (Fig. 5 B).

Meanwhile, the simple-Brownian diffusion of both Cy3-DOPE and TfR in the actin-depleted blebbed PM indicates that actin filaments (and their associated transmembrane picket proteins) are responsible for compartmentalizing the PM (Fig. 3 A). This conclusion is consistent with previous observations, including the changes of the compartment sizes after the treatments with actin-modulating chemicals, latrunculin, cytochalasin D, and jasplakinolide (Fujiwara et al., 2002; Fujiwara et al., 2016; Murase et al., 2004), the equality of the compartment sizes determined by the lipid diffusion data and the actin mesh sizes on the apical PM cytoplasmic surface determined by electron tomography (Morone et al., 2006) and super-resolution microscopy (Xia et al., 2019), and the results of single-molecule optical trapping (Sako and Kusumi, 1995; Sako et al., 1998). New data about the hop diffusion of membrane molecules in the basal PM and even in the focal adhesions in the basal PM are reported in the companion paper.

We believe that the PM compartmentalization is fundamentally important for understanding the PM function. It would allow the cells to locally enhance phosphorylation at particular places on the PM, by confining the kinases in the compartment (due to higher local concentrations of kinases than phosphatases, which are more prevalent in the cytoplasm) (Kalay et al., 2012; Kusumi et al., 2012a), to localize stimulation-induced stabilized raft domains (Kusumi et al., 2004; Kusumi et al., 2011, 2012a, b; Suzuki et al., 2012; Kusumi et al., 2020), and to suppress the diffusion of engaged receptor oligomers (oligomerization-induced trapping) (Iino et al., 2001). However, due to the difficulties in using large gold probes and visualizing them at high frame rates, very few scientists have directly investigated and detected hop diffusion in the PM. With the developed ultrafast camera system with single fluorescent-molecule sensitivities, PM compartmentalization and hop/confined diffusion of membrane molecules could be studied quite readily by many more scientists, and thus their investigations could progress much faster.

This camera system now allows us to perform the fastest PALM, with a data acquisition rate of 1 kHz in the largest view-field (640 x 640 pixels; 35.3 x 35.3 µm^2^ with the pixel size of 55.1 nm) (Figs. 7 and 8). The data acquisition of 333 – 10,000 frames takes only 0.33 – 10 s, enabling live-cell PALM for certain structures such as caveolae and FA (Figs. 7 and 8). Although a single-molecule localization precision of 29 nm (for mEos3.2 at 1 kHz) is worse as compared with 2 – 25 nm (Betzig et al., 2006), 12 – 29 nm (Shroff et al., 2007), and 12 nm (Zhang et al., 2012) using EM-CCD and sCMOS sensors, the gain in time resolution and the applicability to observations of live-cell events could compensate for the spatial resolution deterioration. The ultrafast PALM also affords the possibility of performing PALM imaging of live cells at video rate, when new fluorescent dyes and proteins that can be photoconverted or photoactivated more efficiently become available.

Meanwhile, ultrafast PALM and ultrafast single fluorescent-molecule imaging could be performed simultaneously, as described in the companion paper (Fujiwara et al., 2021). Namely, the developed ultrafast camera system now allows us to observe subcellular structures in/on the PM with nano-scale precisions, and at the same time, to detect how individual single molecules interact with or behave in these structures. Therefore, we conclude that the ultrafast camera system developed in the present work will become an especially powerful tool for cell biology.

## Materials and methods

### Ultrahigh-speed intensified CMOS camera system: design and operation

*Detailed description of the camera system* (Fig. 1; Figs. S1-S3; **Table 1**) See the schematic diagram of the developed camera system shown in Fig. 1 A. The new ultrahigh-speed camera system consists of the following three major components.

(a) A Hamamatsu image intensifier (V8070U-74), comprising a third-generation GaAsP photocathode with a quantum efficiency of 40% at 570 nm (Fig. 1 A-a-α), a three-stage microchannel plate allowing maximal gain over 10^6^, in which a non-linear noise increase was virtually undetectable (Fig. 1 A-a-β), and a P46 phosphor screen with a decay time of approximately 0.3 µs from 90% to 10% (Fig. 1 A-a-γ).

(b) A CMOS sensor (unless otherwise specified, we report the results obtained with the sensor developed for a Photron 1024PCI camera, but for some tests, we additionally report the results obtained with the newer sensor developed for a Photron SA1 camera), with a global shutter exposure was used (**Table 1**). The 1024PCI (SA1) sensor is composed of 1,024 x 1,024 pixels with a 17 x 17-µm (20 x 20-µm) unit pixel size, and operates up to 1 kHz or every 1 ms (5.4 kHz or every 0.19 ms) for the full-frame readout. The 1024PCI sensor can be operated at 10 kHz (0.1-ms resolution) with a frame size of 256 x 256 pixels, at 45 kHz (0.022-ms resolution) with a frame size of 128 x 64 pixels, and at 110 kHz (0.009-ms resolution) with a frame size of 128 x 16 pixels. The SA1 sensor can also be operated at 16 kHz (0.063-ms resolution) with a frame size of 512 x 512 pixels, and at 45 kHz (0.022-ms resolution) with a frame size of 256 x 256 pixels (**Table 1**).

However, the actual observation frame rates for single-molecule tracking with reasonable single-molecule localization precisions (say ≤50 nm) were limited to 10 - 30 kHz. This is because the number of fluorescent photons that can be emitted from a single fluorophore during < 0.033 - 0.1 ms (the integration time of a camera frame); i.e., the frequency that a single fluorophore can be excited, is limited, as quantitatively described in “Estimation of the number of photons that can be emitted by a single Cy3 molecule during 0.1 ms: triplet bottleneck saturation” in the **Materials and methods** (also see Fig. 1, E **and** F; Fig. S3). Namely, with the development of our ultrafast camera system, the instrument is no longer the limitation for achieving higher time resolutions (faster than 10 - 30 kHz) for single fluorescent-molecule imaging, and now the availability of fluorescent dyes has become the limitation. With the future development of new dyes with faster excitation and emission, without making a transition to the triplet state, observations even at 110 kHz would be possible. Indeed, we showed that when an average (mean) of five Cy3 molecules were attached to a single Tf protein (5xCy3-Tf), it can be observed at a rate of 45 kHz (Fig. 1 I; Figs. S2 A and S3; **Table 1**).

The readout noise, including the dark noise, of the 1024PCI and SA1 sensors at the full-speed readout at 21°C was 37 root-mean-square electrons/pixel, as measured by the manufacturer (Photron), and provided dynamic ranges of ∼800 and ∼1,200, respectively. The quantum efficiency of the CMOS sensor employed here was 40% (for both sensors), but since the CMOS sensor is placed after the image intensifier, the lower quantum efficiency of the CMOS sensor is not a critical issue in the developed camera system (the quantum efficiency of the image intensifier photocathode matters).

(c) A straight (1:1) optical-fiber bundle directly adhered to the phosphor screen of the image intensifier on one side (Fig. 1 A-a-γ, input side) and to the CMOS sensor on the other side (Fig. 1 A-b, output side). This enhanced the signal reaching the CMOS sensor by a factor of 5-10, as compared with the lens coupling used for our standard camera system employed for single fluorescent-molecule imaging (Suzuki et al., 2012; Kinoshita et al., 2017).

The electrons generated at the CMOS sensor were transferred, amplified (Fig. 1 A-d), and digitized (Fig. 1 A-e) in 64 and 8 parallel paths, respectively, and then transferred to the host PC.

#### Basic design concept of the camera system

##### (A) The use of a CMOS sensor rather than an sCMOS sensor or an EM-CCD sensor

To achieve ultrahigh-speed single fluorescent-molecule imaging, we used a CMOS sensor rather than an sCMOS sensor or an EM-CCD sensor, which are broadly employed in fluorescence microscopy. This is because the maximum frame rates attainable with the sCMOS sensors and EM-CCD sensors (typically 1.2 – 4 ms/frame for a 256 x 256-pixel image) are at least 10 times slower than those of the CMOS sensors employed here (0.1 and 0.015 ms/frame for a 256 x 256-pixel image, using the Photron 1024PCI sensor and SA1 sensor, respectively; no binning). To avoid image distortions and achieve higher rates, we employed CMOS sensors with a global shutter (most sCMOS cameras employ rolling shutters; although some newer sCMOS cameras are equipped with global shutters, their frame rates are slower than those when rolling shutters are employed).

##### (B) The use of an image intensifier (also see the beginning of **Results** in the main text)

An obvious problem when using the CMOS camera is its high readout noise (30∼40 vs. <5 root-mean-square electrons/pixel/frame for CMOS vs. sCMOS; the term “readout noise” used in this report always includes the dark noise of the sensor). As described in the previous section, the readout noise of the 1024PCI and SA1 sensors employed in this study was 37 root-mean-square electrons/pixel/frame (at the full-speed readout at 21°C). Meanwhile, the background noise signal (*b* in the equation in the caption to Fig. S2) was 0.035 ± 0.058 detected photons/pixel/frame on the glass using the oblique illumination (mean ± SD; in the case of TIR illumination, *b* = 0.038 ± 0.059 detected photons/pixel/frame) and 0.068 ± 0.17 detected photons/pixel/frame on the cellular apical PM (oblique illumination). Therefore, the background noise signal is much smaller than the readout noise, as expected (i.e., *N*_p_ << *N*r as described in the subsection “Basic conceptual strategy to address the large readout noise of the CMOS sensor” in the main text). As shown in Eq. 3 there, *G* should be much greater than *N*r/*N*_p_, i.e., *G*>>*N*r/*N*_p_. Therefore, *G*>>37/0.035 (or 0.068) or *G*>>1,100 (or 540) for single fluorescent molecules on the glass (apical PM). As described in the following paragraphs, by also considering *S*_p_, we found a total amplification gain *G* of 8,100 useful for single fluorescent-molecule imaging.

Next, we address the signals from the fluorophores. Consider our standard test conditions for observing a single Cy3 molecule, using a frame rate of 10 kHz and *oblique* illumination for excitation at 23 µW/µm^2^ (i.e., instrumentally less optimal as compared with the TIR illumination, but necessary for observing molecules in the apical PM as well as in the basal PM). Under these conditions, we obtained 34 ± 2.4 detected photons/molecule/frame (Fig. 1 **E top**; including a quantum efficiency of the image intensifier photocathode of 0.4 at 570 nm; i.e., the number of photons that arrived at the photocathode of the intensifier was 85 ± 6.0 photons/molecule/frame), which realized a single Cy3-molecule localization precision of 38 ± 1.7 nm on the coverslip (Fig. 1 F **top**). Namely, an average (total number of) of 34 ± 2.4 photons is detected on the CMOS sensor in the two-dimensional Gaussian image of a standard deviation (SD; radius) of 123 ± 1.1 nm (2.2 ± 0.020 pixels x 55.1 nm/pixel) on the sample plane. (This image size was determined by the Gaussian fitting of each image for 50 Cy3 molecules immobilized on the glass under the TIR illumination at 79 µW/µm^2^; this higher illumination was employed to determine the Gaussian image size more precisely than that obtainable under the oblique illumination at 23 µW/µm^2^, by generating approximately 3 times more detected photons; see Fig. 1 E **top**; image magnification by a factor of 300 [150 x 2]).

Consider how the 34 photons are distributed in the pixelized Gaussian image; i.e., the 2-dimensional intensity profile (point-spread function) of a single-molecule emitter pixelized by the CMOS sensor. The expected number of photons in the pixel located at the integer position (*m*, *n*), *I*(*m*, *n*), is given as (Huang et al., 2011),

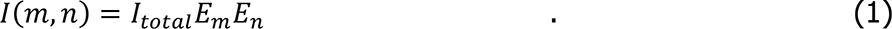

Here *I*_total_ is the total number of detected photons in the image; i.e., 34 photons, and *E*_m_ and *E*_n_ are

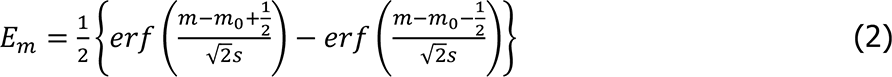

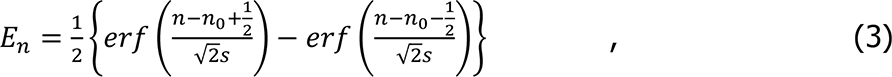

where *erf*represents a Gaussian error function, (*m*_o_, *n*_o_) is the sub-pixel emitter position, and s is the SD of the Gaussian spot profile; i.e., 2.2 ± 0.020 pixels. Using these equations, the peak pixel intensity *I*(0,0) (the number of detected photons at (*m*, *n*) = (0,0)) when a single-molecule emitter is located exactly at the center of the pixel (0,0) is estimated to be only 1.1 photons.

Under such low signal conditions, in which the pixel intensities within the Gaussian spot profile fluctuate frame by frame at the level of single photons, the photoelectrons emitted at the photocathode of the image intensifier should be amplified until the readout noise of 37 root-mean-square electrons/pixel/frame, which also fluctuates spatiotemporarily, becomes negligible. To satisfy this condition, the photon amplification gain of the image intensifier of 33,200 (the ratio of the number of photons emitted from the phosphor screen of the image intensifier vs. the number of detected photons) was used throughout this study (see the following estimates for details). The image intensifier was coupled to the CMOS sensor by an optical fiber bundle. Therefore, due to the 39% loss (61% coupling efficiency) by the optical fiber bundle and the quantum efficiency of the CMOS sensor of 40%, the overall electron amplification (after photon detection) was a factor of 8,100 (33,200 x 0.61 x 0.4). Since the quantum yield of the photocathode of the image intensifier was 0.4, the total amplification of the incident photon of this camera system was 3,240 (one photon that arrived on the photocathode of the image intensifier generated an average of 3,240 electrons in the CMOS chip).

The overall electron amplification of 8,100x (after photon detection) was selected based on the following two considerations. First, the probability that the electron signal at the CMOS sensor amplified from a single detected photon (at the photocathode of the image intensifier) becomes greater than the readout noise should be sufficiently high. As shown in Fig. S1 E, the probability of detecting a single photon (as normalized by the number of detected photons/frame at the saturating levels of electron amplification) was increased to 90.0% at an overall electron amplification of 8,100x.

Second, the upper limit of the overall electron amplification is given by the full-well capacity of 30,000 electrons for the individual pixel (for the 1024PCI sensor employed for most experiments in this study). Considering the stochastic fluctuation of the signal, we set the electron amplification of the image intensifier so that the average signal in the peak pixel of 1.1 electrons/pixel/frame was amplified to less than 1/3 of the full well capacity (<10,000 electrons); i.e., amplification by a factor of <9,090 (= 10,000/1.1). Combining these two considerations, we employed an overall electron amplification by a factor of 8,100 (33,200 at the image intensifier), 10% less than the factor 9,090, to further reduce the possibility of pixel intensity saturation. (For detecting dimers and larger oligomers, the image intensifier gain should be reduced, although the probability of missing monomers would increase slightly.)

##### (C) The design and conditions for visualizing and tracking single molecules for many frames (Fig. 2 C)

Note that these illumination and amplification conditions were selected so that we could track single molecules for many frames in the apical PM (e.g., for longer than 99 frames for 14% of the Cy3 molecules immobilized on the glass, under the conditions of 3-frame gap closing, as shown in Fig. 2 C). When only much shorter tracking is required (such as a single frame or in the case of PALM), much higher excitation laser power density could be employed for better single-molecule localization precisions. The best single-molecule localization precision we achieved with this camera system was 2.6 ± 0.099 nm (mean ± SEM), with a maximum number of detected photons/molecule/frame of 11,400 ± 700 (mean ± SEM; *n* = 50 Cy3 molecules; Fig. 1 G **and** H and Fig. S2 D), using a TIR illumination power density at 79 µW/µm^2^ and a signal integration time of 16.7 ms. This is because, under these conditions, most Cy3 molecules become photobleached within a single frame (16.7 ms; i.e., virtually all of the photons possibly obtained from single molecules are concentrated in a single frame) due to the use of a TIR illumination density of 79 µW/µm^2^, by which the photon emission rate from a single Cy3 molecule is saturated (Fig. 1 E).

### Estimation of the number of photons that can be emitted by a single Cy3 molecule during 0.1 ms: triplet bottleneck saturation (Fig. 1 E and F; Figs. S2 − S3)

In ultrahigh-speed single fluorescent-molecule imaging, the number of photons emitted from a single molecule during the duration of a single camera frame is the key limiting factor. This critically depends on how fast a molecule returns to the ground state from the excited state. In a conventional three-state model (ground, singlet-excited, and triplet-excited states), the maximum photon emission rate, *k*_em_ (photons/s), can be described as (Schmidt et al., 1995)

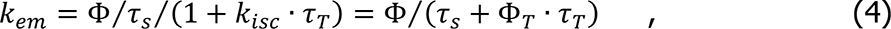

where *τ*_s_ and *τ*_T_ represent the lifetimes of the singlet- and triplet-excited states, respectively, *k*_isc_ (= Φ_T_⁄*τ*_s_) is the intersystem crossing rate, Φ is the fluorescence quantum yield, and Φ_T_ is the quantum yield for triplet formation. Namely, the maximum photon emission rate, *k*_em_, critically depends on the triplet yield and lifetime, and hence this phenomenon is known as “triplet bottleneck saturation”.

For Cy3, *τ*_s_ ≈ 1 ns, *τ*_T_ ≈ 1 µs (in a specimen equilibrated with atmospheric molecular oxygen, the triplet lifetime is dominated by the collision rate of molecular oxygen with the fluorophore, which is about 10^6^/s) (Kusumi et al., 1982), Φ = 0.15, and Φ_T_ = 0.003 were employed (Sanborn et al., 2007; Dempsey et al., 2011). These values provide a *k*_em_ of 3.8 x 10^7^ photons/s. Therefore, the maximum number of photons available to form an image of a single Cy3 molecule, during the integration duration of 0.1 ms for each frame, is approximately 3,800. Although this number would vary, depending on the spectroscopic parameters in the given environment, it provides a baseline for understanding such fast imaging. Assuming that 5-10% of the emitted photons will reach the intensifier photocathode (Rasnik et al., 2007; Gould et al., 2009), 190 - 380 photons will reach the photocathode of the image intensifier and be detected with a ∼40% quantum efficiency of the GaAsP photocathode at the Cy3 peak emission wavelength of 570 nm, providing the maximum number of photons detected during a 0.1-ms camera frame time of 80 - 150.

This estimate agrees well with the experimental results for evaluating the maximal photon emission rate of Cy3 by TIR illumination (here, since we are discussing saturation conditions, we focus on the results obtained by the TIR illumination rather than the oblique illumination) shown in Fig. 1 E **top** and Fig. S2 A **left** (the data displayed in Fig. S2 **A left** show the relationship of the single-molecule localization error with the number of detected photons/spot/frame obtained at a frame rate of 10 kHz, which was found to be consistent with the theory developed previously; Mortensen et al., 2010). In Fig. 1 E **and** F **(top)**, with an increase of the illumination intensity, both the number of detected photons/molecule/frame and the single-molecule localization precision were enhanced. However, the extents of the enhancement were saturated, because the number of photons that a single molecule can emit during the 0.1-ms frame time was limited. Under the near-saturation conditions at a laser power density of 79 µW/µm^2^ at the sample plane, the number of detected photons was ∼100 photons/frame (Fig. 1 E **top**). This confirmed that the experimental result is consistent with the estimate of the maximum photon emission rate based on the three-state model (ground, singlet-excited, and triplet-excited states; 80 – 150 photons during 0.1 ms) as described in the previous paragraph, indicating that the saturation of Cy3 photoemission can be explained by the “triplet bottleneck saturation”. Under the Cy3 saturation conditions, the improvement of the single-molecule localization precisions with an increase of the laser excitation intensity was limited to (saturated at) ∼20 nm for recordings at 10 kHz (Fig. 1 F **top** and Fig. S2 A **left**).

Among the eight dyes examined here, JF549, TMR, JF646, Atto647N, SeTau647, and Cy5 exhibited saturation more readily than Cy3 and Alexa555 (Fig. S2 B). Furthermore, the average numbers of detected photons from single molecules during the 0.1-ms frame time obtainable at saturation were smaller than those of Cy3 and Alexa555, and thus their single-molecule localization precisions at 10 kHz at saturation were worse than those of Cy3 and Alexa555 (Fig. S3). At saturation, Cy3 provided slightly better single-molecule localization precision at 10 kHz than Alexa555 (Fig. S3 B). Therefore, we primarily employed Cy3 throughout the remaining part of this report. Note that this result depended not only on the photophysical properties of individual dyes, but also on the wavelength dependence of the photocathode quantum yield of the image intensifier. The quantum yield is lower (∼0.2) for the near-IR dyes, JF646, Atto647N, SeTau647, and Cy5, as compared with that (∼0.4) for the Cy3, Alexa555, JF549, and TMR dyes.

### Determination of the number of detected photons/molecule/frame (*N*) (Fig. 1, E and G; Fig. 2, A and B; Fig. 6, A-b and c; Figs. S2−S3)

All of the fluorescent dye molecules used for determining the number of detected photons/molecule/frame were covalently bound to the coverslip coated with 3-APS, and 5xCy3-Tf was adsorbed on the coverslip coated with poly-D-lysine (see the previous subsection). The number of detected photons/molecule/frame was evaluated by multiplying the pixel intensity and the gain conversion factor of the developed camera system. This factor was determined by detecting the same number of photons on both the developed camera system and an Andor iXon DU-897 back-illuminated EM-CCD camera, with the known gain conversion factor of 0.03622 photons/pixel count at an EM gain of 50 and an imaging depth of 16 bits (measured by Zeiss for an ELYRA P.1 PALM system). Briefly, the images of fluorescent beads immobilized on a coverslip were split by a 50/50 beam splitter, and each image was projected onto the new camera system and the EM-CCD camera at the same time. The number of detected photons from each fluorescent bead/frame was obtained by multiplying the gain conversion factor of the EM-CCD camera and the entire pixel intensities of the spot on the EM-CCD camera. The number of photons that reached the EM-CCD camera was then obtained by dividing it by the quantum efficiency of the EM-CCD camera (0.95). The same number of photons must have reached the photoelectric cathode of the developed camera, and therefore, by multiplying with the quantum efficiency of the developed camera (0.4), the number of detected photons/frame/spot was obtained. By comparing this number with the entire pixel count of the fluorescent spot, the conversion factor of the developed camera system was evaluated. At the typical overall electron amplification of 8,100x employed for the observations at a frame rate of 10 kHz (0.1-ms frame time) in the present research, the gain conversion factor of the developed camera system was estimated to be 0.00363 (0.00136) photons/pixel count in an imaging depth of 10 (12) bits for the 1024PCI (SA1) sensor.

### Detecting single-molecule fluorescence spots and determining their positions in the image

Each and every fluorescent spot in an image was detected and its position was determined, by using an in-house computer program based on a spatial cross-correlation matrix (Gelles et al., 1988; Fujiwara et al., 2002, 2016). The image (signal intensity profile) was correlated with a symmetric 2-dimensional Gaussian point spread function (PSF) with a standard deviation of 123 nm (kernel; 123 nm was used for Cy3 and the value was individually determined for each fluorescent probe; see the caption to Fig. S2) (Mashanov and Molloy, 2007). The “maximum entropy” thresholding (Sahoo et al., 1988) was applied to the cross-correlation matrix. The spot representing a single molecule was detected as that containing an area with a size greater than or equal to 5 pixels, which was defined as the size of the area where the values of the cross-correlation matrix were greater than or equal to the threshold value (a constant value for the whole image). The spot position (x, y coordinates) was determined in the following way. First, the centroid of the cross-correlation matrix was calculated, and then an 830-nm-diameter circular region (55.1 nm/pixel x 15 pixels) with its center placed at the centroid was determined as the region that includes the intensity profile of the fluorescent spot. Second, the spot position was determined by fitting the intensity profile in the circular region to a symmetric 2-dimensional Gaussian PSF, using the following formula:

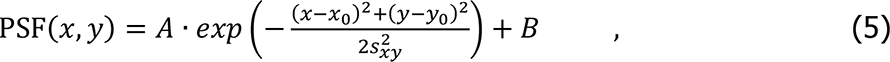

where the fitting parameters were *A* (amplitude), (*x*_o_, *y*_o_) (the sub-pixel center position of the spot), and *s*_xy_ (standard deviation of the symmetric Gaussian PSF), and the value for *B* (background intensity offset) was measured (background signal intensity divided by the number of pixels).

### Observing the time series of the number of detected photons/0.1-ms-frame from single Cy3 molecules, determining the signal-to-noise ratios (SNRs) of single Cy3 molecule images, and evaluating the durations in which single Cy3 molecules could be consecutively observed (the bright/dark periods [on/off-periods] in the images of single Cy3 molecules and the gap closing for short off-periods) (Fig. 2)

Time-dependent changes in the number of detected photons/0.1-ms-frame from single Cy3 molecules immobilized on a coverslip were observed at 10 kHz, under our standard oblique-angle laser illumination conditions of 23 µW/µm^2^ (Fig. 2 A). The molecules that were observable in the first frame of the image sequence were selected. The number of detected photons during a single image frame in an 830 nm diameter (55.1 nm/pixel x 15 pixels) circular region including a fluorescent spot, called the “spot intensity”, was measured and plotted against the frame number at 10 kHz. To evaluate the signal intensity in the background, the number of detected photons during a single image frame in an adjacent region with the same shape and size, called the “BG signal intensity” was measured. On- and off-periods were determined based on the thresholding employed for the spot detection.

The signal-to-noise ratio (SNR) for the image of a single molecule was determined in the following way (Fig. 2 B). From the time-dependent fluorescence signal intensities, the histograms of the numbers of detected photons/0.1-ms for the images of single molecules; i.e., the histograms for the “spot signal intensities” and the “background [BG] signal intensities” during the on-periods, were obtained, and their mean signal intensities (*I*_Spot_ and *I*_BG_, respectively) and standard deviations (*σ*_Spot_ and *σ*_BG_, respectively) were calculated. The SNR was evaluated using the following equation,

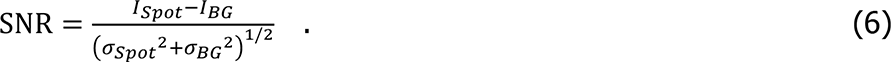

The mean SNR of single Cy3 molecules immobilized on the coverslip was 2.5 ± 0.11 for 50 Cy3 molecules with on-periods of 15 - 150 frames at 10 kHz, whereas the mean SNR on the apical PM was 1.8 ± 0.056 for 50 Cy3-DOPE molecules with on-periods of 1,000 frames at 10 kHz, as obtained under our standard oblique-angle laser illumination conditions of 23 µW/µm^2^.

The distributions of the durations of the on-periods and those after neglecting the off-periods lasting for 1, 2, or 3 frames (called “gap closing”) are shown in Fig. 2 C. The off-periods are induced by the occurrences of non-emission periods of a single dye molecule (which could induce no or dim images in a frame, depending on the duration of the non-emitting period; the off-periods tend to occur more often during the observations of living cells, probably due to the existence of various chemicals in the cell and the culture medium). The off-periods also occur when the signal from a single molecule is missed for short periods, due to the fluctuations of the fluorophore signal intensity (*I*_Spot_) as well as the higher background (*I*_BG_) and its fluctuation (*σ*_BG_). Therefore, gap closing for 1-3 frames is often employed in single-molecule tracking studies. For describing the results obtained in living cells in the present report, we used 3-frame gap closing.

### Cell culture

Human T24 epithelial cells (the same as the ECV304 cell line used in the previous research [Murase et al., 2004], which turned out to be a sub-clone of T24; Dirks et al., 1999) were grown in Ham’s F12 medium (Sigma-Aldrich) supplemented with 10% fetal bovine serum (Sigma-Aldrich). Cells were cultured on 12-mm diameter glass-bottom dishes (IWAKI), and single-molecule observations were performed two days after inoculation. For culturing cells expressing mGFP-paxillin or mEos3.2-paxillin, the glass surface was coated with fibronectin by an incubation with 5 µg/ml fibronectin (Sigma-Aldrich) in phosphate-buffered saline (PBS) (pH 7.4) at 37°C for 3 h.

To form PM blebs (up to 20 µm in diameter) depleted of the actin-based membrane-skeleton, the cells were incubated with 1 mM menadione for 5 h at 37°C (Fujiwara et al., 2002), and then treated with 100 nM latrunculin A (a gift from G. Marriott, University of California, Berkeley) on the microscope stage for 5 min at 37°C. All measurements were performed within 5-15 min after the addition of latrunculin A (Suzuki et al., 2005; Fujiwara et al., 2016).

### Examination of the photo-induced damage to the cell during ultrahigh-speed single fluorescent-molecule imaging (Fig. 2, D and E)

The viability of the cells was examined by staining with 1 µM TOTO-3 iodide (Molecular Probes), which only stains dead cells (Zuliani et al., 2003), at 37°C for 5 min and then observing the stained cells using epi-illumination with a 594-nm laser. To obtain reference images of dead cells, the cells were treated with 100 µM H_2_O_2_ at 37°C for 1 h. The fluorescence intensity of TOTO-3 in the 5.5 x 5.5-µm area inside the nucleus was measured, and the histograms of signal intensities were obtained. Based on the negative and positive controls (Fig. 2, D **and** E), a threshold fluorescence intensity of 4.0 x 10^5^ (arbitrary unit = AU) was selected to categorize the live and dead cells (96% [90%] of the negative [positive] control cells were categorized as alive [dead]).

Note the following. Once the desired single-molecule localization precision is chosen for each experiment, the required total number of photons emitted from the molecule during a frame to achieve the precision will be the same, irrespective of the frame rate (inverse of the frame time). Therefore, the total number of photons elicited by laser excitation during a frame time will be the same, regardless of whether the observations are made at video rate (33-ms frame time) or at 10 kHz (0.1-ms frame time, if photon emission saturation does not occur). Namely, the direct photo-damage to the cell per frame due to the excitation laser illumination is expected to remain constant even when the laser illumination intensity and the frame rate are increased, as long as the single-molecule localization precision required for the experiment and *the number of observed frames* (not the total observation duration) are the same (in the absence of photon emission saturation).

However, direct experiments are required to test the phototoxicity to cells, as shown here. This is partly because slight photon-emission saturation occurred under our standard illumination conditions of 23 µW/µm^2^ (Fig. 1 E), and also because higher illumination photon densities could induce two-photon events and enhance the secondary reactions of the photo-induced molecules produced at high densities in the cell, which could generate cytotoxic substances.

### Preparation of Cy3-DOPE and Cy3-Tf, and cell surface labeling

Cy3-DOPE was prepared and incorporated in the PM as described previously (Fujiwara et al., 2002; Murase et al., 2004), except that the final concentration of Cy3-DOPE added to the cells was 10 nM. Cy3-Tf used for the measurements at 6 kHz (0.167-ms resolution) was generated by incubating 6.5 µM monofunctional sulfo-Cy3 (GE Healthcare) with 6.3 µM human holo Tf (Sigma-Aldrich) in 0.1 M carbonate buffer (1 ml, pH 9.0) at 25°C for 60 min, followed by desalting column chromatography (PD-10, GE Healthcare, equilibrated and eluted with PBS) to remove the unreacted dye. The dye/protein molar ratio of the Cy3-Tf was 0.2. At this ratio, more than 90% of the fluorescent spots in the image are expected to represent single Cy3 molecules, based on the Poisson distribution, which was confirmed by single-step photobleaching of the fluorescent spots of Cy3-Tf (as well as Cy3-DOPE). Tf conjugated with multiple Cy3 molecules, used for the measurement at 45 kHz (0.022-ms resolution), was generated by incubating 1.25 mM monofunctional sulfo-Cy3 with 6.3 µM human holo Tf (mixed at a molar ratio of ∼200:1), which provided Cy3-Tf with the dye/protein molar ratio of ∼5 (called 5xCy3-Tf).

To label TfR in the PM with Cy3-Tf, first the Tf (originated from FBS) already bound to TfR on the cell surface was partially removed by incubating the cells in 1 ml Hanks’ balanced salt solution buffered with 2 mM TES (Dojindo), at pH 7.4 (HT medium) and 37°C for 10 min, and then after washing, the HT medium containing 10 nM Cy3-Tf (or 5xCy3-Tf) was added to the cells at a final concentration of 0.5 nM. At this concentration, single molecules of Cy3-Tf bound to the apical PM could be observed without replacing the medium with the Cy3-Tf-free medium. The observations were completed within 10 min after the Cy3-Tf addition.

### Ultrahigh-speed imaging of single fluorescent molecules and its application to fluorescent probes bound to coverslips

Each individual fluorescently-labeled molecule on the apical and basal PMs was observed at 37 ± 1°C, using the oblique-angle and TIR illumination modes, respectively, of a home-built objective lens-type TIRF microscope (based on an Olympus IX70 inverted microscope), which was modified and optimized for the camera system developed here. The beam was attenuated with neutral density filters, circularly polarized, and then steered into the edge of a high numerical aperture (NA) oil immersion objective lens (UAPON 150XOTIRF, NA = 1.45, Olympus), focused on the back focal plane of the objective lens. Both TIR and oblique-angle illuminations were used for Cy3 at 10 kHz and 30 kHz, and 5xCy3-Tf at 45 kHz. For other molecules and conditions, only the TIR illumination was used. For exciting Cy3, Alexa555 (Invitrogen), JF549 (Janelia Fluor 549; Tocris Bioscience), and TMR (Molecular Probes and Promega), a 532-nm laser (Millennia Pro D2S-W, 2W, Spectra-Physics; 1.8 to 16 mW for the ∼14-µm diameter observation area) was employed. For exciting JF646 (Janelia Fluor 646; Tocris Bioscience), Atto647N (ATTO-TEC), SeTau647, and Cy5 (sulfo-Cy5; GE Healthcare), a 660-nm laser (Ventus, 750 mW, Laser Quantum; 0.41 to 4.5 mW for the ∼12-µm diameter observation area) was used.

The illumination laser power density was determined as follows. When the field stop was fully open, the excitation laser illuminated a circular (2-dimensional Gaussian) area with a 12.5-µm radius (standard deviation) on the sample plane. By adjusting the field-stop size, the central part of the excitation laser beam was selected to form a circular area with a radius of 7 µm on the sample plane, approximating the largest view-field of 14 x 14 µm^2^ employed in this research, and the laser power after the objective lens was measured. Since the dimmest intensity in the 7-µm Gaussian area was ∼85% of the peak intensity, the laser power density was calculated assuming a uniform laser intensity in the circular area with a 7-µm radius; i.e., the measured power was simply divided by the area size of 154 µm^2^ (π x 7^2^).

To observe single fluorescent molecules immobilized on coverslips, dye molecules were covalently linked to 3-aminopropyltriethoxysilane (APS)-coated coverslips (custom-ordered from Matsunami). Briefly, 0.2 – 1 nM succinimidyl ester-modified dye molecules in HT medium were placed on the 3-APS-coated coverslip at 25°C (10-min incubation), and then the coverslip was washed five times with 1 ml HT medium. To immobilize 5xCy3-Tf molecules on the glass surface, the 12-mm diameter glass-bottom dishes were coated with poly-D-lysine, and then incubated in HT medium containing 0.1 nM 5xCy3-Tf at 25°C for 10 min.

The method for evaluating the position determination precisions of Cy3-labeled molecules bound on the coverslips is described in the caption to Fig. S2. The position determination precisions for the molecules in the apical PM were estimated using the ensemble-averaged MSD-Δ*t* plots for single-molecule trajectories of Cy3-DOPE and Cy3-Tf (5xCy3-Tf) bound to TfR, as shown in Fig. S4.

### Quantitative analysis of single-molecule trajectories based on the MSD-Δ*t* plots (Fig. 4, A-a and B)

For each single-molecule trajectory, the one-dimensional MSD(*x* or *y*) for the x- or y-direction for every time interval was calculated according to the following formula:

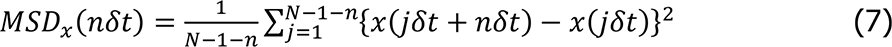

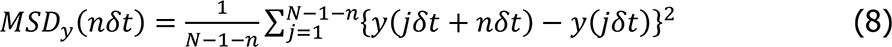

where *δt* is the frame time, (*x*(*jδt* + *nδt*) and *y*(*jδt* + *nδt*)) describe the position of the molecule following a time interval *nδt* after starting at position (*x*(*jδt*), *y*(*jδt*)), *N* is the total number of frames in the sequence, *n* and *j* are positive integers, and *n* determines the time increment. In the quantitative analysis of the trajectories, 1-dimensional MSD-Δ*t* plots were employed.

Each single-molecule trajectory was classified into a suppressed-, simple-Brownian-, or directed-diffusion mode using the 2-dimensional MSD-Δ*t* plot, which is the sum of the MSD-Δ*t* plots for the x- and y-directions (Fig. 4 A-a). The basic idea for the classification is shown in Fig. 4 B-a (Kusumi et al. 1993). The classification was based on the relative deviation (RD), which describes the long-term deviation of the actual mean-square displacement, *MSD*(*N*, *n*) at the time *nδt*, from the expected MSD based on the initial slope of the MSD-Δ*t* plot for molecules undergoing ideal simple-Brownian diffusion, 4*D*_2-4_ · *nδt*; i.e., *RD*(*N*, *n*) = *MSD*(*N*, *n*)/[4*D*_2-4_ · *nδt*] (*N* = the number of frames in a full trajectory, *n* = the increment number of frames used for the analysis with the MSD-Δ*t* plot, 1 ≤ *n* ≤ *N*, and *δt* = duration of each frame. Therefore, Δ*t* = *nδt*, which is the x-axis of the *MSD*(*N*, *n*)-Δ*t* plot; and *D*_2-4_ is the short-time diffusion coefficient determined from the slope of the second, third, and fourth points in the MSD-Δ*t* plot). The RD value is ≪, ≈, or ≫ 1, when the molecules are undergoing suppressed, simple-Brownian, or directed diffusion, respectively.

In Fig. 4 B-b, the distributions of *RD*(*N*, *n*)s for the Cy3-DOPE (*RD*(1,000, 100), *δt* = 0.1 ms) and TfR trajectories (*RD*(1,000, 200), *δt* = 0.167 ms) (shaded bars; *n* = 50 and 20 for the intact and actin-depleted blebbed PM), as well as for simple-Brownian particles generated by a Monte Carlo simulation (open bars; *n* = 5,000), are shown. Based on the distribution of the *RD*(*N*, *n*) values determined for simulated simple-Brownian particles (open bars), the *RD*(*N*, *n*) values giving the 2.5 percentiles of the particles from both ends of the distribution, referred to as *RD*_min_ and *RD*_MAX_, were obtained (red and blue vertical lines in Fig. 4 B-b, respectively; they depend on both *N* and *n*). Each experimental single-molecule trajectory was classified into the suppressed (confined and hop)-diffusion mode if its *RD*(*N*, *n*) value (shaded bar) was smaller than *RD*_min_ (and into the directed-diffusion mode if its *RD*(*N*, *n*) value was larger than *RD*_MAX_).

### Monte Carlo simulations for the hop diffusion in the intact PM and simple-Brownian diffusion in the actin-depleted blebbed PM (Fig. 5 B)

Monte Carlo simulations for the hop diffusion were performed as described previously (Ritchie et al., 2005). The hop diffusion simulation parameters are the following (*p*represents the probability that a hop movement to an adjacent compartment takes place when the diffusing molecule enters the boundary).

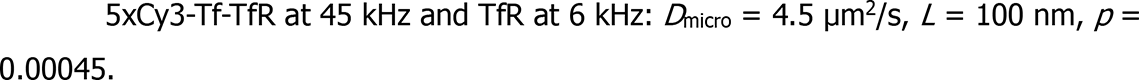

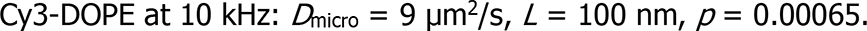

Simulation parameters for simple-Brownian diffusion in the actin-depleted blebbed PM are the following.

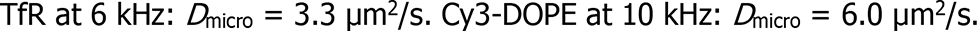

Gaussian localization error of 50 nm was added (see Fig. S4 B). The MSD-Δ*t* plot ensemble-averaged over 1,000 trajectories was obtained, and the offset due to the localization error was estimated and subtracted based on the y-intercept (by extrapolation) of the linear-fit function for the second, third, and fourth steps in the MSD-Δ*t* plot, as performed for experimental trajectories.

### Ultrafast live-cell PALM of caveolin-1-mEos3.2 and mEos3.2-paxillin (Figs. 6-8)

The plasmid encoding mouse caveolin-1 (GenBank: U07645.1) fused to mEos3.2 (caveolin-1-mEos3.2) was generated by replacing the cDNA encoding the EGFP protein, in the caveolin-1-EGFP plasmid (a gift from T. Fujimoto, Nagoya University, Japan; Kogo and Fujimoto, 2000), with that of mEos3.2 (Zhang et al., 2012) (generated by making 3 point mutations, I102N, H158E, and Y189A in the Addgene mEos2 plasmid #20341 [http://n2t.net/addgene:20341; RRID:Addgene_20341], a gift from L. Looger, Janelia Research Campus; McKinney et al., 2009). The plasmid encoding mEos3.2-paxillin was generated from the mGFP-paxillin plasmid, used in the companion paper (Fujiwara et al., 2021). First, the cDNA encoding human paxillin isoform alpha (NCBI reference sequence: NM_002859.3; cloned from the human WI-38 cell line) fused to mGFP at the N-terminus was subcloned into the EBV-based episomal vector, pOsTet15T3 (gift from Y. Miwa, University of Tsukuba, Japan), which bears the tetracycline-regulated expression units, the transactivator (rtTA2-M2), and the TetO sequence (a Tet-on vector) (Shibata et al., 2012, 2013). The mEos3.2-paxillin plasmid was then generated by replacing the cDNA encoding mGFP in the mGFP-paxillin plasmid with that encoding mEos3.2. T24 cells were transfected with the cDNAs encoding caveolin-1-mEos3.2 and mEos3.2-paxillin, using Nucleofector 2b (Lonza) according to the manufacturer’s recommendations.

Data acquisitions of ultrafast live-cell PALM were performed on the basal PM at 37 ± 1°C. We used the TIR illumination mode of our home-built TIRF microscope (based on a Nikon Ti-E inverted microscope) equipped with two ultrafast camera systems developed here and a high NA oil immersion objective lens (CFI Apo TIRF 100x, NA = 1.49, Nikon). Photoconversion of mEos3.2 was induced by a 405-nm diode laser (PhoxX, 120 mW, Omicron) with its laser intensity exponentially increased from 0.014 to 0.036 µW/µm^2^ during the PALM data recording period of 10 s with an e-folding time *τ* of 10.6 s (i.e., the photoconversion laser intensity at time *t* = 0.014 µW/µm^2^ x exp[t/10.6], and thus that at the end of the 10-s data accumulation time = 0.036 µW/µm^2^ as *t* = 10 s), to keep the number densities of mEos3.2 spots in the FA region similar during the e-folding period.

The photoconverted mEos3.2-paxillin was excited using a 561-nm laser (Jive, 500 mW, Cobolt) at 30 µW/µm^2^, and single-molecule spots were recorded at a frame rate of 1 kHz (with an integration time of 1 ms). The fluorescence image isolated by a bandpass filter of 572-642 nm (FF01-607/70, Semrock) was projected onto the photocathode of the developed ultrafast camera system (more specifically, the photocathode of the image intensifier). The reconstructions of a PALM image with a pixel size of 10 nm were performed using the ThunderSTORM plugin for ImageJ (Ovesny et al., 2014) installed in the Fiji package (Schindelin et al., 2012) and Gaussian rendering with a localization precision of 29 nm (Fig. 6 A-d).

### Detection and characterization of FA-protein islands using Voronoï-based segmentation of the PALM images (Fig. 8)

The SR-Tesseler software based on Voronoï polygons (Levet et al., 2015) was applied to the locations of mEos3.2-paxillin to automatically segment the PALM data. Since the blinking correction function of the SR-Tesseler software evaluated that the average number of blinks was 1.1 (the 1-frame gap closing was applied to sequences of captured images using the “Merging” post-processing function in ThunderSTORM, with the maximum off frame of 1 and the maximum search distance of twice the mean localization precision of mEos3.2 in the basal PM of 29 nm; Fig. 6 A-d), further blinking correction was not performed with the SR-Tesseler software. FA contour detection was performed by using a Voronoï polygon density factor of 1.2 (within the ROI), with a minimum area of 2 pixels. The contour of the islands of FA proteins was determined basically in the same way, by using a Voronoï polygon density factor of 1.2 (within the FA contour), with a minimum area of 0.05 pixels.

## Data availability

Data supporting the findings of this study are available from the corresponding author upon reasonable request.

## Code availability

The code is available from the corresponding author upon reasonable request.

## Online supplemental material

Fig. S1 shows how we determined the electron amplification gain of the image intensifier. Fig. S2 shows the relationship of the single-molecule localization precision with the number of detected photons/molecule/frame for various fluorescent dye molecules. Fig. S3 shows that the numbers of detected photons/molecule/frame from single molecules are saturated under stronger excitation laser intensities, and thus the improvement of single-molecule localization precision with an increase of the laser intensity is limited. Fig. S4 shows the MSD-Δ*t* plots ensemble averaged over all trajectories obtained for Cy3 molecules immobilized on coverslips or diffusing in the apical and basal PM of T24 cells, providing estimates of single-molecule localization errors. Fig. S5 shows the TILD method, the expected distribution of the dwell lifetimes (theoretical derivation), and the derivation of the equation used for hop diffusion fitting. **Video 1** shows single Cy3 molecules covalently linked to the glass surface observed at 10 kHz. **Video 2** shows single Cy3-DOPE molecules diffusing in the intact apical PM in a view-field of 256 x 256 pixels, observed at 10 kHz. **Video 3** shows enlarged views of single Cy3-DOPE molecules diffusing in the intact apical PM and the actin-depleted blebbed PM, observed at 10 kHz. **Video 4** shows enlarged views of single TfR molecules, bound by Cy3-Tf, diffusing in the intact apical PM and the actin-depleted blebbed PM, observed at 6 kHz.

## Acknowledgements

We thank Profs. G. Marriott of the University of California, Berkeley, Y. Miwa of the University of Tsukuba, T. Fujimoto of Nagoya University School of Medicine, and L. Looger of the Janelia Research Campus, for their kind gifts of latrunculin A, the pOsTet15T3 vector, the cDNA encoding caveolin-1-EGFP, and the mEos2 plasmid, respectively. We also thank Mr. K. Hanaka for his enthusiastic encouragement of this research, Ms. M. Yahara and Ms. A. Chadda for constructing various cDNAs, Ms. J. Kondo-Fujiwara and Mr. K. Kanemasa for preparing the figures, and the members of the Kusumi laboratory for helpful discussions. We are grateful to Prof. Ken Jacobson of the University of North Carolina and Prof. Kathalina Gaus of the University of New South Wales Sydney for critical reading of the manuscript and constructive comments. This work was supported in part by Grants-in-Aid for Scientific Research from the Japan Society for the Promotion of Science (JSPS) (Kiban B to T.K.F. [16H04775, 20H02585], Kiban B to K.G.N.S. [18H02401], and Kiban S and A to A.K. [16H06386 and 21H04772, respectively]), a Grant-in-Aid for Challenging Research (Exploratory) from JSPS to T.K.F. (18K19001), a Grant-in-Aid for Innovative Areas from the Ministry of Education, Culture, Sports, Science and Technology of Japan (MEXT) to K.G.N.S. (18H04671), Japan Science and Technology Agency (JST) grants in the Core Research for Evolutional Science and Technology (CREST) program in the field of “Biodynamics” to A.K. and in the field of “Extracellular Fine Particles” to K.G.N.S. (JPMJCR18H2), and a JST grant in the program of the Development of Advanced Measurements and Analysis Systems to A.K. and T.K.F. WPI-iCeMS of Kyoto University is supported by the World Premiere Research Center Initiative (WPI) of MEXT.

## Author contributions

T.K.F. and A.K. conceived and formulated the project. T.K.F., S.T., Y.N., K.I., and A.K. developed the ultrahigh-speed camera system, T.K.F., K.I., T.A.T., and A.K. developed an ultrafast single-molecule tracking station based on the newly developed camera system, and T.K.F., T.K., T.A.T., and A.K. tested the camera system on the developed station. T.K.F., T.A.T., K.G.N.S., and A.K. designed the biological experiments and participated in discussions. T.K.F. performed virtually all of the ultrahigh-speed single-molecule imaging-tracking experiments, the ultrafast PALM experiments, and the data analysis. T.K.F. and K.P.R. developed the compartment detection algorithm. Z.K., based on discussions with T.K.F., derived the equation describing the MSD-Δ*t* plot for particles undergoing hop diffusion and developed the theory for the distribution of the dwell lifetimes within a compartment for particles undergoing hop diffusion. T.K.F., K.P.R., Z.K., and A.K. evaluated the data. T.K.F., Z.K., and A.K. wrote the manuscript, and all authors discussed the results and participated in revising the manuscript.

## Supplementary Figures

**Figure S1.**
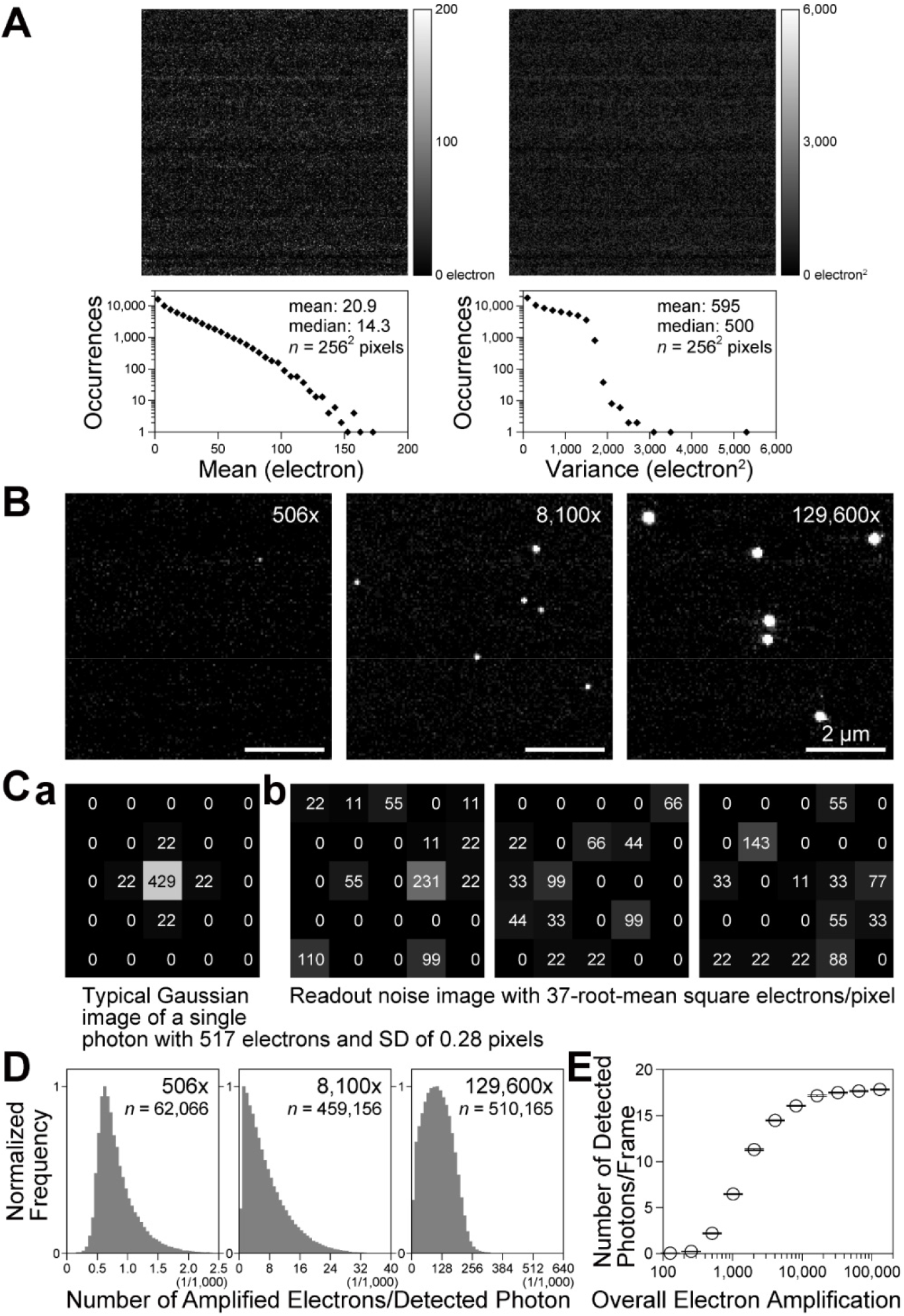
Establishing the intensifier set up for detecting single photons using the electron amplification of the image intensifier. (**A**) The readout noise of individual pixels of the CMOS chip used here. The map (**top**) of the means (**left**) and variances (**right**) of pixels (256 x 256 pixels) and their histograms (**bottom**) (Huang et al., 2013) for 1,000 frames obtained at 10 kHz. (**B**) Typical images of single photons obtained as a function of the intensifier gain, using the newly developed camera system. They were obtained by amplifying single electrons emitted from the image-intensifier photocathode by the arrivals of single photons (uniform Köhler illumination by the strongly attenuated halogen lamp of the microscope). The amplifications were 506, 8,100, and 129,600 (increases by a factor of 16 from left to middle and middle to right images). Note that the term “overall electron amplification (of the camera system)” always excludes the 40% quantum efficiency of the image intensifier photocathode throughout this report, because we discuss the amplification of the number of electrons from the photoelectrons emitted by the photocathode. The range of the gray levels of the images shown here (both **B and C**) was set for 8 bits from 0 to 550 electrons/pixel at the CMOS sensor (from black to white). (**C**) As an example, we show that an overall electron amplification of 506x by the camera system would not be sufficient for the consistent detection of single photons, due to the readout noise. (**a**) A schematic figure showing a 7 x 7 pixelated image for a single detected photon amplified to a total of 517 electrons (without noise and background; computer-generated assuming an overall amplification of 506x with an SD of 0.28 ± 0.085 pixels as in **B left**). The number of amplified electrons in each pixel on the CMOS sensor is shown. A full well of 45,000 electrons/pixel of the CMOS sensor (SA1) is scaled to 12 bits (4,095 camera counts, and hence a unit camera count = 10.99 [∼11] electrons per pixel), and thus each number is a multiple of 11 electrons. (**b**) Three arbitrarily selected (experimentally obtained) 7 x 7 pixelated images representing the spatial distributions of the readout noise of the CMOS sensor employed in this study (37-root-mean-square electrons/pixel/frame; see “Ultrahigh-speed intensified CMOS camera system: design and operation” in **Materials and methods**). To detect a (photon-converted) emitted electron, its image, such as that shown in **a**, must be detectable in the presence of spatiotemporally varying noise, as shown here. The detectability will be enhanced by an increase of the electron amplification. See **D**. (**D**) Stochastic gain variations (fluctuations) of the image intensifier. At the level of detecting single photons, the gain variations are large (at the level of detecting single molecules, relative variations will become smaller due to averaging over all detected photons). In these histograms, the distributions of the number of total electrons stored at the CMOS sensor of the camera system for each single detected photon at the image-intensifier photocathode (i.e., for each discernible spot in images like those in **B**) are shown for the overall electron amplifications of 506x, 8,100x, and 129,600x (the y axis is normalized by the peak value in each histogram; the full x scale is increased by a factor of 16 from the left figure to the middle figure and from the middle figure to the right figure). The spots in the images (like those in **B**) were identified and the total number of electrons in each spot was evaluated by using the functions of the ThunderSTORM plugin of ImageJ (Ovesny et al., 2014). For the spot detection with localization precisions at the pixel level (the peak pixel), we employed the “Wavelet filtering” (B-Spline order = 3 and B-Spline scale = 2.0) and the “Local maximum method” (Peak intensity threshold = 4.5 and Connectivity = 8-neighborhood). For the determination of the total number of electrons in a spot (and subpixel localization of each spot), we performed the Gaussian fitting of the image (i.e., the number of electrons/pixel in 11×11 pixels surrounding the peak pixel) and then integrated the best-fit function, using the Subpixel localization of molecules. These histograms were obtained by using six 5,000-frame image sequences recorded at 10 kHz with a frame size of 13.4 x 13.4 µm on the focal plane, detecting 62,066, 459,156, and 510,165 photons (electrons emitted from the photocathode of the image intensifier) for overall amplifications of 506x, 8,100x, and 129,600x, respectively. They represent both the stochastic gain variations and the detectability of a photon image produced by the amplified electrons. See the histogram for 506x. The occurrences of the number of amplified electrons/detected photon sharply decreased when the number of amplified electrons was reduced below ∼650 electrons (the peak in the histogram). This is very likely due to a sharp reduction in the detectability of the spots produced by <650 electrons. Namely, when the signal intensity (the number of electrons) after amplification is small, the chance that the signal becomes less than the readout noise increases, because the readout noise pattern also fluctuates spatiotemporally (see **C**). (**E**) The probability of detecting a single photon was increased to 90.0% of the saturation level, at an overall electron amplification of 8,100x. Here, the number of detected photons per frame (mean ± SD); i.e., the number of discernible spots in the images like those in **B** (detected by the method described in **D**), is plotted as a function of the overall electron amplification (averaged over six 5,000-frame image sequences). With an increase in the overall electron amplification, the number of discernible spots increases. At an overall electron amplification of 8,100x, the number of discernible spots was 90.0% of the saturated number of spots (i.e., at the maximal overall electron amplification possible with the present instrument, which is 129,600x amplification). At the overall amplifications giving the saturated number of spots, virtually every photoelectron emitted from the photocathode is considered to be detected. Therefore, throughout the present research, we employed 8,100x as the overall electron amplification of the image intensifier (see “Ultrahigh-speed intensified CMOS camera system: design and operation” in **Materials and methods**).

**Figure S2.**
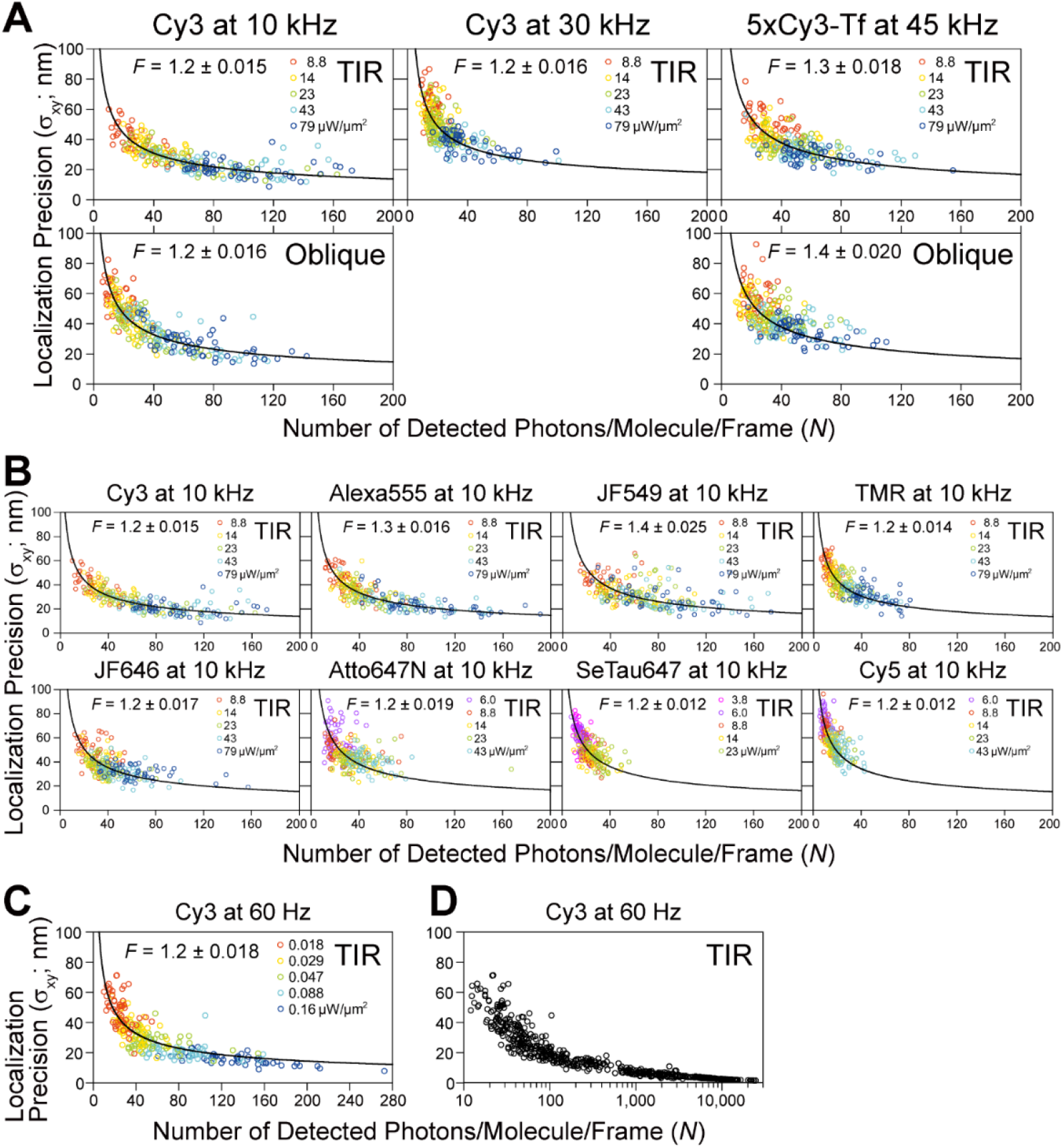
Plots providing the number of detected photons (proportional to the emitted photons) from a single fluorescent molecule during a single frame time (*N*; x-axis), used for obtaining the desired single-molecule localization precisions (*σ*_xy_ = [*σ*_x_ + *σ*_y_]/2; y-axis) and clarifying which fluorescent dye molecules can give the required precisions. The results for 30, 10, and 0.06 kHz; i.e., the frame times of 0.033, 0.1 and 16.7 ms, respectively, are shown (45 kHz/0.022 ms for 5xCy3-Tf is also shown). For the method to evaluate *N*, see **Materials and methods**, “Determination of the number of detected photons/molecule/frame (*N*)”. In high-speed single fluorescent-molecule imaging, one of the key problems is whether single fluorescent molecules emit sufficient numbers of photons (during a single frame time) required for obtaining the desired single-molecule localization precisions. The results shown here indicate that a 10-kHz frame rate is applicable for various dye molecules, and Cy3 could even be used at 30 kHz. These plots were fitted well by the theoretical equation derived previously (Mortensen et al., 2010), indicating that the developed camera system functions as planned, even at high frequencies. The “excess noise” factor (*F*) of the developed camera system was evaluated by this fitting. Throughout this report (except for the measurements in the PM, as described in Fig. S4), the localization precision (*σ*_xy_) is defined as [*σ*_x_ + *σ*_y_]/2, where *σ*_x_ and *σ*_y_ are the standard deviations of the *x* and *y* position determinations, respectively, following the convention of the super-resolution imaging field (Dietrich et al., 2002; Martin et al., 2002). *σ*_xy_ was determined in 15 consecutive frames for *n* = 50 trajectories for each condition. All of the fluorescent dye molecules were covalently bound to coverslips coated with 3-aminopropylethoxysilane, and 5xCy3-Tf was adsorbed on the coverslip coated with poly-D-lysine (**Materials and methods**). (**A** and **B**) Plots for single Cy3 molecules observed at 10 and 30 kHz and single 5xCy3-Tf molecules observed at 45 kHz (**A**) and those for various fluorescent molecules observed at 10 kHz (**B**). Five TIR laser illumination intensities were employed for each dye (50 molecules for each laser intensity), as indicated by the different colors of the data points. Various ranges of the laser power densities were used for different dyes, because the dyes are saturated differently (shown in each box). The plots (*σ*_xy_ vs. *N*) shown in **A** and **B** could be fitted well (non-linear least-squares fitting by the Levenberg–Marquardt algorithm) using the following equation derived previously (Mortensen et al., 2010).

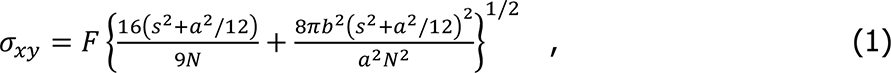

where *F* is the sole fitting parameter, representing the “excess noise” factor (a coefficient describing the stochastic gain fluctuation in the electron amplification process in the image intensifier; *F* is shown in each box, but its value is 1.2 – 1.4 for all cases), *s* is the standard deviation of the Gaussian spot profile, 123 ± 1.1 nm for Cy3 on the sample plane (determined by the Gaussian fitting of each image for 50 Cy3 molecules immobilized on the glass excited by the TIR illumination at 79 µW/µm^2^; compared with our standard condition of the oblique illumination at 23 µW/µm^2^, these observation conditions provided approximately 3 times more detected photons; see Fig. 1 E**, top**; note that *s* depends on the observed fluorescent molecules), *a* is the pixel size (55.1 nm), and *b* is the standard deviation of the background noise. (For example, 0.038 ± 0.059 detected photons/pixel/frame [mean ± SD] for the TIR illumination and 0.035 ± 0.058 detected photons/pixel/frame for the oblique illumination at 10 kHz; *n* = 76,800 pixels = 32 x 32 pixels x 15 frames x 5 different positions.) The estimated excess noise factor *F* of the image intensifier shows that it is comparable to or slightly smaller (less noisy) than that of the EM-CCD electron multiplier (*F* = 1.4). Cy3 exhibited the least tendency to saturate, and thus provided better single-molecule localization precisions, consistent with the analysis results shown in Fig. S3, A **and** C. Note that *s* and *b* were determined for each fluorescent probe (with different illumination and excitation wavelengths and optics). 5xCy3-Tf data are considered to represent fluorescent spots generated by various numbers of Cy3 molecules placed within a few nanometers, mostly in the range of 3 to 8 molecules (1 and 2 Cy3 molecules/Tf, representing ∼12% of the 5xCy3-Tf spots, gave low signals, inducing extremely large errors in single-molecule localizations; meanwhile, the probability of 9 or more Cy3 molecules being attached to a Tf molecule will be less than 7%). Due to the photobleaching of multiple Cy3 molecules bound to a Tf molecule, the numbers detected on a Tf molecule decreased quickly upon laser illumination. (**C**) Plot for single Cy3 molecules observed at 60 Hz, with TIR illumination laser power densities ≤ 0.16 µW/µm^2^ (indicated by different colors of the data points; 50 molecules for each laser intensity). The excess noise factor *F* was estimated to be 1.2, consistent with the results shown in **A**. (**D**) Summary plot for single Cy3 molecules observed at 60 Hz, with TIR illumination laser power densities up to 79 µW/µm^2^: 0.018, 0.029, 0.047, 0.088, 0.16 (employed for the plot in **c**), 0.48, 1.6, 4.8, 14, 23, 43, and 79 µW/µm^2^ (note that in this plot, in contrast to the others, the x-axis is in the log scale). The single-molecule localization precisions obtained with the laser power densities equal to and greater than 14 µW/µm^2^ were calculated by using Eq. 1 with *F* = 1.2, as found in **C**. This method for obtaining the single-molecule localization precisions employed here is different from that used for evaluating the precisions shown in Fig. 1 F, Fig. S2, A-C, and Fig. S3, B and C. Since most Cy3 molecules were photobleached within a single 16.7 ms frame, the more-prevalent method could not be employed. The x-axis of this figure covers the entire practical scale for the number of detected photons/molecule/frame (*N*) for a single Cy3 molecule, from 25.0 ± 1.4 at a laser power density of 0.018 µW/µm^2^ up to 11,400 ± 700 at 79 µW/µm^2^ (mean ± SEM). This upper limit was given by the photobleaching and excitation power saturation of Cy3, and provided the best single-molecule localization precision of 2.6 ± 0.099 nm (mean ± SEM) for Cy3 (no further improvements could be obtain even by employing higher laser intensities; Fig. 1, G **and** H).

**Figure S3.**
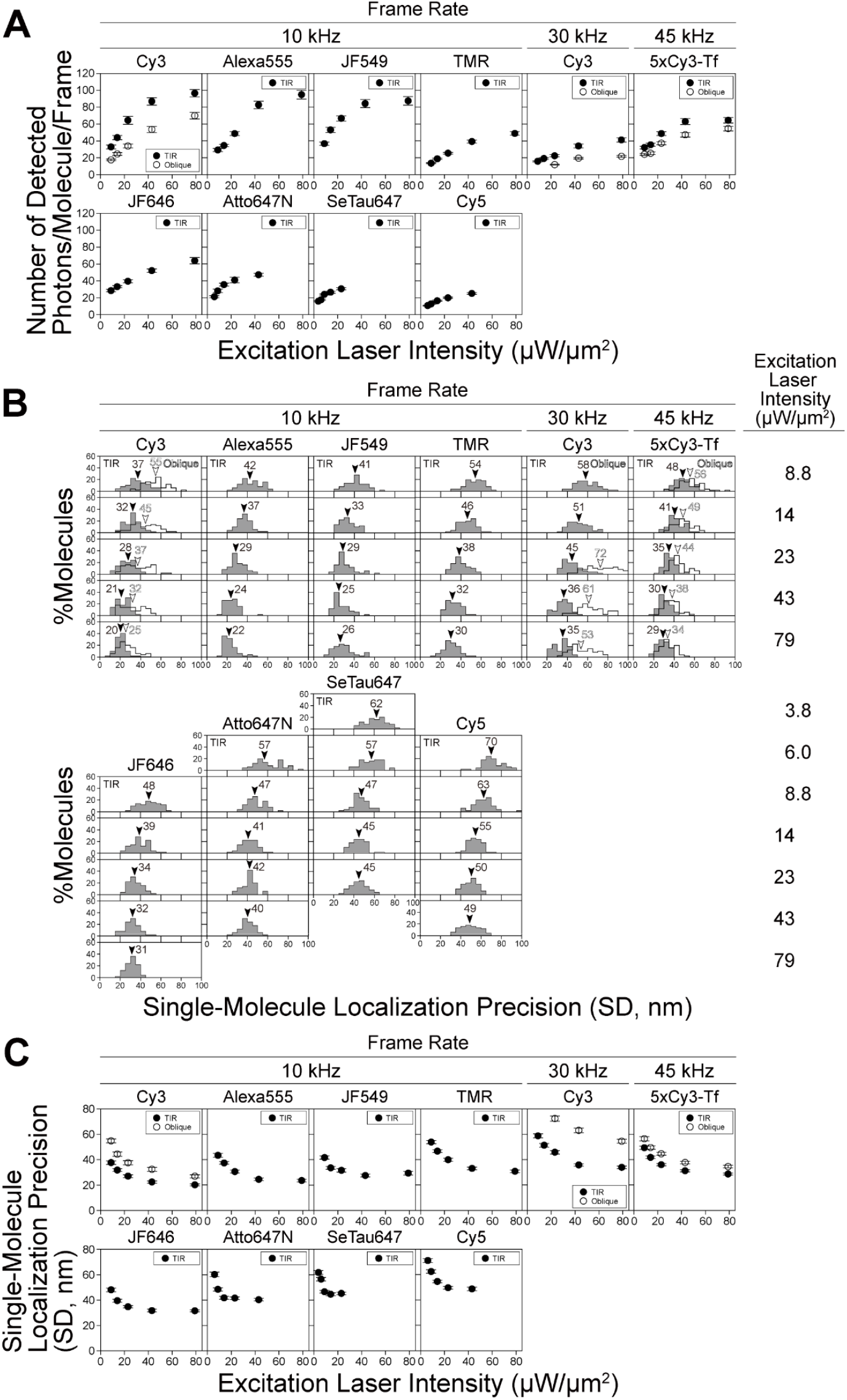
The numbers of detected photons/frame from single molecules (*N*, proportional to the emitted photons during a frame time) are saturated under stronger excitation laser intensities, and thus the improvement of single-molecule localization precision (*σ*_***xy***_) with an increase of the laser intensity is limited. For experimental details, see the caption to Fig. S2. (**A**) The numbers of detected photons/molecule/frame plotted against the laser power density at the focal plane (TIR illumination; the additional examinations using oblique illuminations were only performed for Cy3 and 5xCy3-Tf). With an increase in the excitation laser intensity, the number of detected photons from single dye molecules initially increased proportionally, and then leveled off (saturation occurred), probably due to “triplet bottleneck saturation” (see “Estimation of the number of photons that can be emitted by a single Cy3 molecule during 0.1 ms: triplet bottleneck saturation” in **Materials and methods**). This occurred from around 23 µW/µm^2^ for Cy3 (the results of Cy3 at 10 and 30 kHz shown here are the same as those shown in Fig. 1 E, and are reproduced here for ease of comparison with the results of other dyes). Cy3 and Alexa555 are less prone to saturation, as compared with the other dyes tested here. In the present study, we primarily used Cy3. (**B**) Distributions of the localization precisions of single molecules of eight fluorophores observed at 10 kHz (an integration time of 0.1 ms), single Cy3 molecules at 30 kHz (0.033 ms), and 5xCy3-Tf at 45 kHz (0.022 ms), evaluated at various laser illumination intensities (provided on the right). Arrowheads indicate the median values. (**C**) Single-molecule localization precisions (mean ± SEM) plotted against the laser intensity (the results for Cy3 at 10 and 30 kHz shown here are the same as those shown in Fig. 1 F, and are reproduced here for ease of comparison with the results of other dyes). These results show that Cy3 and Alexa555 provide better single-molecule localization precisions at higher laser intensities, due to their lower tendency to saturate. Since Cy3 provided slightly better single-molecule localization precision at 10 kHz at saturation than Alexa555, we primarily used Cy3 throughout the remaining part of this report. Based on the results described in **A** and **C**, and also due to the versatility of the oblique-angle illumination to enable the observations of single molecules in both the basal and apical PMs, as well as in endomembranes and the cytoplasm in general (see the subsection “TIR and oblique illuminations for ultrafast single-molecule imaging” in the **Main Text**), we comprehensively tested and performed ultrafast single-molecule imaging-tracking under the oblique-angle illumination conditions at a laser power density of 23 µW/µm^2^ at the specimen plane. These are the “**standard test conditions**”, using Cy3 molecules on the coverslip as the standard sample.

**Figure S4.**
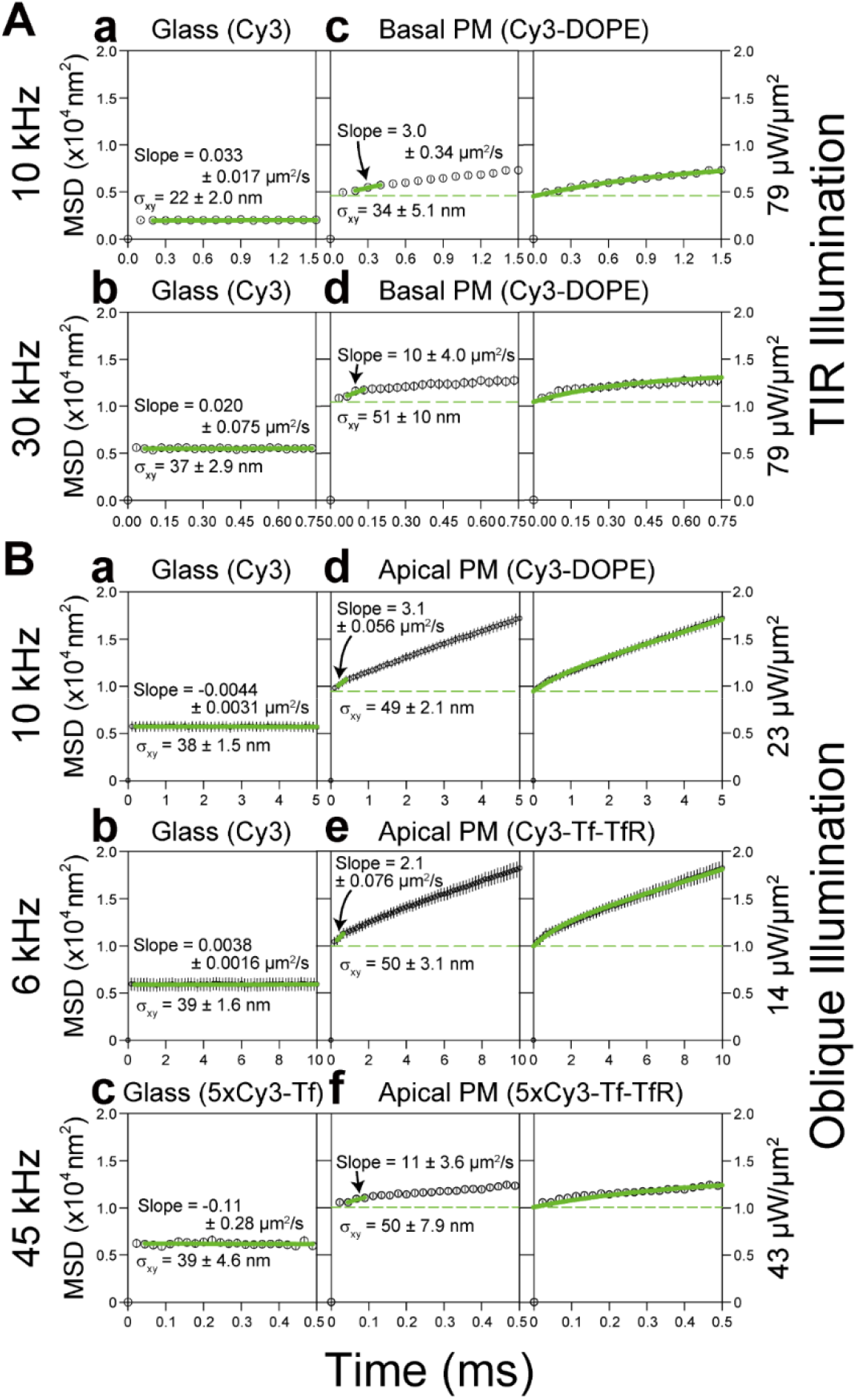
MSD-Δ*t* plots ensemble averaged over all trajectories, obtained by ultrahigh-speed single-molecule imaging of Cy3 molecules immobilized on coverslips or diffusing in the apical and basal PMs of T24 cells (using oblique-angle and TIR illuminations, respectively), providing estimates of single-molecule localization errors for diffusing molecules (as well as immobile molecules). All of the SEMs, including the error bars, are shown in the figure. The purposes of showing these figures are (1) to explain how to determine the single-molecule localization precisions of diffusing molecules in the PM using the MSD-Δt plot (because the method described in the caption to Fig. S2 is only useful for immobilized molecules) and (2) to show the actual localization precisions of diffusing molecules in the apical and basal PMs. First, see the **panels in the left column**, showing the MSD-Δ*t* plots for molecules immobilized on the glass. Experimental MSD-Δ*t* plots even for immobile molecules are expected to exhibit an offset, due to the position determination error; i.e., the flat MSD-Δ*t* plot with a constant value (against Δ*t*), which equals 4*σ*_xy_^2^ (where *σ*_xy_ = [*σ*_x_ + *σ*_y_]/2) (Dietrich et al., 2002; Martin et al., 2002). The linear fitting indeed showed that the slopes were ∼0; the localization precisions determined here for Cy3 on the glass at 10 and 30 kHz were 22 and 37 nm, respectively (TIR illumination at 79 µW/µm^2^). These results are consistent with those for immobilized molecules determined by the first method described in Fig. S2 (based on the standard deviations of the *x* and *y* position determinations for 15 consecutive frames) and shown in Fig. S3, B **and** C (20 and 34 nm, respectively; all SEMs for these values are given in the figure; also see Fig. 1 F and Table 1). Next, see **the panels in the middle and right columns**, showing the MSD-Δ*t* plots for molecules undergoing diffusion in the PM. As shown in the **panels in the left column** (immobilized molecules), the MSD values almost plateaued (which is the offset value) by the second step (Δ*t* =0.2 and 0.066 ms for Cy3 at 10 and 30 kHz, respectively). This means that the offset value of the MSD-Δ*t* plot for diffusing molecules in the PM can be estimated as the y-intercept (by extrapolation) of the linear-fit function for the second, third, and fourth steps in the MSD-Δ*t* plot (Fujiwara et al., 2002) (**middle column**; see green keys), which is 4*σ*_xy_^2^. Therefore, the MSD-Δ*t* plot, representing the diffusion effects, is in fact given by the plot of MSD - 4*σ*_xy_^2^ against Δ*t*, as shown in Fig. 4 A. The single-molecule localization precisions determined this way (**middle column**) for Cy3-DOPE in the basal PM at 10 and 30 kHz were 34 and 51 nm, respectively, which were inferior to those found for the same molecules fixed on the glass (22 and 37 nm, respectively; **left column**). For Cy3-DOPE in the apical PM at 10 kHz, the localization precision was 49 nm, which was much worse than that in the basal PM (34 nm). The single-molecule localization precisions in the basal PM were worse than those determined on the glass, probably due to the higher background caused by cellular autofluorescence and the diffusional blurring of single-molecule spots. (**A**) The *TIR laser illumination*results of the ensemble-averaged MSD-Δ*t* plots, for single Cy3 molecules covalently linked to the cover-glass surface coated with 3-aminopropyltriethoxysilane (**left column**) and for single Cy3-DOPE incorporated in the *basal* PM (**middle and right columns**), using the highest laser intensities of TIR illumination available for this instrument at 532 nm (79 µW/µm^2^). This provided the *best single-molecule localization precisions* for recordings of Cy3 at 10 and 30 kHz, which were 22 and 37 nm on the glass (**left column, a and b**; *n* = 40 and 50, respectively), and 34 and 51 nm on the *basal* PM (**middle column, c and d**; *n* = 50 and 50, respectively). In the panels in the **right column**, the green curves are the best-fit functions describing the MSD-Δ*t* plots for the confined-diffusion model, in which molecules undergo free diffusion while totally confined within a limited area during the observation period (Eqs. 11-13 in Kusumi et al., 1993). Since the observation durations for single molecules (1.5 and 0.75 ms full x-axis scales) were shorter as compared with the dwell time of Cy3-DOPE within a compartment, the confined fitting, rather than the hop fitting, was employed. (**B**) The *oblique-angle laser illumination*results of the ensemble-averaged MSD-Δ*t* plots, for single Cy3 molecules and 5xCy3-Tf on the coverslip (**left column**) and for single Cy3-DOPE, Cy3-Tf (with a dye-to-protein molar ratio of 0.2, so that virtually all of the single Tf molecules are labeled with either 0 or 1 Cy3 molecule) bound to TfR, and 5xCy3-Tf bound to TfR in the *apical* PM (**middle and right columns**). The oblique-angle laser illumination is widely applicable and useful because it can illuminate molecules located deeper in the cytoplasm as well as those present in the apical PM. Therefore, it is extensively used in the present research (standard conditions using an oblique-angle laser illumination power density of 23 µW/µm^2^). The numbers of examined spots: **a** and **b**, *n* = 17 and 50; **c**, *n* = 40 and 150 (glass and apical PM, respectively). The illumination laser power densities were selected so that they were just beneath the level where dye saturation is obvious, and the single-molecule localization errors for various Cy3 specimens observed at different frame rates were similar to each other (see Fig. S3, B **and** C). More specifically, Cy3 and Cy3-DOPE at 10 kHz and 23 µW/µm^2^ (**a, d**), Cy3 and Cy3-Tf at 6 kHz and 14 (23 x [6/10]) µW/µm^2^ (**b, e**) (because the frame time is [10/6]-times longer at 6 kHz), and 5xCy3-Tf at 45 kHz at 43 µW/µm^2^ (**c, f**). **Panels in the left column (molecules on the glass)**. The localization precisions determined here for Cy3 at 10 kHz (38 nm at 23 µW/µm^2^) and for 5xCy3-Tf at 45 kHz (39 nm at 43 µW/µm^2^) were consistent with those found in Fig. S3 B (37 and 38 nm, respectively). **Panels in the middle and right columns (molecules in/on the *apical* PM).** The single-molecule localization precisions determined for Cy3-DOPE (10 kHz, 23 µW/µm^2^), Cy3-Tf (6 kHz, 14 µW/µm^2^), and 5xCy3-Tf (45 kHz, 43 µW/µm^2^) in/on the *apical* PM were 49, 50, and 50 nm, respectively, which were inferior to those found for the same molecules fixed on the glass (38, 39, and 39 nm, respectively; **left column**). The localization errors were greater in the apical PM, probably due to the higher background caused by cellular autofluorescence and the blurring of single-molecule spots due to molecular diffusion within a frame time. In the **right column** in panels **d** and **e**, the green curves are the best-fit functions describing the MSD-Δ*t* plots for an idealized hop-diffusion model (hop-diffusion fitting; see the subsection “Hop-diffusion fitting: the function describing the MSD-Δ*t* plot for particles undergoing hop diffusion” in the caption to Fig. S5). In panel **f**, since the observation duration for single 5xCy3-Tf molecules employed here (0.5 ms full x-axis scale) was shorter as compared with the dwell time of TfR within a compartment, the confined fitting, rather than hop fitting, was employed.

**Figure S5.**
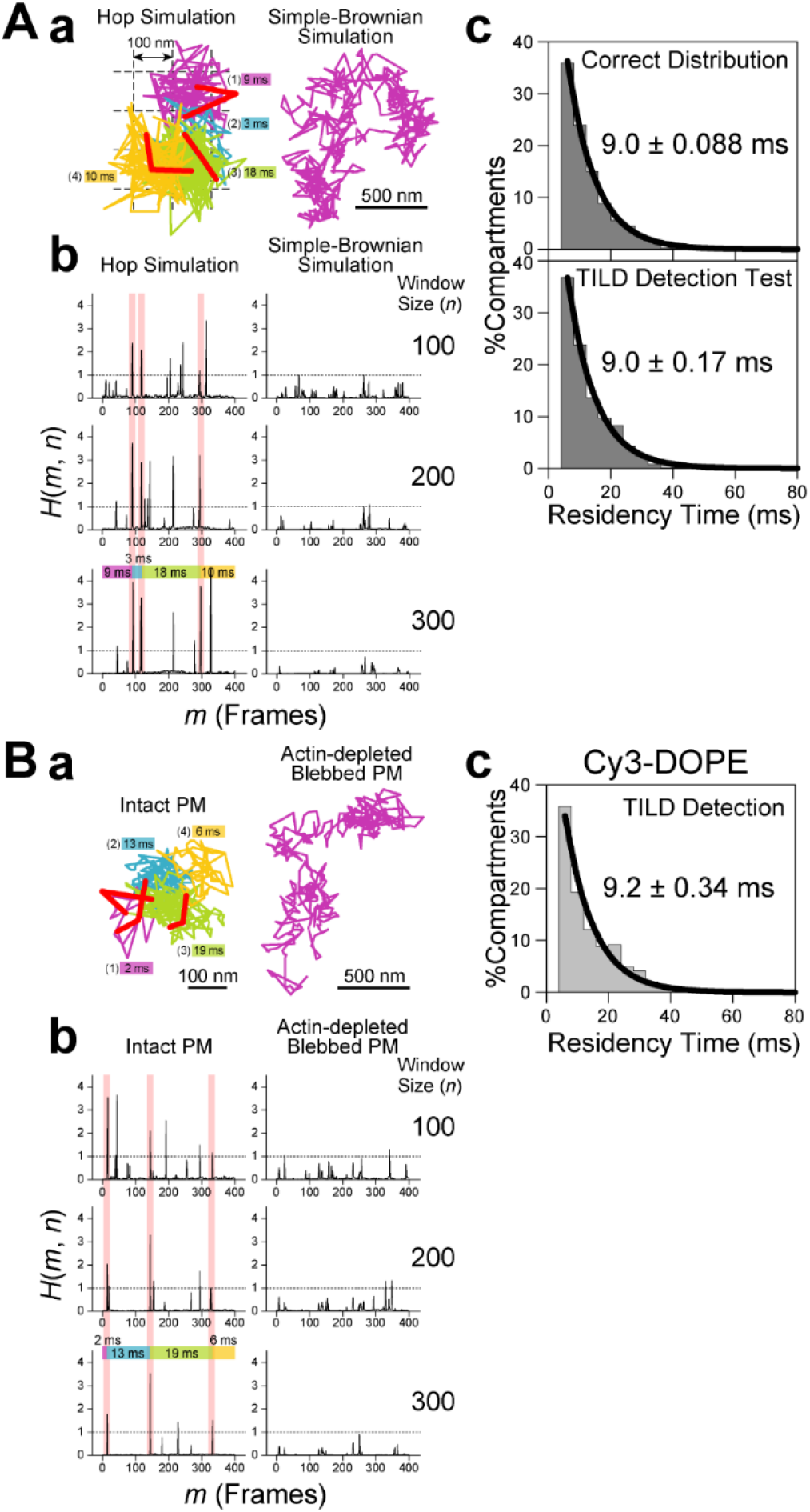
The TILD method for detecting the moment (instance) when a diffusing molecule in the PM undergoes the hop movement from a compartment to an adjacent one in the PM, the theory for describing the distribution of the dwell lifetimes of molecules (within a compartment) undergoing hop diffusion, and the function for hop-diffusion fitting. We developed an improved method for detecting the hop moment (instance). This method detects the <u>T</u>ransient <u>I</u>ncrease of the effective <u>L</u>ocal <u>D</u>iffusion coefficient (TILD) in a single-molecule trajectory. TILDs are likely to occur when a molecule hops between two membrane compartments, but the analysis itself remains model-independent. When a molecule undergoes an intercompartmental hop, it experiences two compartments rather than one during the time window including the hop moment, thus increasing the effective local diffusion coefficient. In a given trajectory, consider a window with a size of *n* frames (*n* steps), starting from a frame *m* (*m* ∼ *m*+ *n*). Within this window, the center of the *n*recorded coordinates (*n* points) is defined, the radial displacements for all of these *n* points from this center are calculated, and then their maximal value *R*_MAX_(*m*, *n*) is determined. Next, the relative diffusion coefficient for this window is defined as

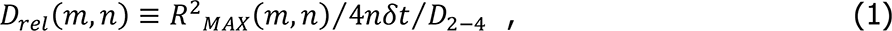

where *δt* is the time step in the trajectory (inverse of the frame rate), and *D*_2-4_ is the average short-term diffusion coefficient determined for the time interval between 2*δt* and 4*δt* (averaged over the whole trajectory) (Kusumi et al., 1993), which is included for normalization. For free Brownian diffusion, *D*_rel_(*m*, *n*) is ∼1 (allowing for statistical variations), independent of *m* or *n*. If a molecule is temporarily trapped in a finite domain, then as the window size *n* increases, *D*_rel_(*m*, *n*) decreases due to entrapment. When the window size has increased sufficiently to include the release of the diffusing particle from the finite domain, there will be a sharp increase in *D*_rel_(*m*, *n*), due to the extended range of diffusion. To flag releases, the function

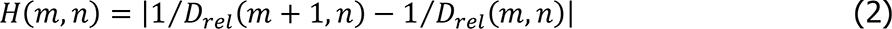

was employed, and by scanning all possible *m* and *n* pairs in the full trajectory, a map of *H*(*m*, *n*) was produced. Releases from the trapped domain are flagged by sharp peaks in *H*(*m*, *n*) for both the special starting position (e.g., if position *m* is before a hop and position *m*+ 1 is after a hop, then for all window sizes *n*, *D*_rel_(*m*, *n*) will be greater than *D*_rel_(*m* + 1, *n*)) and for the combination of a starting position and a window size (e.g., if the trajectory, starting from position *m* with a window size *n*, and ending at point *p* = *m* + *n*, is wholly within an entrapped domain, and if extending the window size by 1 includes release from the entrapped domain, then *D*_rel_(*m*, *n* + 1) will be greater than *D*_rel_(*m*, *n*) for all *m* and *n* values, such that *m*+ *n* = *p*). The minimum number of points within an entrapped domain required to allow the detection of the moments of TILDs is determined by the stochastic nature of diffusion (+ the diffusion coefficient within the domain and the domain size) and the noise present in the position determination. As such, due caution was exercised to avoid choosing a plethora of small compartments erroneously. Typically, the total length of each trajectory analyzed here was 1,000 steps. The *n* value was varied from 21 to *N*-*m*, where *N* is the total number of steps in a trajectory (= 1,000). In addition to this anterograde direction of analysis over a single trajectory, the same trajectory was analyzed in the retrograde direction in the same way as for the anterograde direction, in order to determine whether the sudden increase in *H*(*m*, *n*) can also be observed for the same *m* in the anterograde analysis. Therefore, for a single *m* value, *H*(*m*, *n*) was calculated for [*N* - 40] windows ([*N* - *m* - 20] + [*m* - 20]). Here, we first describe the results of testing the TILD method using Monte-Carlo simulated hop and simple-Brownian trajectories with a single-molecule localization precision of 50 nm (error for Cy3 in the apical PM, which is the worst precision in the present report) (**A**). Second, we describe the application of the TILD method to detect the hop moments in single-molecule trajectories of Cy3-DOPE, observed in the intact apical PM as well as in the actin-depleted blebbed PM (**B**). (**A**) **Testing the TILD detection method using computer-generated hop and simple-Brownian diffusion trajectories, and examination of the exponential distributions of the residency times within a compartment.** (**A-a**) Typical hop-diffusion (**left**) and simple-Brownian (**right**) trajectories generated by Monte Carlo simulation (every 0.1 ms; the initial 400-steps of the 1,000-step-long trajectories are shown here, whereas the TILD analysis was performed for the full 1,000 steps for all of the trajectories). The moments of TILDs determined by the developed protocol are shown by the thick, short red subtrajectories (three-frame trajectories defined by the TILD moment ± 1 frame; note their tilde-like shapes). This particular hop-diffusion trajectory (400-frames long) on the **left** contained 3 TILDs. The simple-Brownian trajectory (400-frames long) on the **right** exhibits no TILD. Hop diffusion trajectories were generated as described previously (Ritchie et al., 2005), except that the unit time step was 0.1 µs. Using a two-dimensional square array of partially permeable barriers separated by 100 nm (*L*), with a probability of transmission per attempt of 0.0005, the experimental *D*_MACRO_ for Cy3-DOPE in the intact apical PM (0.30 µm^2^/s; Table 2) was reproduced (*D*_micro_ was set at 9 µm^2^/s, as shown in Fujiwara et al., 2002). A single-molecule localization error of 0, 25, or 50 nm was added as the Gaussian noise to each x- and y-coordinate in the hop trajectories (here, the trajectories including a 50-nm localization error are shown). Among 100 trajectories generated by the simulation (1,000 frames per trajectory), 97, 94, and 80 trajectories were statistically classified into the suppressed diffusion mode in the presence of single-molecule localization errors of 0, 25, and 50 nm, respectively. Test trajectories of simple-Brownian particles were generated by Monte Carlo simulations, using a diffusion coefficient of 6 µm^2^/s, as experimentally obtained in blebbed PMs (Table 2; employing 9 µm^2^/s made virtually no difference), and Gaussian localization errors were added. (**A-b**) Typical plots of *H*(*m*, *n*) vs. *m* (only the results with window sizes of *n* = 100, 200, and 300 for the trajectories shown in **A-a**. The sharp changes (peaks) in *H*(*m*, *n*) are likely to represent the hop movements (transient increase of local diffusion coefficient), and spurious peaks from statistical variations and noise can mostly be distinguished because they only appear in the displays for limited numbers of windows. Briefly, for the same frame number *m*, *H*(*m*, *n*) was calculated for all possible *n*’s (*N*-40 windows), and when the percentage of windows in which *H*(*m*, *n*) ≥ 1 was greater than 20% among a total of [*N* - 40] windows, the molecule was regarded as undergoing the process of intercompartmental hopping. In this figure, the peaks that satisfied these thresholds are highlighted by vertical pink bars, indicating the occurrences of TILDs, i.e., hop events. Other peaks in this display did not satisfy the threshold conditions explained here, and do not represent TILDs. In the application of the TILD detection method to individual trajectories, 3-frame running averaging (replacing the position of the *k*th frame with the position averaged for the *k* - 1, *k*, and *k* + 1 frames) was first applied to each trajectory, to minimize the effect of apparently large displacements that stochastically occurred due to the 50-nm single-molecule localization error. The detectability percentages of hop events (given by the simulation program) for simulated trajectories classified into suppressed diffusion were 82, 76, and 66%; and the accuracies of the predicted hop events were 75, 66, and 60% at localization errors of 0, 25, and 50 nm, respectively. In our standard conditions for long-term single-molecule tracking experiments (1,000 steps; 0.1-ms resolution; a 561-nm laser excitation laser power of 23 µW/µm^2^ at the sample), the single-molecule localization error was 49 nm in the apical PM. (**A-c**) Distributions of the residency lifetimes within a compartment for Monte-Carlo simulated test particles undergoing hop diffusion. The residency time of a particle within a compartment was obtained as the duration between two consecutive TILDs. **Top** (**Ground Truth Distribution Given by the Simulation**) The correct distribution of the residency times determined from the hop events (given by the simulation program), which occurred in 100 simulated 1,000-frame long hop trajectories. The residency times shorter than 40 frames (4 ms) were neglected, due to various uncertainties in the short time ranges. The distribution could be fitted with a single exponential function with a decay time constant of 9.0 ± 0.088 ms (mean ± SEM; SEM is provided as the fitting error of the 68.3% confidence interval). The exponential distribution of the residency times found for simulated hop-diffusion trajectories can actually be predicted theoretically, as summarized in the subsection “**Expected distribution of the residency times: development of the hop diffusion theory**” at the end of this caption. The theory also predicts that its decay time constant can be described by *L*^2^/4*D*_MACRO_. In the present simulation, the average dwell time calculated using this equation was 8.3 ms (*L* = 100 nm, the average *D*_MACRO_ obtained from the simulation was 0.3 µm^2^/s). This value agrees quite well with the dwell lifetimes obtained by simulated hop diffusion trajectories (9.0 ± 0.088 ms). **Bottom** (**TILD Detection Test**) The residency time distribution, determined by the TILD-detection method from 100 simulated 1,000-frame long hop trajectories that included a single-molecule localization precision of 50 nm (residencies in 763 compartments with durations longer than 4 ms). The decay time constant was 9.0 ± 0.17 ms. This agrees well with the correct distribution, suggesting that the developed protocol is useful for evaluating the residency lifetime, although at the level of individual hops, our software misses hops (66% detectability) and incorrectly detects hops (60% accuracy). The number of detected TILDs per 1,000-frame simulated *simple-Brownian trajectories*, using a diffusion coefficient of 6 µm^2^/s and a localization precision of 50 nm, was only 0.35/trajectory (*n* = 100 trajectories; 0.33 when a diffusion coefficient of 9 µm^2^/s was assumed). (**B**) **Detecting TILDs in Cy3-DOPE and TfR trajectories obtained in intact and actin-depleted blebbed PMs.** (**B-a**) Typical Cy3-DOPE trajectories (0.1-ms resolution) in the intact apical PM (**left**, the 40-ms-long initial part of the trajectory shown in Fig. 3 C) and in the actin-depleted blebbed apical PM (**right**) (typical among 50 and 20 trajectories, respectively). The moments of TILDs, detected as shown in **B-b**, are shown by the thick, red 3-step subtrajectories. The trajectory obtained in the intact apical PM contained three TILDs, whereas that in the actin-depleted blebbed apical PM exhibited no TILDs. In the trajectory obtained in the intact apical PM (**left**), the colors of the trajectories are changed across the short red TILD trajectory, and this convention was used throughout this report. (**B-b**) The plots of *H*(*m*, *n*) vs. *m* for the trajectories shown in **B-a**. TILDs were detected in all of the experimental trajectories of Cy3-DOPE and TfR obtained in the intact apical PM (see Fig. 3 D; 50 trajectories with a length of 1,000 frames for Cy3-DOPE at a frame rate of 10 kHz and for TfR at a frame rate of 6 kHz). The average numbers of detected TILDs (with intervals longer than 2 ms) per 1,000-frame long trajectory classified into the suppressed-diffusion mode were 7.8 (= 297 events/38 trajectories) for Cy3-DOPE and 4.4 (= 175 events/40 trajectories) for TfR. Meanwhile, in the actin-depleted blebbed PM, where more than 90% of the trajectories were statistically classified into the simple-Brownian diffusion mode (20 trajectories were examined for both Cy3-DOPE and TfR; Table 2), the numbers of detected TILDs per 1,000-frame long trajectory classified into the simple-Brownian diffusion mode were only 0.22 (= 4 events/18 trajectories) for Cy3-DOPE and 0.68 (= 13 events/19 trajectories) for TfR. (**B-c**) Distributions of the dwell times within a compartment for Cy3-DOPE (50 trajectories; 337 residencies) in the intact apical PM, with the best-fit exponential curves and dwell lifetimes of 9.2 ± 0.34 ms. The exponential shape of this distribution is consistent with the hop diffusion model (under strong-type confinements; see the subsection below). Furthermore, the exponential residency lifetime found for Cy3-DOPE (9.2 ms) agrees well with that found for the simulated hop-diffusion trajectories, using parameters similar to the experimentally determined values for Cy3-DOPE (9.0 ms; **A-c**, **top**).

## Expected distribution of the residency times: development of the hop diffusion theory

In the hop-diffusion literature, the average residency time within a compartment has been intuitively approximated as *L*^2^⁄4*D*_MACRO_. However, this expression has never been proven rigorously. In addition, the distribution of the residency times has remained unknown even for an idealized hop diffusion model, in which molecules undergo free diffusion (in viscous media) in the presence of equally spaced, equi-potential semi-permeable diffusion barriers. Here, we demonstrate that the residency time distribution is given by the sum of exponential distributions, which can be approximated well by a single exponential function with a decay constant of *L*^2^⁄4*D*_MACRO_, if the confinement effect is strong.

Consider a Brownian particle in a square region with semi-permeable boundaries, and calculate the distribution of first exit times. The residency time distribution is analogous to the distribution of the first time a particle exits a certain region. We assume that diffusion along the *x* and *y* directions is independent, which allows us to solve the problem for a one-dimensional (1D) system, and then generalize the results to the two-dimensional (2D) case.

Consider a Brownian particle in 1D, initially placed at *x* = *x*_o_ between two boundaries located at *x* = 0 and *x* = *L*, that can diffuse freely in this bounded region. When the particle attempts to leave this region, it faces resistance, and thus the boundaries can be considered as partially permeable barriers. Once the particle crosses the boundary, it can never go back. Therefore, the time when it leaves the region is always the first exit time. The probability distribution of such a particle is governed by the diffusion equation,

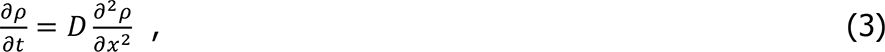

with the boundary conditions,

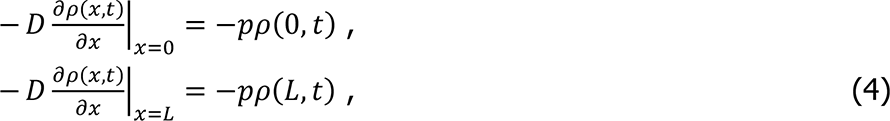

where *D* is the microscopic diffusion coefficient within a domain, and *p* is a constant related to the permeability of the boundaries, such that *p* → 0 and *p* → ∞ correspond to impenetrable and completely permeable boundaries, respectively. The solution can easily be obtained by using separation of variables (Carslaw and Jaeger, 1986), and is given in the following form

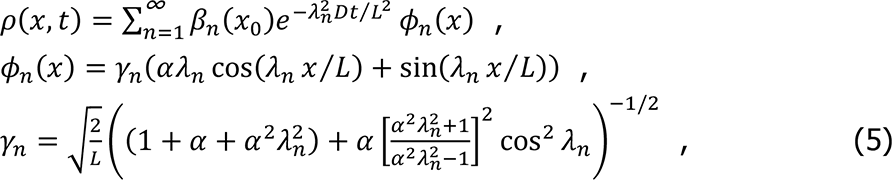

where *β*_n_(*x*_o_)’s are determined by the initial conditions, *α* = *D*⁄*pL*, and *λ*_n_’s, *n* = 1, 2, 3, …, are the positive solutions of

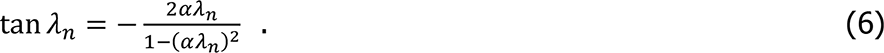

If the particle is initially at *x* = *x*_o_ such that *ρ*(*x*, *t*) = *δ*(*x* - *x*_o_), then the coefficient *β*_n_ becomes

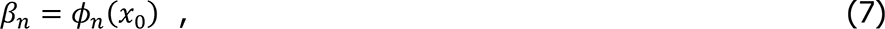

so that

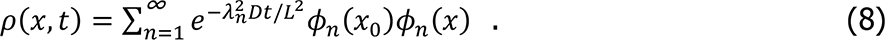

The cumulative probability distribution can be expressed as

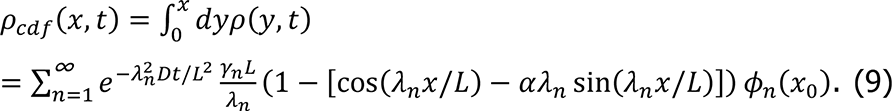

Finally, the distribution of exit times *f*_ex_(*t*), and its cumulative *F*_ex_(*t*), can be calculated from the cumulative probability distribution above (Redner, 2001)

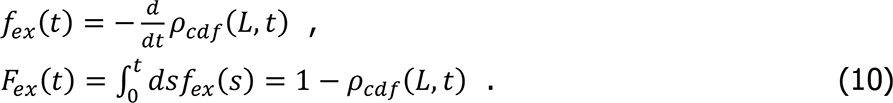

Now we need to generalize this result to two dimensions, and obtain the distribution of exit times from a square compartment of area *L*^2^. This was readily achieved by realizing that the time of the first exit is the minimum of the first exit time in the *x* and *y* directions. Using the well-known result for the distribution of the minimum of two independent random variables (Gumbel, 2004), we obtain the cumulative probability distribution for the exit time distribution as

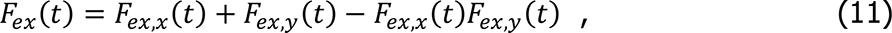

where *F*_ex,w_(*t*) stands for the probability that the particle exits through one of the boundaries along the *w* direction until time *t*, and is explicitly given by

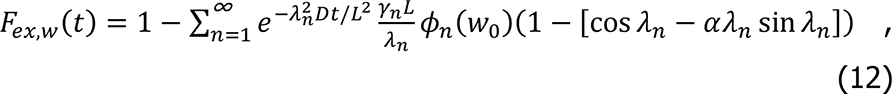

where *w*_o_ is the initial position along the *w* axis. Usually, the initial conditions are not accessible, so it is reasonable to average all of the different initial states. By averaging *F*_ex,w_(*t*) over all initial positions *w*_o_ between 0 and *L* with equal weight, we obtain

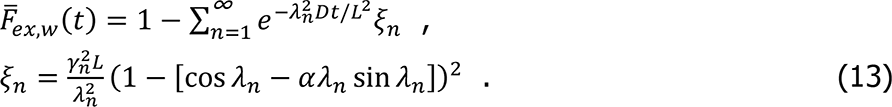

Using Eq. 13, Eq. 11 further simplifies to

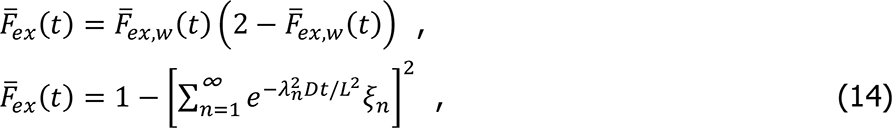

as there is no difference in the statistics of the position between the *x* and *y* directions after averaging over the initial position. The probability distribution for the exit times, or the residency time distribution, is given by the first derivative of Eq. 14 with respect to time

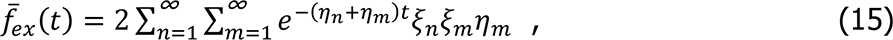

Where 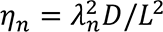.

The contributions of the higher order terms in Eq. 15 are quickly minimized, due to the exponential term. In many cases of practical interest, the first term with *n* = 1, *m*= 1 alone could be a good approximation of the result. To assess the validity of this argument, let us consider the ratio of the exponential factors in the first two terms of the double summation in Eq. 15. This ratio is equal to

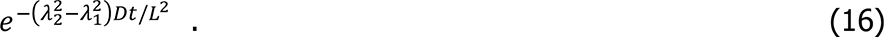

Inspecting the equation for *λ*_n_’s, Eq. 6, we notice that *λ*_n+1_ ≈ *λ*_n_ + *π*, such that

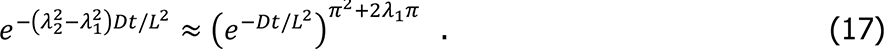

Therefore, the higher order terms decay with almost the 10th or greater powers of *e*^-Dt/L2^, which are always less than 1 for *t* > 0. Similarly, it can be shown that the coefficient of the exponential, *ξ*_n_*ξ*_m_*η*_m_, also decreases as *n* and *m* increase.

When the confinement effect is strong, the Brownian particle spends a sufficient amount of time in each compartment to cover it uniformly before escaping.

Mathematically, this corresponds to the limit

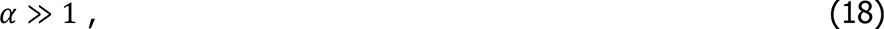

in which *ξ*_n_ is approximately equal to 1. Thus, the residency time distribution given in Eq. 15 can be approximated by a single exponential

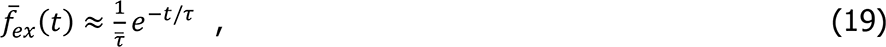

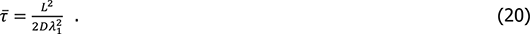

As the confinement effect becomes stronger, we would expect the average residency time to diverge. Therefore, in the strong confinement limit, *λ*_1_ would be close to 0 such that tan *λ*_1_ ≈ *λ*_1_, and Eq. 6 can be replaced by an approximate form

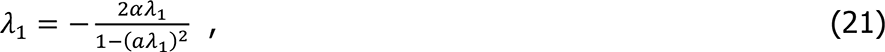

which gives

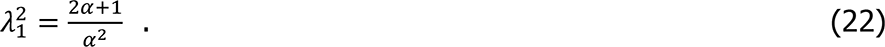

Therefore, the average residency time is equal to

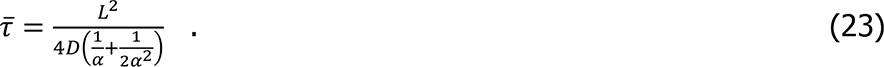

We now need to express *α* in terms of a quantity that can be experimentally measured. In the strong confinement limit that we are interested in, the Brownian particle behaves much like a random walker in a 2D lattice with lattice spacing *L*, for durations longer than the average residency time. In this picture, each lattice site corresponds to a square compartment of area *L*^2^, and the random walker takes steps between adjacent lattice sites at a rate 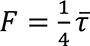. As the random walker can move in 4 directions in 2D, the rate of escape is 4*F*, such that the average residency time is *τ̅*. This description is appropriate as long as the random walker explores most of each compartment before it leaves. This ensures that the probability distribution within each compartment quickly becomes uniform and that the escape time distribution can be well approximated by a single exponential. The probability of finding the random walker at a lattice site (*m*, *n*) is governed by a Master equation (Hughes, 1995)

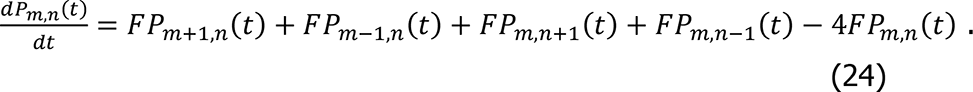

With a straightforward calculation, the mean square displacement of a random walker that is initially at the origin is given by

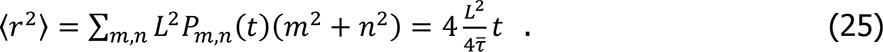

As this description is appropriate for times longer than *τ̅*, the diffusion coefficient associated with this random walk is the macroscopic diffusion coefficient,

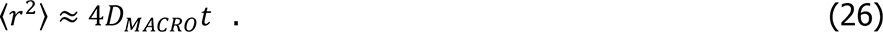

Combining Eqs. 23, 25 and 26, we obtain

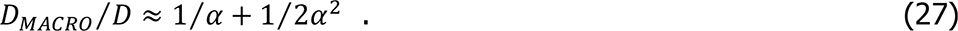

From many published reports (see the Table 1 summary in Kusumi et al., 2005), generally

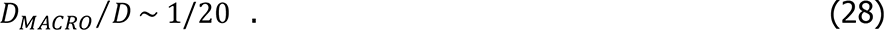

By solving Eq. 27 using this value,

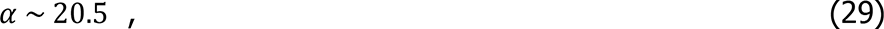

which satisfies the condition for strong confinement (Eq. 18, i.e. ≫ 1). Accordingly, under the strong confinement conditions, the second order term proportional to *α*^-2^ in Eq. 31 can be neglected, and Eq. 23 can simply be expressed as

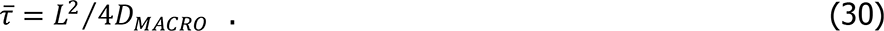

Therefore, the decay time constant of the exponential distribution of the residency time within a compartment, given by Eq. 23, can be simplified to the expression given by Eq. 30.

We obtained the distribution of the dwell lifetimes in a compartment by directly measuring the dwell time using the TILD analysis (**A-c** and **B-c**). In the real PM, the dwell time of diffusing molecules within a compartment would be affected by the variations in the compartment sizes and shapes and in the properties of the compartment boundaries, due to the differences in actin binding proteins and TM picket proteins. However, as shown in **B-c** and Fig. 5 A, the residency time distribution in the PM is well approximated by an exponential function, probably because these variations both lengthen and shorten the dwell lifetimes quite randomly.

## Hop-diffusion fitting: the function describing the MSD-Δ*t* plot for particles undergoing hop diffusion

We developed an equation for the MSD-Δ*t* plot for particles undergoing idealized hop diffusion; i.e., free diffusion in the presence of equally spaced, equi-potential semi-permeable diffusion barriers. Such hop diffusion can be characterized by the following three parameters: the compartment size (the distance between barriers), *L*, the microscopic diffusion coefficient within a compartment (true diffusion coefficient in the absence of the compartments), *D*_micro_, and the long-term diffusion coefficient over many compartments, *D*_MACRO_. One of the key parameters for hop diffusion, the residency time within a compartment, can be calculated from these parameters as *L*^2^⁄4*D*_MACRO_, as shown in the previous subsection.

By employing the notations *γ* = *D*_MACRO_⁄(*D*_micro_ - *D*_MACRO_) and *τ* = 4*D*_micro_*t*⁄*L*^2^, the 1-dimensional MSD in the real time domain, which is the inverse Laplace transform of Eq. 18 in (Kenkre et al., 2008) for a 1-dimensional MSD averaged over all initial locations, can be obtained as

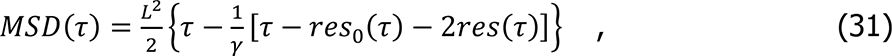

where the terms *res*_o_(*τ*) and *res*(*τ*) are the sum of the residues that arise from the inverse Laplace transform. The zeroth residue, *res*_o_(*τ*), is expressed as

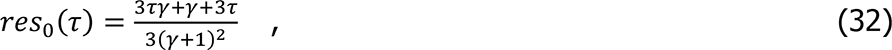

whereas the term *res*(*τ*), representing half the sum of all other residues, is expressed as

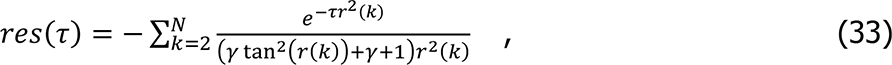

where *r*(*k*) (*k* = 2, 3, 4, . ..) is the *k*th root of 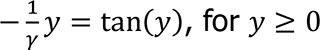. As performed in (Kenkre et al., 2008), we employed the bisection method to find the roots numerically with high precisions and obtained the accurate result for a sufficiently large *N*. Namely, by rearranging terms in Eq. 33, we obtain the following equation.

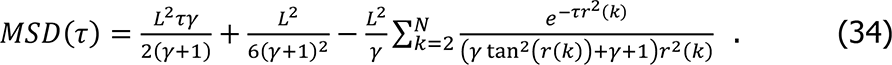

We used this equation for fitting the experimental MSD values. As for the physical meaning, the first term characterizes the long term behavior of MSD, which is approximately equal to 2*D*_MACRO_*t* for strong confinement (γ ∼ 0) and 2*D*_micro_*t* for weak confinement (γ → ∞). The second and third terms describe the transient behavior of the MSD. At around *τ* ∼ 0, the second and third terms cancel each other out. In the longer time limit, the third term becomes negligible, and the second term becomes equal to the intercept of the linear MSD.

## Captions for Videos 1 to 4

### Video 1

Single Cy3 molecules covalently linked to the glass surface, observed at 10 kHz (every 0.1 ms) (observation period of 1,500 frames = 150 ms; Replay, 167x-slowed from real time) using oblique-angle illumination at 23 µW/µm^2^. a, b, and c represent the fluorescent spots corresponding to those in Fig. 2, A **and** B. The on-periods including short off-periods of up to 3 frames are highlighted by the orange circle, and the off-periods are highlighted by the cyan circle.

### Video 2

Single Cy3-DOPE molecules diffusing in the intact apical PM, observed at 10 kHz (every 0.1 ms) (observation period of 750 frames = 75 ms; Replay, 167x-slowed from real time) using oblique-angle illumination at 23 µW/µm^2^. Most fluorescent spots were photobleached within 100 frames after starting the illumination, but a small percentage of the spots lasted longer than 300 frames. The long-lasting spot enclosed within the yellow square in the movie on the left is enlarged and highlighted by the circle in the movie on the right.

### Video 3

Single Cy3-DOPE molecules diffusing in the intact apical PM and the actin-depleted blebbed PM (left and right respectively), observed at 10 kHz (every 0.1 ms) (see Fig. 3, C **and** D) (observation periods of 750 and 300 frames = 75 and 30 ms, respectively; Replay, 167x-slowed from real time). The statistical analysis method developed in this study classified the trajectories into the suppressed-diffusion and simple-Brownian diffusion modes, respectively. The flickering signals seen in the movies, in addition to the long-lasting spots (indicated by the trajectories), are probably due to the fluctuation of the autofluorescence signal from the cytoplasm, because they are also found in the movies of Cy3-Tf (**Video 4**).

### Video 4

Single TfR molecules, bound by Cy3-Tf, diffusing in the intact apical PM or the actin-depleted blebbed PM (left and right respectively), observed at 6 kHz (every 0.167 ms) (observation periods of 600 and 180 frames = 100 and 30 ms, respectively; Replay, 167x-slowed from real time) (see Fig. 3 D). The statistical analysis method developed in this study classified the trajectories into the suppressed-diffusion and simple-Brownian diffusion modes, respectively.

